# A single-cell and spatial atlas of autopsy tissues reveals pathology and cellular targets of SARS-CoV-2

**DOI:** 10.1101/2021.02.25.430130

**Authors:** Toni M. Delorey, Carly G. K. Ziegler, Graham Heimberg, Rachelly Normand, Yiming Yang, Asa Segerstolpe, Domenic Abbondanza, Stephen J. Fleming, Ayshwarya Subramanian, Daniel T. Montoro, Karthik A. Jagadeesh, Kushal K. Dey, Pritha Sen, Michal Slyper, Yered H. Pita-Juárez, Devan Phillips, Zohar Bloom-Ackerman, Nick Barkas, Andrea Ganna, James Gomez, Erica Normandin, Pourya Naderi, Yury V. Popov, Siddharth S. Raju, Sebastian Niezen, Linus T.-Y. Tsai, Katherine J. Siddle, Malika Sud, Victoria M. Tran, Shamsudheen K. Vellarikkal, Liat Amir-Zilberstein, Deepak S. Atri, Joseph Beechem, Olga R. Brook, Jonathan Chen, Prajan Divakar, Phylicia Dorceus, Jesse M. Engreitz, Adam Essene, Donna M. Fitzgerald, Robin Fropf, Steven Gazal, Joshua Gould, John Grzyb, Tyler Harvey, Jonathan Hecht, Tyler Hether, Judit Jane-Valbuena, Michael Leney-Greene, Hui Ma, Cristin McCabe, Daniel E. McLoughlin, Eric M. Miller, Christoph Muus, Mari Niemi, Robert Padera, Liuliu Pan, Deepti Pant, Carmel Pe’er, Jenna Pfiffner-Borges, Christopher J. Pinto, Jacob Plaisted, Jason Reeves, Marty Ross, Melissa Rudy, Erroll H. Rueckert, Michelle Siciliano, Alexander Sturm, Ellen Todres, Avinash Waghray, Sarah Warren, Shuting Zhang, Daniel R. Zollinger, Lisa Cosimi, Rajat M. Gupta, Nir Hacohen, Winston Hide, Alkes L. Price, Jayaraj Rajagopal, Purushothama Rao Tata, Stefan Riedel, Gyongyi Szabo, Timothy L. Tickle, Deborah Hung, Pardis C. Sabeti, Richard Novak, Robert Rogers, Donald E. Ingber, Z. Gordon Jiang, Dejan Juric, Mehrtash Babadi, Samouil L. Farhi, James R. Stone, Ioannis S. Vlachos, Isaac H. Solomon, Orr Ashenberg, Caroline B.M. Porter, Bo Li, Alex K. Shalek, Alexandra-Chloé Villani, Orit Rozenblatt-Rosen, Aviv Regev

**Affiliations:** Klarman Cell Observatory, Broad Institute of MIT and Harvard, Cambridge, MA 02142, USA, USA; Broad Institute of MIT and Harvard, Cambridge, MA 02142, USA; Program in Health Sciences & Technology, Harvard Medical School & Massachusetts Institute of Technology, Boston, MA 02115, USA; Institute for Medical Engineering & Science, Massachusetts Institute of Technology, Cambridge, MA 02139, USA; Koch Institute for Integrative Cancer Research, Massachusetts Institute of Technology, Cambridge, MA 02139, USA; Ragon Institute of MGH, MIT, and Harvard, Cambridge, MA 02139, USA; Harvard Graduate Program in Biophysics, Harvard University, Cambridge, MA 02138, USA; Center for Immunology and Inflammatory Diseases, Department of Medicine, Massachusetts General Hospital, Boston, MA 02114, USA; Center for Cancer Research, Massachusetts General Hospital, Harvard Medical School, Boston, MA 02114, USA; Harvard Medical School, Boston, MA 02115, USA; Massachusetts Institute of Technology, Cambridge, MA 02139, USA; Data Sciences Platform, Broad Institute of MIT and Harvard, Cambridge, MA 02142; Precision Cardiology Laboratory, Broad Institute of MIT and Harvard, Cambridge, MA 02142, USA; Department of Epidemiology, Harvard School of Public Health; Division of Infectious Diseases, Department of Medicine, Massachusetts General Hospital, Boston, MA 02114, USA; Department of Medicine, Harvard Medical School, Boston, MA 02115, USA; Department of Pathology, Beth Israel Deaconess Medical Center, Boston, MA 02115, USA; Harvard Medical School Initiative for RNA Medicine, Boston, MA 02115, USA; Cancer Research Institute, Beth Israel Deaconess Medical Center, Boston, MA 02115, USA; Infectious Disease and Microbiome Program, Broad Institute of MIT and Harvard, Cambridge, MA 02142, USA; Institute for Molecular Medicine Finland, Helsinki, Finland; Analytical & Translational Genetics Unit, Massachusetts General Hospital, Harvard Medical School, Boston, MA 02115, USA; Department of Medicine, Beth Israel Deaconess Medical Center, MA 02115, USA; Division of Gastroenterology, Hepatology and Nutrition, Department of Medicine, Beth Israel Deaconess Medical Center, Boston, MA 02215, USA; Department of Systems Biology, Harvard Medical School, Boston, MA 02115, USA; FAS Center for Systems Biology, Department of Organismic and Evolutionary Biology, Harvard University, Cambridge, MA 02138, USA; Division of Endocrinology, Diabetes, and Metabolism, Beth Israel Deaconess Medical Center, Boston, MA 02115; Boston Nutrition and Obesity Research Center Functional Genomics and Bioinformatics Core Boston, MA 02115, USA; Department of Organismic and Evolutionary Biology, Harvard University, Cambridge, MA, USA; Divisions of Cardiovascular Medicine and Genetics, Brigham and Women’s Hospital, Harvard Medical School, Boston, MA 02115, USA; NanoString Technologies Inc., Seattle, WA 98109, USA; Department of Radiology, Beth Israel Deaconess Medical Center, Boston, MA 02215, USA; Department of Pathology, Massachusetts General Hospital, Harvard Medical School, Boston, MA 02115, USA; Department of Genetics and BASE Initiative, Stanford University School of Medicine; Massachusetts General Hospital Cancer Center, Department of Medicine, Massachusetts General Hospital, Boston, MA 02114, USA; Center for Genetic Epidemiology, Department of Preventive Medicine, Keck School of Medicine, University of Southern California, Los Angeles, CA, USA; Department of Pathology, Brigham and Women’s Hospital, Boston, MA 02115; John A. Paulson School of Engineering and Applied Sciences, Harvard University, Cambridge, MA 02138; Harvard-MIT Division of Health Sciences and Technology, Cambridge MA; Department of Pathology, Harvard Medical School, Boston, MA 02115, USA; Harvard Stem Cell Institute, Cambridge, MA, USA; Center for Regenerative Medicine, Massachusetts General Hospital, Boston, MA 02114, USA; Infectious Diseases Division, Department of Medicine, Brigham and Women’s Hospital, Boston, MA, USA; Department of Medicine, Massachusetts General Hospital, Harvard Medical School, Boston, MA 02114, USA; Duke University School of Medicine, Durham, NC; Department of Genetics, Harvard Medical School, Boston, MA 02115, USA; Department of Molecular Biology and Center for Computational and Integrative Biology, Massachusetts General Hospital, Boston, MA 02114, USA; Department of Immunology and Infectious Diseases, Harvard T.H. Chan School of Public Health, Harvard University, Boston, MA, USA; Howard Hughes Medical Institute, Chevy Chase, MD, USA; Massachusetts Consortium on Pathogen Readiness, Boston, MA, USA; Wyss Institute for Biologically Inspired Engineering, Harvard University; Massachusetts General Hospital, MA 02114, USA; Vascular Biology Program and Department of Surgery, Boston Children’s Hospital, Harvard Medical School, Boston, MA USA; Program in Computational & Systems Biology, Massachusetts Institute of Technology, Cambridge, MA 02139, USA; Program in Immunology, Harvard Medical School, Boston, MA 02115, USA; Department of Chemistry, Massachusetts Institute of Technology, Cambridge, MA 02139, USA; Genentech, 1 DNA Way, South San Francisco, CA, USA

## Abstract

The SARS-CoV-2 pandemic has caused over 1 million deaths globally, mostly due to acute lung injury and acute respiratory distress syndrome, or direct complications resulting in multiple-organ failures. Little is known about the host tissue immune and cellular responses associated with COVID-19 infection, symptoms, and lethality. To address this, we collected tissues from 11 organs during the clinical autopsy of 17 individuals who succumbed to COVID-19, resulting in a tissue bank of approximately 420 specimens. We generated comprehensive cellular maps capturing COVID-19 biology related to patients’ demise through single-cell and single-nucleus RNA-Seq of lung, kidney, liver and heart tissues, and further contextualized our findings through spatial RNA profiling of distinct lung regions. We developed a computational framework that incorporates removal of ambient RNA and automated cell type annotation to facilitate comparison with other healthy and diseased tissue atlases. In the lung, we uncovered significantly altered transcriptional programs within the epithelial, immune, and stromal compartments and cell intrinsic changes in multiple cell types relative to lung tissue from healthy controls. We observed evidence of: alveolar type 2 (AT2) differentiation replacing depleted alveolar type 1 (AT1) lung epithelial cells, as previously seen in fibrosis; a concomitant increase in myofibroblasts reflective of defective tissue repair; and, putative TP63^+^ intrapulmonary basal-like progenitor (IPBLP) cells, similar to cells identified in H1N1 influenza, that may serve as an emergency cellular reserve for severely damaged alveoli. Together, these findings suggest the activation and failure of multiple avenues for regeneration of the epithelium in these terminal lungs. SARS-CoV-2 RNA reads were enriched in lung mononuclear phagocytic cells and endothelial cells, and these cells expressed distinct host response transcriptional programs. We corroborated the compositional and transcriptional changes in lung tissue through spatial analysis of RNA profiles *in situ* and distinguished unique tissue host responses between regions with and without viral RNA, and in COVID-19 donor tissues relative to healthy lung. Finally, we analyzed genetic regions implicated in COVID-19 GWAS with transcriptomic data to implicate specific cell types and genes associated with disease severity. Overall, our COVID-19 cell atlas is a foundational dataset to better understand the biological impact of SARS-CoV-2 infection across the human body and empowers the identification of new therapeutic interventions and prevention strategies.

## Introduction

Severe acute respiratory syndrome coronavirus 2 (SARS-CoV-2) causes coronavirus disease 2019 (COVID-19), which has resulted in over 1 million deaths globally as of November 2020 (https://covid19.who.int/). The variable host immune response to infection can result in a range of clinical outcomes spanning from an asymptomatic state, to severe illness, organ failures, and death. The vast majority of deaths are due to acute lung injury and acute respiratory distress syndrome (ARDS), or direct complications thereof that can lead to multiple organ failure^1–4^. Progression to ARDS is thought to reflect: a combination of increasing viral load; cytopathic effects; translocation of virus into pulmonary tissue; and, inappropriate or insufficient host immune responses^1–6^. Collectively, this contributes to clinical deterioration in the acute phase of systemic illness, leading to ineffective viral clearance and collateral tissue damage—which can impact the lung^1–8^, gastrointestinal tract^5^, kidney^6, 7^, liver^3, 6^, vasculature^8^, heart^2, 6, 9, 10^ and brain^6, 11–14^—resulting in single or multi-organ failure^1–4^.

While clinical knowledge of severe COVID-19 is developing rapidly, our molecular and cellular understanding of disease pathogenesis and pathophysiology remains limited. Current data indicate that severe COVID-19 is accompanied by an inappropriate host immune response, characterized by cytokine storms involving pro-inflammatory cytokines (TNF-α, IL-1 and IL-6), chemokines (IL-8) and a diminished antiviral interferon response^15–18^. A central unanswered question is why some patients enter a second phase of disease that can lead to death, and what are the molecular characteristics of this condition. In particular, we do not yet know: (**1**) the cell types and states altered in COVID-19 affected tissues; (**2**) how these cellular compositions and programming differ from those observed in healthy tissues or other relevant diseases; (**3**) the cell types infected by the virus; (**4**) how viral infection alters local cellular responses; and, (**5**) how genetic loci associated with severe COVID-19 in Genome Wide Association Studies (GWAS) may drive disease. Addressing these questions using relevant human tissue sources will be essential to inform the identification of new therapeutic targets and prevention strategies.

It has been challenging to tackle these questions as it is rare to obtain samples from essential affected organs of living patients. Autopsies are thus a critical path to gaining knowledge of severe COVID-19 pathology^19–28^, but are usually followed by established molecular histopathology approaches relying on marker preselection, such as immunohistochemistry, which may limit discovery of novel molecular insights into COVID-19 pathogenesis. Conversely, comprehensive genomic studies remain challenging as autopsy specimens are often collected after a prolonged post-mortem interval (PMI), which can lead to degradation of biomolecules. Furthermore, the heterogeneous composition of, and unpredictable cellular responses within, post-mortem tissues require high resolution analyses that can distinguish individual cells and their features.

To overcome these challenges, we leveraged ongoing clinical COVID-19 autopsy efforts across three major Boston area hospitals to orchestrate a concerted study and assemble a comprehensive biobank of approximately 420 autopsy specimens spanning 11 organs from 17 donors who succumbed to COVID-19. We then utilized recently developed methods for single-nucleus RNA-Seq (snRNA-Seq) from frozen specimens^29, 30^ and for spatial RNA profiling from formalin fixed paraffin embedded (FFPE) tissues^31^, and developed and applied analytical strategies to overcome ambient RNA in autopsy specimens and relate annotations across datasets. Together, this enabled us to generate comprehensive single-cell/single-nucleus tissue atlases of lung, kidney, liver and heart tissues from COVID-19 donors, and to further analyze several distinct lung regions with spatial RNA profiling methods. Focusing on lung tissue, we charted cell composition and transcriptional programs associated with severe COVID-19 illness. We additionally resolved which cell subsets are enriched for SARS-CoV-2 RNA and host-virus dependencies, as well as corroborated select observations through spatial analyses. In parallel, we constructed and annotated tissue atlases for kidney, liver, and heart tissues—all sites with potential pathological involvement. Finally, we examined the potential cellular basis for COVID-19 associated genetic risk loci. Overall, our atlas provides critical insights into the pathology, pathogenesis and pathophysiology of severe COVID-19, and should help inform future therapeutic development and prophylactics.

## RESULTS

### A multi-organ autopsy cohort of COVID-19 and unique sample processing pipeline

We assembled a COVID-19 autopsy cohort of eleven male and six female deceased donors, spanning >30 to >89 years of age, diverse racial and ethnic backgrounds, and a range of intermittent mandatory ventilation (IMV; 0-24 days) periods and days from symptom onset to death (S/s to death) (Fig. 1a). Though different organ failures lead to a patient’s demise, lung involvement was reported in all donors (Fig. 1a, **Supplemental Table 1**). The autopsies were performed across three Boston area hospitals: Beth Israel Deaconess Medical Center (BIDMC), Brigham and Women’s hospital (BWH) and Massachusetts General Hospital (MGH) (**Methods**). The tissue types collected, as well as postmortem intervals (PMI, 1.4-24 hours), varied across hospital collection sites (Fig. 1a). From all donors across all sites, we systematically collected at least one portion of the lung, heart, and liver. When possible, we also collected one or more specimens of kidney, spleen, trachea, peribronchial/subcarinal lymph node, skeletal muscle, nasal scraping, and oral mucosa. Brain tissue was collected for the single donor who presented with neurological symptoms. Immediately after autopsy, we preserved tissue specimens via FFPE for future spatial analysis on-site, while a subset of specimens were transported on ice to the Broad Institute for further processing (**Methods**) (Fig. 1b).

**Figure 1.**
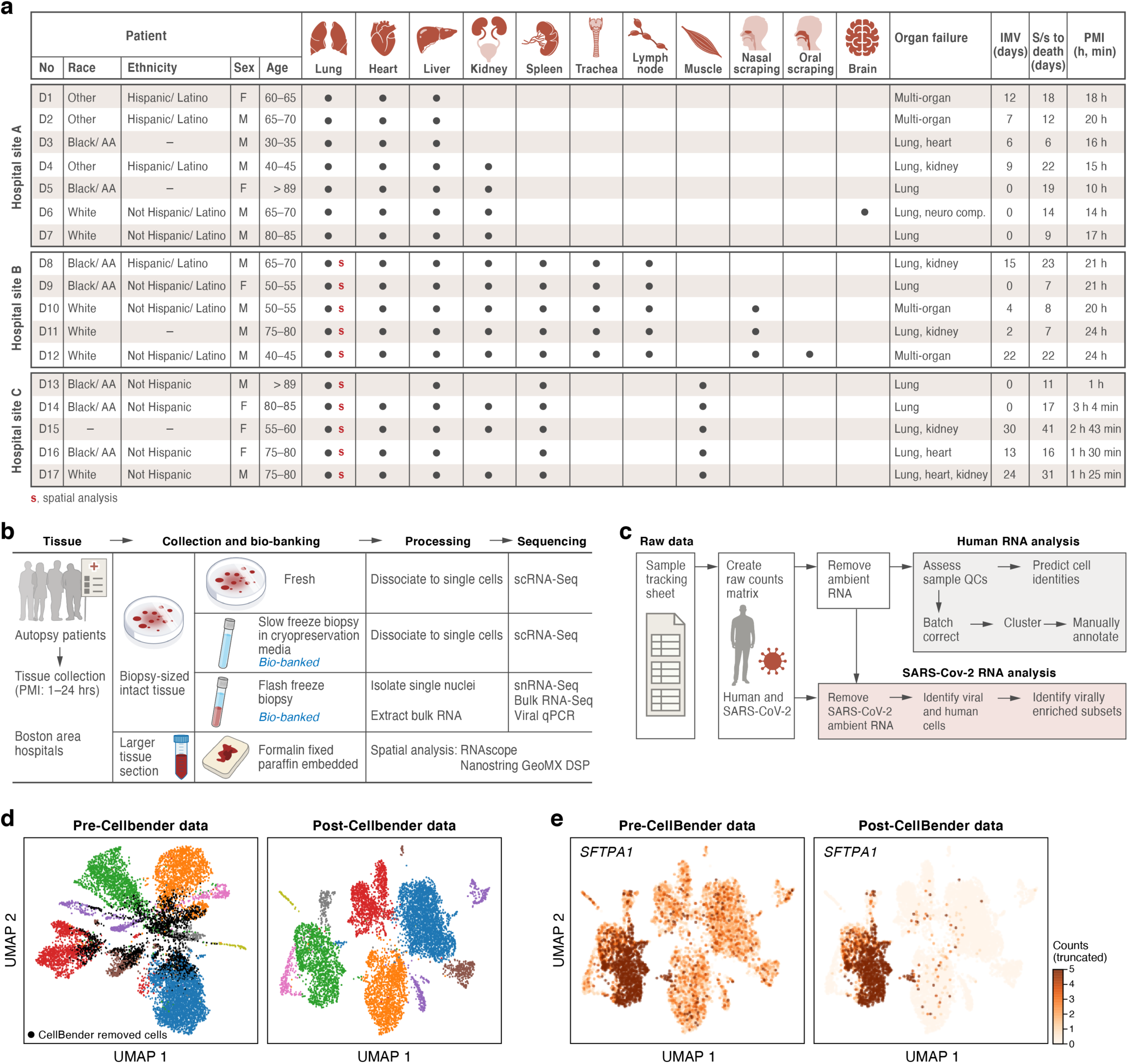
A COVID-19 autopsy cohort for a single cell and spatial atlas. **a.** Cohort overview. IMV: intermittent mandatory ventilation days, S/s: time from symptom onset to death in days; PMI: post-mortem interval. Red bold s: donors for which we collected spatial profiles in the lung. **b**. Sample processing pipeline overview. **c.** sc/snRNA-Seq analysis pipeline overview. **d,e.** CellBender ‘remove-background’ improves cell clustering and expression specificity by removing ambient RNA and empty (non-cell) droplets. UMAP plot of sc/snRNA-Seq profiles (dots) either before (left) or after (right) CellBender processing, colored by clusters and by doublet status (black) (**d**), or by expression of the surfactant protein *SFTPA1* (**e**). Color scale in **e** is linear and truncated at 5 counts to visualize small counts.

To overcome long PMI (up to 24 hours) and differences in tissue properties, we optimized protocols to generate high-quality single-cell RNA-Seq (scRNA-Seq) and snRNA-Seq data within the constraints of a Biosafety Level 3 (BSL3) facility, and developed workflows to profile RNA from different cellular compartments spatially while distinguishing regions with viral RNA with NanoString GeoMx (**Methods**). More specifically, upon receiving specimens at the Broad, we dissected tissue from each organ and collected multiple biopsies from each in a BSL3 facility using a specimen processing pipeline (Fig. 1b) that included: (**1**) dissociating biopsies into single-cell suspensions followed by immediate scRNA-Seq profiling; (**2**) viably freezing biopsies for future scRNA-Seq; and (**3**) flash freezing biopsies for future snRNA-Seq, bulk RNA-Seq, and viral qPCR (**Methods**).

In total, we created a biobank of over 420 specimens across tissues and donors using this pipeline to empower downstream analyses and future follow-up studies (Fig. 1b). Here, we specifically analyzed four tissue types (lung, heart, liver, kidney) in up to sixteen donors by snRNA-Seq and/or scRNA-Seq. We also examined fourteen COVID-19 autopsy donor lung tissues and three controls by Nanostring GeoMx® Digital Spatial Profiler (DSP). For nine donors (D8, D10-D17), we generated matched sc/snRNA-Seq and GeoMx data.

### An atlas of affected tissues from COVID-19 autopsies at single-cell resolution

We generated single-cell/single-nucleus atlases of the lung (*n*=16 individuals, *k*=106,792 cells/nuclei), heart (*n*=15, *k*=36,662), liver (*n*=16, *k*=47,001) and kidney (*n*=11, *k*=29,568). We initially tested both scRNA-Seq and snRNA-Seq on tissue samples collected from COVID-19 autopsies through methods we recently established for analyzing human tumor biopsies and resections^29, 30^ (**Methods**). In these pilots, snRNA-Seq, which is well-suited for processing hard-to-dissociate or damaged tissues, performed better in systematically capturing the complex tissue cellular ecosystem (Supplemental Fig. 1 and *data not shown*) and was thus chosen for profiling most COVID-19 autopsy samples.

We developed a computational pipeline (Fig. 1c) to tackle key technical challenges posed by this dataset. These include: (**1**) ambient RNA contamination, which can reduce the cell type-specificity of transcriptional profiles; (**2**) the need to efficiently and uniformly process a large dataset across tissues; and, (**3**) the inclusion of diverse tissues with many cell types, which we needed to annotate systematically to allow comparisons between donors and to existing reference atlases. First, the long PMI associated with many autopsies in this cohort likely promoted significant cell death and tissue damage. As a result, we observed substantial amounts of ambient RNA in our sc/snRNA-Seq profiles, reflected by reduced separation of cell subsets in low dimensionality projections relative to what is observed in typical tissue atlases (*e.g.*, Fig. 1d), and in lower cell type specificity for known marker genes (*e.g.*, Fig. 1e, Supplemental Fig. 2). To address this, we used CellBender remove-background^32^ (**Methods**), which removes ambient RNA and likely empty (non-cell) droplets from droplet-based sc/snRNA-Seq based on a principled generative model of the various technical errors and contaminants in such data. This improved separation of cell subsets and increased the specificity of marker gene expression among key cell subsets (*e.g.*, AT1 and AT2 cells in lung, Fig. 1d,e, Supplemental Fig. 2).

Second, to efficiently and systematically process our data, we relied on the cloud-based platform Cumulus^33^, which provides a standard pipeline for processing sc/snRNA-Seq data at scale with cloud computing resources. Cumulus generated gene-count matrices, filtered low quality cells, reduced dimensions, clustered, and generated UMAP visualizations of the data. The sample-specific results allowed rapid sample quality control and preparation for integrated analyses.

Third, we devised an automated approach to annotate individual cells/nuclei by type through transferring labels from previously annotated datasets of matched tissues from diverse sources (*e.g.*, healthy or other diseases; different organ regions; single-cell or single-nucleus) to our unlabeled expression profiles. Briefly, we trained a logistic regression classifier on individual expression profiles from public sc/snRNA-Seq datasets of matched tissues to assign cell types to individual cells/nuclei without cell clustering or prior knowledge of marker genes (Fig. 2a, **Methods**). To further refine these cell type labels, we also performed unsupervised sub-clustering on batch-corrected integrated data for each of the main cell lineages and manually annotated sub-clusters using known lineage markers and established gene signatures (Fig. 2b-g, Supplemental Fig. 3, 4, 5, **Methods**). The automated annotation approach allowed us to compare cells/nuclei to other data resources, including atlases of the same tissues in health and disease, and was conducted on a per-cell/nucleus basis (without the need for batch correction). Our manual annotation, meanwhile, enabled us to refine cell identity assignments with detailed domain knowledge to generate richer labels and describe clusters of cell states that may be specific to COVID-19 host-immune response and thus would not be readily captured through automated annotation using non-COVID-19 atlases.

**Figure 2.**
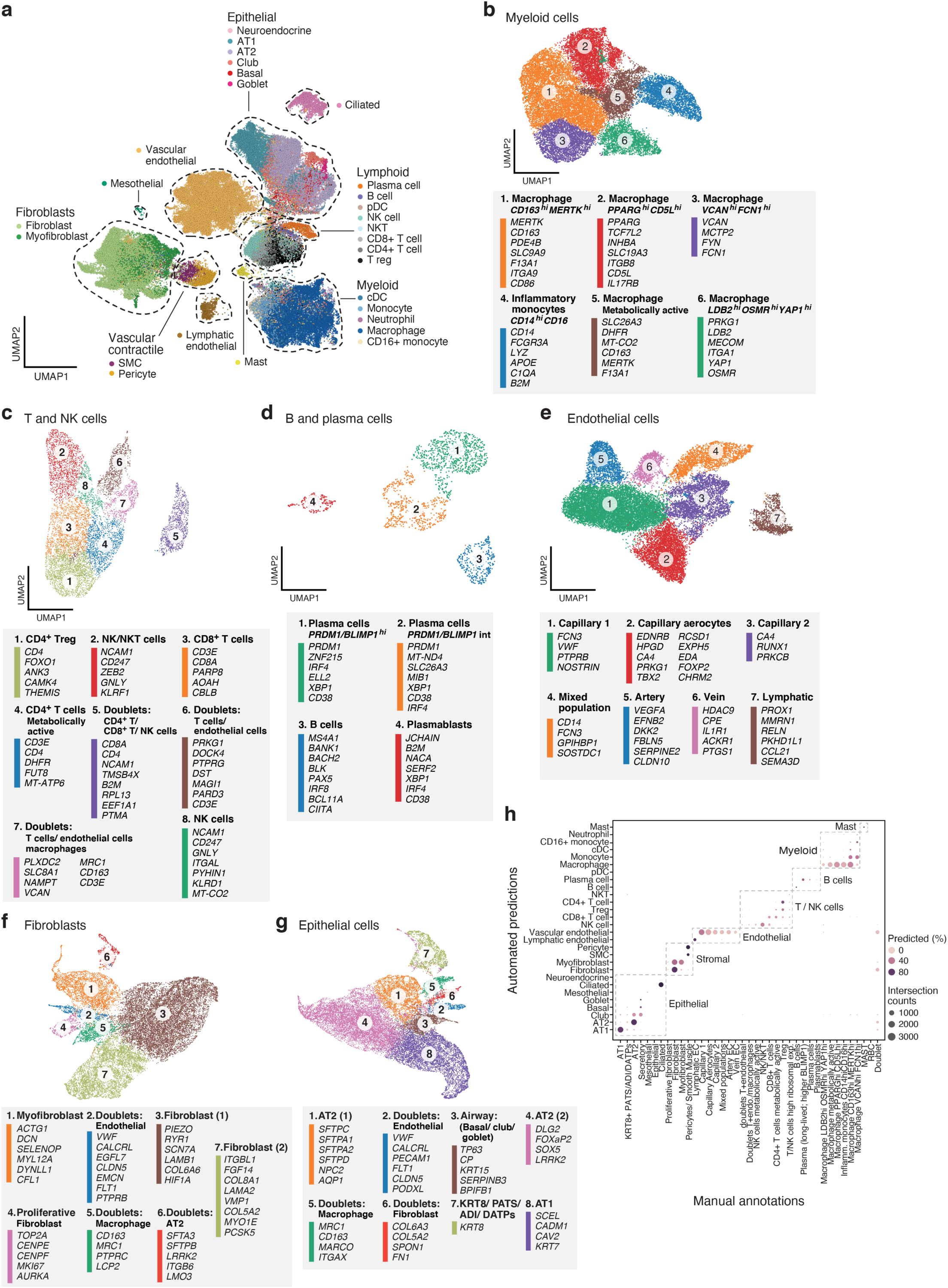
A single cell and single nucleus atlas of COVID-19 lung. **a.** Automatic prediction identifies cells from 28 subsets across epithelial, immune and stromal compartments. UMAP embedding of 106,792 harmonized scRNA-Seq and snRNA-Seq profiles (dots) from all 16 COVID-19 lung donors, colored by their automatically predicted cell type (legend). **b-g**. Refined annotation of cell subsets within lineages. UMAP embeddings of each selected cell lineages with cells colored by manually annotated sub-clusters. Color legends highlight highly expressed marker genes for select subsets. **b.** myeloid cells (24,417 cells/ nuclei), **c.** T and NK cells (9,950), **d.** B and plasma cells (1,693), **e.** endothelial cells (20,366), **f.** fibroblast (20,925), **g.** epithelial cells (21,700). **h.** High consistency between automatic and manual annotations. The proportion (color intensity) and number (dot size) of cells with a given predicted annotation (rows) in each manual annotation category (columns).

### A cell census in the COVID-19 lung spanning epithelial, immune, and stromal cells

We assembled a COVID-19 lung atlas comprised of 106,792 sc/snRNA-Seq profiles (99,735 single nucleus, 4,608 cryopreserved single cells, and 2,449 freshly processed single cells) from 24 lung tissue samples from 16 COVID-19 deceased donors (Fig. 1a, Fig. 2, Supplemental Fig. 3). Following batch correction with Harmony^34^ (**Methods**), cells/nuclei formed distinct clusters that were generally well-mixed across donors and lab protocols (Supplemental Fig. 3a-c, adjusted rand index between sample and cluster at 3.8%). Thus, despite variation between donors, intrinsic cell profiles generalized well across samples, and all major parenchymal, endothelial, and immune cell subsets were captured. We defined 28 cell subsets based on annotations automatically transferred from six sc/snRNA-Seq lung datasets (for cell types present in at least two datasets), spanning different lung regions from healthy donors and from donors with lung fibrosis, with the classifier capturing canonical cell type markers well (Fig. 2a, **Supplemental Table 2**, **Supplemental Table 3**). The annotations were robust by cross-validation (∼80% accuracy on two held out datasets), and cell type mis-assignments were typically between cell subtypes from the same lineage. For finer annotations, especially of cell states that may not have been observed in the reference datasets (*e.g.*, in immune and endothelial cells), we partitioned cells (after batch correction) into six main cell subgroupings – epithelial cells, endothelial cells, fibroblasts, myeloid cells, T and NK cells, and B and plasma cells – and annotated each by sub-clustering and manual analysis (Fig. 2b-g, Supplemental Fig. 4b-e, **Methods**).

In the immune subgroupings, among the 24,417 myeloid cells, we distinguished six cell subsets: *CD14*^high^*CD16^h^*^igh^ inflammatory monocytes expressing transcripts with antimicrobial properties (*e.g., LYZ*, *S100A6*); and five macrophage subsets enriched for distinct key immune genes (Fig. 2b, Supplemental Fig. 4d, 5a), including either scavenger receptors (*e.g.*, *CD163, STAB1*), toll-like receptor ligands (*e.g.*, *VCAN*) and inflammatory transcriptional regulators (*e.g.*, *LDB2*, *YAP1*); or genes associated with metabolism (*e.g. DHFR, INO80D*) and higher mtRNA reads. Among the 9,950 T and NK cells, we annotated six subsets including: two CD4^+^ subsets, including a regulatory T cell subset expressing *FOXO1* and *ANK3*, and a metabolically active subset enriched for *DHFR* expression and high mtRNA reads; one CD8^+^ subset; and, two T/NK cell subsets (Fig. 2c, Supplemental Fig. 4d, 5b), including one enriched for cytotoxic effector genes (*e.g.*, *GNLY, PRF1*). In addition to the subsets, there were three doublet clusters, one containing *CD4^+^* T cells, *CD8^+^* T cells and NK cells (cluster 5); one with transcripts from both T and endothelial cells (cluster 6); and one with T cells, endothelial cells, and *VCAN*-expressing macrophages (cluster 7). Finally, among the 1,693 B and plasma cells, we identified four subsets, including: plasma cells expressing high levels of transcription factor *BLIMP-1* (*PRDM1*); plasma cells expressing intermediate levels of *PRDM1*; B cells; and, *JCHAIN* expressing plasmablasts (Fig. 2d, Supplemental Fig. 4d).

In the stromal grouping, among 21,391 endothelial cells (ECs), we identified seven cell subsets (Fig. 2e): Three were annotated using known signatures^35^ as arterial endothelial cells (cluster 5), venous endothelial cells (cluster 6) and lymphatic endothelial cells (cluster 7, Supplemental Fig. 4d, 4e). The remaining four subpopulations were identified using imputation and comparison to other lung and endothelial cell atlases^35, 36^ (Supplemental Fig. 4f). They included: a capillary aerocyte subpopulation (cluster 2) based on expression of *EDNRB, HPGD, CA4,* and *PRKG1*; a capillary EC-1 population (cluster 1) by *VWF* and *PTPRB*; a capillary EC-2 population (cluster 3) by *RUNX1, CD44,* and mitochondrial genes, a pattern typically observed in states of cellular stress or dying cells; and a EC subpopulation (cluster 4) whose gene expression signature overlaps with multiple EC subpopulations in the healthy lung reference data and is enriched with predicted doublets (**Methods**). This unidentified subpopulation was also rich in whole cells (fresh and cryopreserved) compared to the more specific nuclear preparation of the other EC subpopulation (Supplemental Fig. 4g). Within the fibroblast lineage, among 20,925 cells, we identified fibroblasts, proliferative fibroblasts, myofibroblasts, and doublet clusters^35^ (Fig. 2f, Supplemental Fig 4d).

In the epithelial compartment, among 21,700 cells, we observed eight cell clusters, including club/secretory cells, AT1 cells, AT2 cells, a proliferative state of the AT2 cells, and three sets of cell doublets (Fig. 2g). The eighth cluster corresponds to an intermediate cell state (*KRT8+* PATS/ADI/DATP) previously described as a transitional state from AT2 cells to AT1 cells during alveolar regeneration^37–39^ (Fig. 2g, Supplemental Fig. 4h), which we discuss below.

Deconvolution of bulk RNA-Seq profiles from the same lung samples largely agreed with our cell type classifications, suggesting that our snRNA-Seq reflects tissue composition. We first used sn/scRNA-Seq profiles to deconvolve bulk RNA-Seq and infer relative composition across 11 major cell subsets (Supplemental Fig. 5c-e, **Methods**). Most samples had at least 25% inferred fibroblast content, which may indicate lung fibrosis (Supplemental Fig. 5c), and substantial variation in inferred myeloid cell composition, which was consistent between replicates but varied from 4-42% between samples. Conversely, the inferred proportion of epithelial and endothelial cells was often lower (0.05 −38% and 0.03 - 43% respectively). Overall, predicted cell type composition was broadly consistent with sc/snRNA-Seq, identifying the same cell types as most prevalent, with the exception of T+NK cells that we inferred at low proportions in bulk RNA-Seq, but present at ∼10% on average in sc/snRNA-Seq (Supplemental Fig. 5e).

Importantly, our two annotation strategies were highly coherent, such that 94% of the lineage assignments matched between the manual and automatic annotations (Fig. 2h, Supplemental Fig. 5a, 5b). Deconvolution of bulk RNA-Sequencing data also largely agreed with sc/snRNA-Seq cell type classifications (Supplemental Fig. 5c-5e, **Methods**). Manual annotations were particularly important for resolving specific cell states not in the classification model, such as the pre-alveolar type-1 transitional cell state (PATS), *TP63*^+^ intrapulmonary basal-like progenitor cells, and macrophage and endothelial cell subsets; whereas the automatic annotations allowed us to next compare cell compositions to non-COVID-19 lung atlases.

### Substantial increases in immune and stromal cell subsets and depletion of AT2 cells in COVID-19 lung

Although most cell subsets were present in all samples (Supplemental Fig. 3a,b), there was variability in the cellular composition of the lung samples across donors. For example, the proportion of lymphoid cells ranged from 0.4% to 33% across snRNA-Seq samples between donors, and epithelial cells ranged from 3.8% to 52% (Supplemental Fig. 6a,b). These two compartments were strongly anti-correlated (though it is not possible for us to distinguish in such analyses between causal relations and the expected zero-sum game due to sampling).

We also compared the cellular composition of COVID-19 lungs to that of normal lung, by contrasting against a snRNA-Seq lung dataset from a matching tissue region, generated using similar lab protocols and profiling technology in healthy deceased donors (MS, ORR, AR, *unpublished data*), and using the automatic cell type classification model to transfer labels between the two studies (Fig. 3a, **Methods**). The largest change was a significant decrease in the proportion of AT2 cells (p-value = 2*10^-16^), dropping from 40% of cells (the largest population) in healthy lung lobes to 10% in COVID-19 samples (**Methods**). As AT2 cells were previously identified as likely targets of SARS-CoV-2^40–42^, this may reflect widespread, virally-induced AT2 cell death. Conversely, immune cells — including dendritic cells (p-value=0.001), macrophages (p-value = 6.4*10^-11^), and NK cells (p-value = 0.008) — all increased in their relative abundance in severe COVID-19. Fibroblasts (p-value = 0.0041), lymphatic endothelial cells (p-value = 0.0001), and vascular endothelial cells (p-value = 2.3*10^-5^) were also captured at a greater relative abundance in COVID-19 lungs.

**Figure 3.**
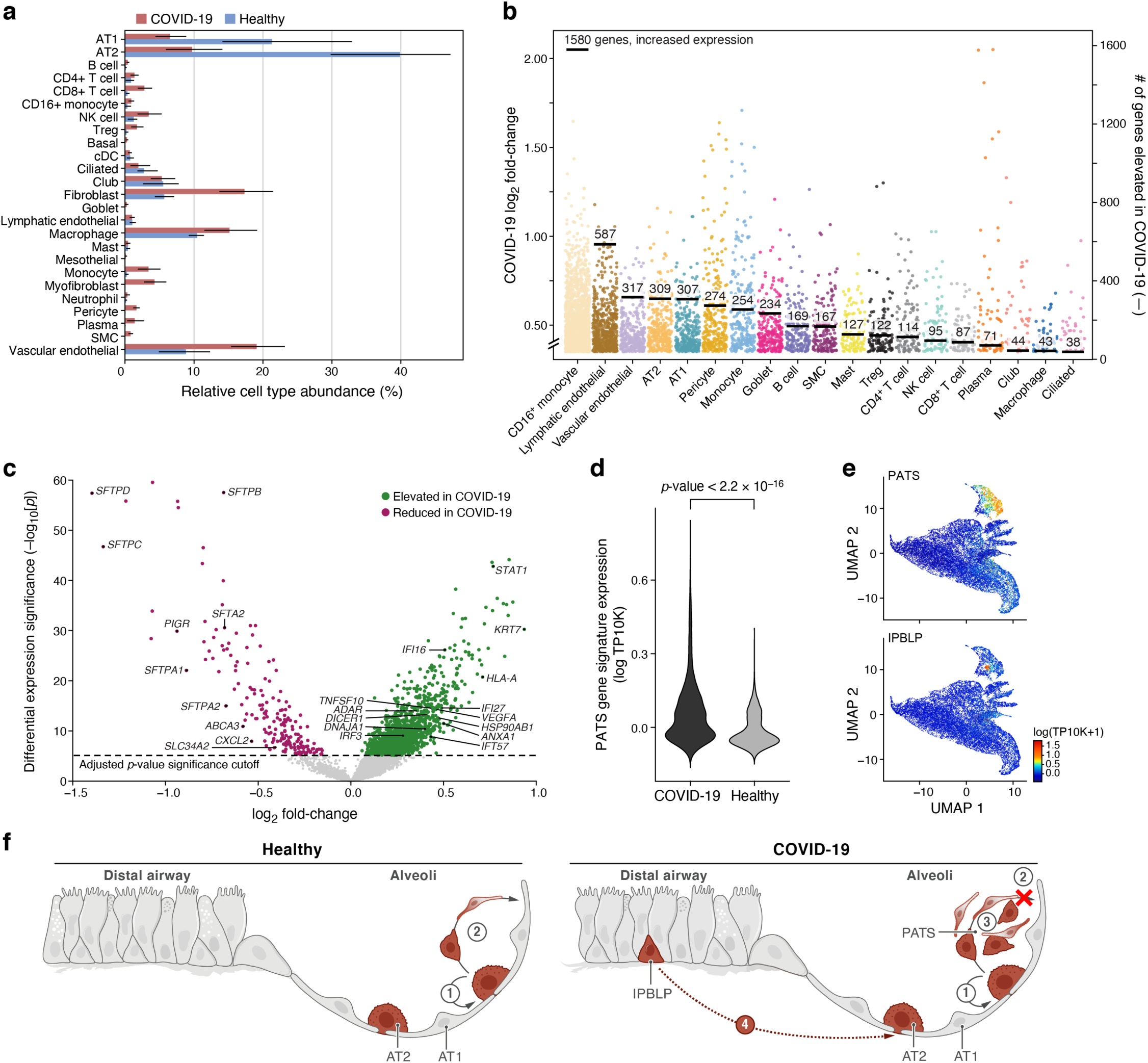
Dramatic remodeling of cell composition and cell intrinsic programs in COVID-19 lung. **a.** Differences in cell composition between COVID-19 and healthy lung. Proportion (*x* axis, mean and 95% confidence intervals) of cells in each subset (*y* axis, by automatic annotation) in COVID-19 snRNA-Seq (red) and a healthy snRNA-Seq dataset (blue). **b,c.** Myeloid, endothelial and pneumocyte cells show substantial changes in cell intrinsic expression profiles in the COVID-19 lung. **b.** Log_2_(fold change) (*y* axis) between COVID-19 and healthy lung for each gene (dot) in each cell subset (*x* axis, by automatic annotation). Black bars: number of genes with significantly increased expression (adjusted p-value < 7.5*10^-6^). **c.** Significance (-Log_10_(P-value), *y* axis) magnitude (log_2_(fold-change), *x* axis) of differential expression of each gene (dots) in 2000 AT2 cells, from a meta-differential expression analysis between COVID-19 and healthy samples across 14 studies. **d.** An increased PATS^37–39^ program in pneumocytes in COVID-19 lung. Distribution of PATS signature scores (*y* axis) for the 17,655 cells from COVID-19 or 24,000 cells from healthy lung (*x* axis). **e.** UMAP embeddings of epithelial cells colored by their expression of cell program signatures (color legend, lower right) for the PATS program (upper panel) and the IPBLP program (lower panel). **f.** Graphical schematic of alveolar cellular turnover. In healthy alveoli (left panel), AT2 cells self-renew (1) and differentiate into AT1 (2). In COVID-19 alveoli (right panel), AT2 cell self-renewal (1) and AT1 differentiation (2) are inhibited, resulting in PATS accumulation (3) and recruitment of airway-derived IPBLP progenitors to alveoli (4).

### Induction of viral, inflammatory and progenitor programs in epithelial cells in COVID-19 lung

To identify cell intrinsic changes in expression associated with severe COVID-19, we created a lung meta-atlas of ∼1,280,000 cells and nuclei from 14 studies (**Supplemental Table 2**), spanning healthy and COVID-19 infected lungs, sampled by biopsy or bronchial alveolar lavage (BAL) and profiled using scRNA-Seq or snRNA-Seq on the 10x Chromium platform. After dataset aggregation, we applied our automated cell annotation to classify each of the 1.28M cells into the same 28 classes identified in our severe COVID-19 lung atlas. We then used a linear regression model to identify differentially expressed genes between COVID-19 and healthy tissues for each abundant cell type (**Supplemental Table 4**, **Methods**).

Differential gene expression showed widespread transcriptional changes in key COVID-19 cell types (Fig. 3b,c). *CD16^+^* monocytes had 1,580 genes with elevated gene expression levels in the late stages of COVID-19. Lymphatic endothelial (578 genes with elevated expression), vascular endothelial (317), AT2 (309), and AT1 (307) cells also showed large transcriptional changes. Some cell types, such as ciliated cells, had very few genes that significantly increased expression (38), showing less involvement in the infection’s late stages.

Within AT2 cells, there was higher expression of genes associated with host viral response (Fig. 3c), including programmed cell death genes (*e.g.*, *STAT1,* p-value = 1.6*10^-43^), and inflammation and adaptive immune response genes (*e.g.*, *EREG* with p-value = 3.2*10^-14^*, CTSE* with p-value = 2.1*10^-2*5*^*, TNFSF13B* with p-value = 2.5*10^-7^, and *SAMD9* with p-value = 8.1*10^-14^). Gene sets associated with cell migration (*e.g.*, *ITGA3* with p-value = 1.3*10^-11^, *AGRN* with p-value = 1.6*10^-14^) and damage response (*e.g.*, *TAOK1* with p-value = 1.6*10^-9^) were also more highly expressed in AT2 cells from COVID-19 lung tissue. By comparison, expression of genes linked to proliferation and apoptosis regulation (*e.g.*, *CD63* with p-value = 7.5*10^-16^, **Supplemental Table 4)** were significantly reduced in COVID-19 lung tissue. Expression of lung surfactant genes were also reduced in COVID-19 lung tissue (*SFTPD, SFTPC, SFTPA1, SFTPB, SFTA2, SFTPA2*), with log_2_ fold changes between −1.4 and −0.6 with p-value < 10^-15^, showing that a phenotype previously reported *in vitro*^38^ is also observed *in vivo*.

Notably, we saw an increase in the expression of the PATS program signature in epithelial cells in COVID-19 lungs compared to healthy lungs (p-value < 2.2*10^-16^, one-sided Mann–Whitney U test), consistent with prior studies showing that this progenitor program is induced during lung injury^37–39^ (Fig. 3d, 3e). Interestingly, these studies all reported that this intermediate state expands in lung diseases, such as idiopathic pulmonary fibrosis (IPF), and lung fibrosis has been documented in patients with severe COVID-19^43, 44^. These studies also highlighted the expansion of myofibroblasts, which we also observe in COVID-19 lungs (Fig. 3a). Additionally, we detected a subset of cells among those that express the PATS program which shares the expression of PATS markers (*KRT8*/*CLDN4*/*CDKN1A*) but also expresses *KRT5*, *TP63*, and *KRT17*, which are not canonical features of the PATS program (Supplemental Fig. 4h, top **Supplemental Table 5**). Notably, these cells are also distinct from *KRT5*^+^/*TP63*^+^ airway basal cells (Supplemental Fig. 4h, bottom). We thus hypothesize that these cells may represent TP63^+^ intrapulmonary basal-like progenitor cells (IPBLP), which were previously identified in H1N1 influenza^45^, and are thought to be an emergency cellular reserve for severely damaged alveoli (Fig. 3e, 3f)^46^. Compared to airway basal cells, the putative IPBLP cells are characterized by the expression of numerous interferon viral-defense pathway genes (*IFI27, IFITM1, IFITM2, IFITM3, IFI6, ISG15, BST2*) as well as genes involved in the differentiation of progenitor cells (*S100A11, PPDPF, S100A16, TNFRSF12A*).

### Viral burden detected by sc/snRNA-Seq and RT-PCR is associated with changes in lung cell composition

To determine SARS-CoV-2 viral load and its association with host responses, we examined the donor and cell type-specific distribution of reads that aligned to the SARS-CoV-2 genome (Fig. 4a-4m, Supplemental Fig. 7**, Methods**). Here, we found substantial variation between donors: in 13 donors (14 samples), we detected at least one SARS-CoV-2-aligned unique molecular identifier (UMI) (from 0 - 4,731 distinct viral UMI per lung, Fig. 4a, Supplemental Fig. 7a). Viral-aligning UMI spanned the entire SARS-CoV-2 genome, and were biased toward positive-sense alignments, with a few cells containing reads aligning to all 28 viral segments, including the negative strand (Supplemental Fig. 7d), which may indicate productive infection. Notably, viral detection from single-cell or single-nucleus transcriptomic data was not driven by confounding technical factors on a per-cell or per-sample basis (Supplemental Fig. 7e-7h). This inter-donor variation reflected the SARS-CoV-2 burden in the tissue microenvironment, as estimated by SARS-CoV-2 RT-PCR on bulk RNA from directly adjacent biopsy-sized tissue pieces (Fig. 4b, Supplemental Fig. 7i, 7j). Importantly, viral load (number of SARS-CoV-2 copies/ng RNA by RT-PCR) in lung parenchyma was negatively correlated to the time interval between donor-reported symptom onset and death (spearman’s rho = −0.68, p-value = 0.005, Fig. 4f), consistent with previous reports ^47, 48^.

**Figure 4.**
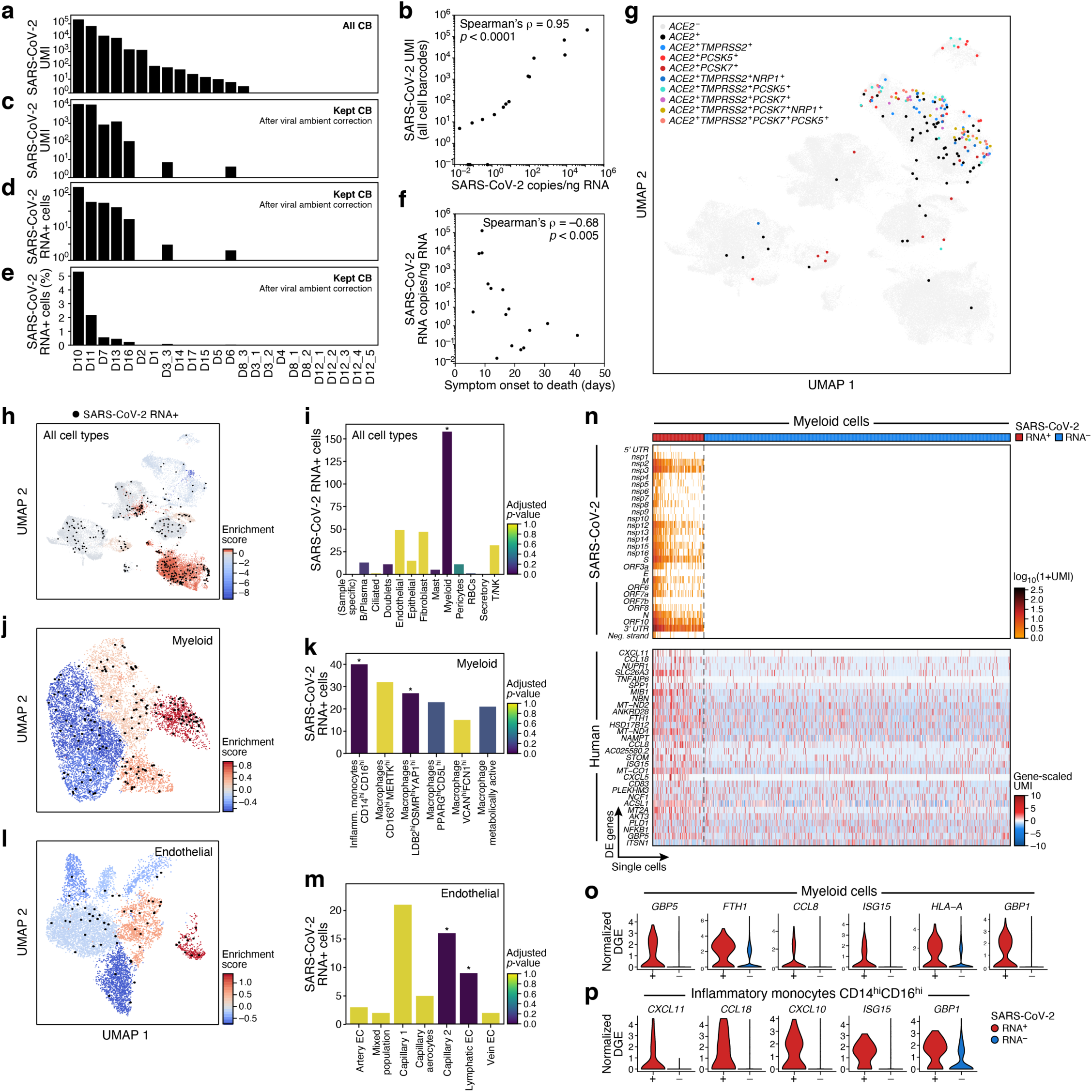
SARS-CoV-2 RNA+ single cells span multiple lineages and are enriched among phagocytic and endothelial cells. **a-e.** Robust identification of SARS-CoV-2 RNA+ single cells. **a.** Number of SARS-CoV-2 UMIs from all cell barcodes (*y* axis). **c.** Number of SARS-CoV-2 UMIs after ambient correction. **d.** and **e.** Number (**d**, *y* axis; Range: 1-169, total: 342, mean +/- SEM: 49 +/- 22) and percent (**e**, *y* axis, Range: 0.02-5.3%, mean +/ SEM 1.25% +/- 0.73%) of SARS-CoV-2+ RNA cells (both after ambient correction), across the samples (*x* axis), ordered by ranking in **a**. **b.** Agreement of overall viral RNA abundance from sc/snRNA-Seq and qPCR on bulk RNA. SARS-CoV-2 copies as measured by CDC N1 qPCR assay on bulk RNA extracted from matched tissue samples (*x* axis) and from number of all SARS-CoV-2 aligning UMI (pre-ambient correction, *y* axis). **f.** Reduction in SARS-CoV-2 RNA with more prolonged S/s interval. Interval between symptom onset and death (*x* axis, days) and lung SARS-CoV-2 copies/ng input RNA (*y* axis) for each donor. **g.** SARS-CoV-2 RNA+ single cells are not closely related to expression of the SARS-CoV-2 entry factors. UMAP embedding (as in Fig. 2a) of all cells/nuclei profiled in the lung, colored by single and multi-gene expression of the SARS-CoV-2 entry factor *ACE2* with different accessory proteins. Co-expression combinations with at least 10 cells are shown. **h-m.** SARS-CoV-2 RNA+ cells are enriched in specific lineages and sub-types. Left panels, **h**, **j**, **l**: Cells from 7 donors containing any SARS-CoV-2 RNA+ cell, and colored by viral enrichment score (color bar; red: stronger enrichment) and by SARS-CoV-2 RNA+ cells (black points). UMAP embeddings of either all cell types (**h**, as in Supplemental Fig. 4a), myeloid cells (**j**, as in Fig. 2b), or endothelial cells (**l**, as in Fig. 2e). Right panels, **i**, **k**, **m**: Number of SARS-CoV-2 RNA+ cells (*y* axis) per cell type/subset (*x* axis), with bars colored by enrichment score (color bar. dark blue: stronger enrichment). * FDR < 0.01. **n-p**. Expression changes in SARS-CoV-2 RNA+ myeloid cells. **n.** Expression of SARS-CoV-2 genomic features (top, log-normalized UMI counts; rows) and significantly DE host genes (bottom, log-normalized and scaled digital gene expression, rows; FDR-corrected p-value < 0.05 and log_2_ fold change > 0.5) across SARS-CoV-2 RNA+ and SARS-CoV-2 RNA-myeloid cells (columns). **o,p.** Distribution of normalized digital expression levels (*y* axis) for select significantly DE genes between SARS-CoV-2 RNA- and SARS-CoV-2 RNA+ cells from myeloid cells (**o**) or from Inflammatory monocytes *CD14*^high^*CD16*^high^ cells (**p**).

Total viral burden per sample (Fig. 4a, 4b, including ambient RNA, **Methods**) was associated with significant changes in lung-resident cell populations. In particular, there was significant positive correlation between viral burden and the proportion of mast cells (Bonferroni-corrected p-value (q) = 0.012, Pearson’s r = 0.67), *VCAN*^high^*FCN1*^high^ macrophages (q = 0.015, r = 0.66), *CD163*^high^*MERTK*^high^ macrophages (q = 0.021, r = 0.65), *LDB2*^high^*OSMR*^high^*YAP1*^high^ macrophages (q = 0.029, r = 0.64), venular endothelial cells (q = 0.038, r = 0.62) and capillary aerocytes endothelial cells (q = 0.049, r = 0.61) (Supplemental Fig. 7k-q).

Differential expression analysis of bulk RNA-Seq profiles from adjacent lung biopsies comparing “highly infected” (viral RNA copies/ng total RNA > 5*10^3^; samples: D7, D10, D11) and “uninfected” (< 1.5 viral RNA copies/ng total RNA; samples: D4, D6, D8, D15, D17) samples (**Methods**), highlighted viral response and innate immune processes. Specifically, genes upregulated (log_2_FC > 1.4, Wald test, FDR-corrected p-value < 0.05) in highly infected specimens included those associated with host response to virus (*IFIT1, ISG15, IFIT3, APOBEC3A,* and *PRF1*), and innate immune and effector responses (*LAG3, GZMB, GREM, CCL7,* and *SEMA4A*) (Supplemental Fig. 7r). These genes were enriched (Kolmogorov-Smirnov statistic, FDR q-value = 3.12E*10^-6^) with genes upregulated in post-mortem lung tissue of COVID-19 donors compared to uninfected biopsies^49^ (e.g., *ISG15, IFIT1, IFIT3, GBP5, CCL7, APOBEC3A*). Genes significantly down-regulated (log_2_FC < 1.4, Wald test, FDR-corrected p-value < 0.05) in highly infected lung tissue were involved in lung fibrosis, surfactant metabolism dysfunction, and lamellar body function (*SFTPA1, SFTPA2, SFTPC, LAMP3*), which serve as secretory vesicles in AT2 cells^50, 51^.

To evaluate viral genetic diversity, we performed metagenomic sequencing on bulk RNA collected from adjacent biopsy-sized tissue from 16 donors. We assembled nine complete genomes from the six unique donors with the highest viral loads. When compared to a previously-published database of SARS-CoV-2 genomes from Massachusetts between January-May 2020, the six genomes derived from this cohort are broadly distributed across local circulating lineages (Supplemental Fig. 7s) and all carry the “G” allele at D614G, which represents the dominant variant observed in Massachusetts. Finally, metagenomic classification of reads from bulk genomic data did not identify co-infections from common respiratory pathogens such as influenza, rhinovirus, enterovirus, metapneumovirus, or other seasonal coronavirus strains (Supplemental Fig. 7t), confirming the expected clinical history.

### SARS-CoV-2 RNA+ cells span multiple subsets, but are largely represented by myeloid cells

Cell types with SARS-CoV-2 RNA+ cells spanned myeloid cells, B and plasma cells, epithelial cells, fibroblasts, pericytes and T/NK cells, with only myeloid cells being significantly enriched (Fig. 4g-4i, Supplemental Fig. 7u-7w). Notably, the presence of SARS-CoV-2 RNA+ cells did not overlap with co-expression of the SARS-CoV-2 entry factors *ACE2* and *TMPRSS2*, nor did they correspond with an expanded set of novel or hypothesized proteases and cofactors thought to contribute to SARS-CoV-2 entry, including *FURIN, CTSL, PCSK7, PCSK2, PCSK5* and/or *NRP1* (Fig. 4g, 4h). We attribute this finding to the lack of expression of *ACE2* among myeloid subsets (the largest contributor of SARS-CoV-2 RNA+ cells), as well as the relative loss of viable epithelial cells (previously identified as highest expressors of *ACE2* among lung resident cell types) among the donors with highest abundances of SARS-CoV-2 RNA (*e.g.*, D10 and D11, Supplemental Fig. 8a-d).

To define SARS-CoV-2 RNA+ cells, we corrected for the variable amount of SARS-CoV-2 reads within the ambient RNA pool with a virus-specific ambient RNA removal step (combining CellBender with a previously described approach^52^ to achieve a more stringent cutoff for cell-associated viral RNA, **Methods**). Following correction and filtering, we retained seven samples (donors: D3, D6, D7, D10, D11, D13 and D16, all from snRNA-Seq data) (Fig. 4c-e, Supplemental Fig. 7b, 7c), and computed an enrichment score to quantify the proportional abundances of SARS-CoV-2 RNA+ cells within each manually-annotated cell type category (Fig. 4h-i, Supplemental Fig. 7u**, Methods**). Myeloid cells were enriched for SARS-CoV-2 RNA+ cells (Fig. 4i, 158 SARS-CoV-2 RNA+ cells, adjusted FDR < 0.001). Mast cells, B cells, and plasma cells had an elevated enrichment score compared to other lineages, but these were not significant (Fig. 4i). Viral RNA+ cells were also found among many other cell types, including endothelial, fibroblasts, AT1, AT2, T/NK, pericytes and myofibroblasts, but none were significantly enriched (Fig. 4h,i). Moreover, cells containing UMIs aligning to the viral negative strand, SARS-CoV-2 UMI/ cell > 100, and SARS-CoV-2 genomic features/cell > 20 - all metrics associated with high confidence in cell-associated viral RNA - were largely found among myeloid cells, and rarely within each of the other listed cell types.

Viral enrichment scores varied among the six myeloid subsets (Fig. 4j,k, Supplemental Fig. 7v, 8e). In particular, SARS-CoV-2 RNA+ cells were significantly enriched within “Inflammatory monocytes *CD*14^high^*CD*16^high^” (adjusted FDR < 0.001, 40 cells, represented by 5 donors) and “Macrophages *LDB2*^high^*OSMR*^high^*YAP1*^high^” (adjusted FDR < 0.009, 27 cells, 5 donors) (Fig. 4j). When calculating enrichment scores in each donor individually (to account for different cell subset proportions across donors), we found large inter-individual variability (Supplemental Fig. 8c-f) and significant hits despite small cell numbers. For example, “Inflammatory monocytes *CD14*^high^*CD16*^high^” were enriched among three donors and “Macrophages *LDB2*^high^*OSMR*^high^*YAP1*^high^” in two donors (Supplemental Fig. 8e). Although each donor displayed relative enrichment of SARS-CoV-2 RNA+ cells among some myeloid subsets, viral RNA containing cells were often present in multiple myeloid subsets within the same sample (Fig. 4j,k, Supplemental Fig. 7v, 8e).

Endothelial cells were the next most abundant cell type containing SARS-CoV-2 RNA+ cells (49 cells, 5 donors), but were not significantly enriched (Fig. 4i). Within endothelial cells, the mixed population subset (cluster 3, adjusted FDR < 0.007, 16 cells, 4 donors) and lymphatic endothelial cells (cluster 7, adjusted FDR < 0.001, 9 cells, 3 donors) were enriched compared to other endothelial subsets (Fig. 4l,m and Supplemental Fig. 7w). Within individual donors, lymphatic endothelial cells were enriched in SARS-CoV-2 RNA+ cells in one donor (D10: adjusted FDR < 0.014, 6 cells) and the capillary 2 subset in two others (D7: adjusted FDR < 0.001, 4 cells, D11: adjusted FDR < 0.001, 9 cells) (Supplemental Fig. 8f).

### Intrinsic and innate immune responses among SARS-CoV-2 RNA+ cells

Next, we tested whether SARS-CoV-2 RNA+ cells had distinct transcriptional programs compared to RNA-counterparts in the same sample and cell type, focusing on subsets with a large enough number of SARS-CoV-2 RNA+ cells (myeloid, epithelial, T/NK, B/Plasma, and endothelial cells, and fibroblasts and select sub-clusters; **Methods**). We found significantly differentially expressed genes (FDR-corrected p-value < 0.05; **Methods**) between SARS-CoV-2 RNA+ and SARS-CoV-2 RNA-cells in epithelial and myeloid cells, and the “Macrophages *PPARG^high^CD15L^high^*” and “Inflammatory monocytes *CD14*^high^*CD16*^high^” subsets (**Supplemental Table 6**).

Genes upregulated in epithelial SARS-CoV-2 RNA+ cells (compared to SARS-CoV-2 RNA-epithelial cells in matched donors) were enriched for TNF signaling, chemokine/cytokine signaling, SARS-CoV-2 driven cell responses following *in vitro* infection^49^, AP1 signaling and pathways involved in keratinization, which may reflect pneumocyte specific responses to injury, and included key chemokines (*e.g.*, *CXCL2* and *CXCL3*), early-response genes, and transcriptional regulators (*e.g.*, *CREB5, JUN, EGR1, ZBTB10, NFKBIA* and *TCIM*) (Supplemental Fig. 8g). Genes upregulated in myeloid SARS-CoV-2 RNA+ cells were enriched for chemokine and cytokine signaling, and responses to interferon, TNF, intracellular pathogens, and viruses (**Methods**, **Supplemental Table 6**, Supplemental Fig. 8h), and included *CXCL11, CXCL10, CCL8, CCL18,* and other interferon-responsive factors (e.g., *ISG15, GBP5, TNFAIP6, GBP1* and *NFKB1*) (Fig. 4n,o, **Supplemental Table 6**). Within “Inflammatory monocytes *CD14*^high^*CD16*^high^”, *CXCL11, CCL18, CXCL10*, *TNFAIP6, ISG15,* and *GBP1* were upregulated in SARS-CoV-2 RNA+ cells (Fig. 4p, **Supplemental Table 6**), and among “Macrophages *PPARG^high^CD15L^high^*”, *NT5DC1*, *NCF1, FLT1*, *CXCL11* and *TNFAIP6* were all upregulated (**Supplemental Table 6**).

### A digital spatial profiling atlas of COVID-19 lung

To put our single-cell atlas in the tissue context and evaluate spatial expression patterns in autopsy tissues, we used Nanostring GeoMx Digital Spatial Profiling (DSP) for highly multiplexed proteomic or transcriptomic profiling from user-defined regions of interest (ROIs) (**Methods**). We profiled a total of fourteen lung tissues: nine COVID-19 donors also studied by snRNA-Seq (D8, D10, D11, and D12; LUL and D13-D17; upper lobe (UL)) and five LUL tissues from COVID-19 donors with no accompanying snRNA-Seq (D9, D18-D21; LUL). We also profiled right upper lobe (RUL) from D18; left lower lobe (LLL) from D19; heart left ventricle from D20; and parenchyma from three SARS-CoV-2 negative patients (D22-D24) (Supplemental Fig. 9, 10a). We stained serial sections from the same FFPE block with an RNAscope probe against the viral genome *S* gene; a GeoMx^®^ Cancer Transcriptome Atlas Panel (CTA, 1,811 genes); an early-access GeoMx^®^ Whole Transcriptome Atlas Panel (WTA, 18,335 genes); and, a DSP protein panel (77 proteins). The CTA and WTA panels were supplemented with 26 human genes associated with SARS-CoV-2 biology (**Methods**). We selected ROIs in the WTA and CTA slides based on the *S* gene RNA probe and a 3-color nuclear (Syto13), immune (CD45), and epithelial (Pan-cytokeratin; PanCK) staining of the investigated slide (Fig. 5a**, Methods**). We also performed a similar protocol using five additional FFPE blocks from subjects D13-D17, where GeoMx^®^ WTA ROIs were selected based on tissue type (bronchial epithelium, artery, alveoli), and where serial sections were DAB stained with an anti-SARS-CoV-2 spike glycoprotein antibody and RNAscope probes for SARS-CoV-2 to complement ROI selection (Supplemental Fig. 11a, **Methods**). We chose ROIs that spanned a range of anatomical structures and viral abundance levels based on RNAscope signals, and, when possible, segmented them to PanCK^+^ and PanCK^-^, inflamed and normal-appearing alveoli areas of interest (AOIs) (Fig. 5a, bottom, Supplemental Fig. 11, **Methods**); We did not use CD45 staining for segmentation due to high off-target binding. We then acquired profiles (WTA or CTA) from matched AOIs for all assays based on distance to morphological landmarks (**Methods**).

**Figure 5.**
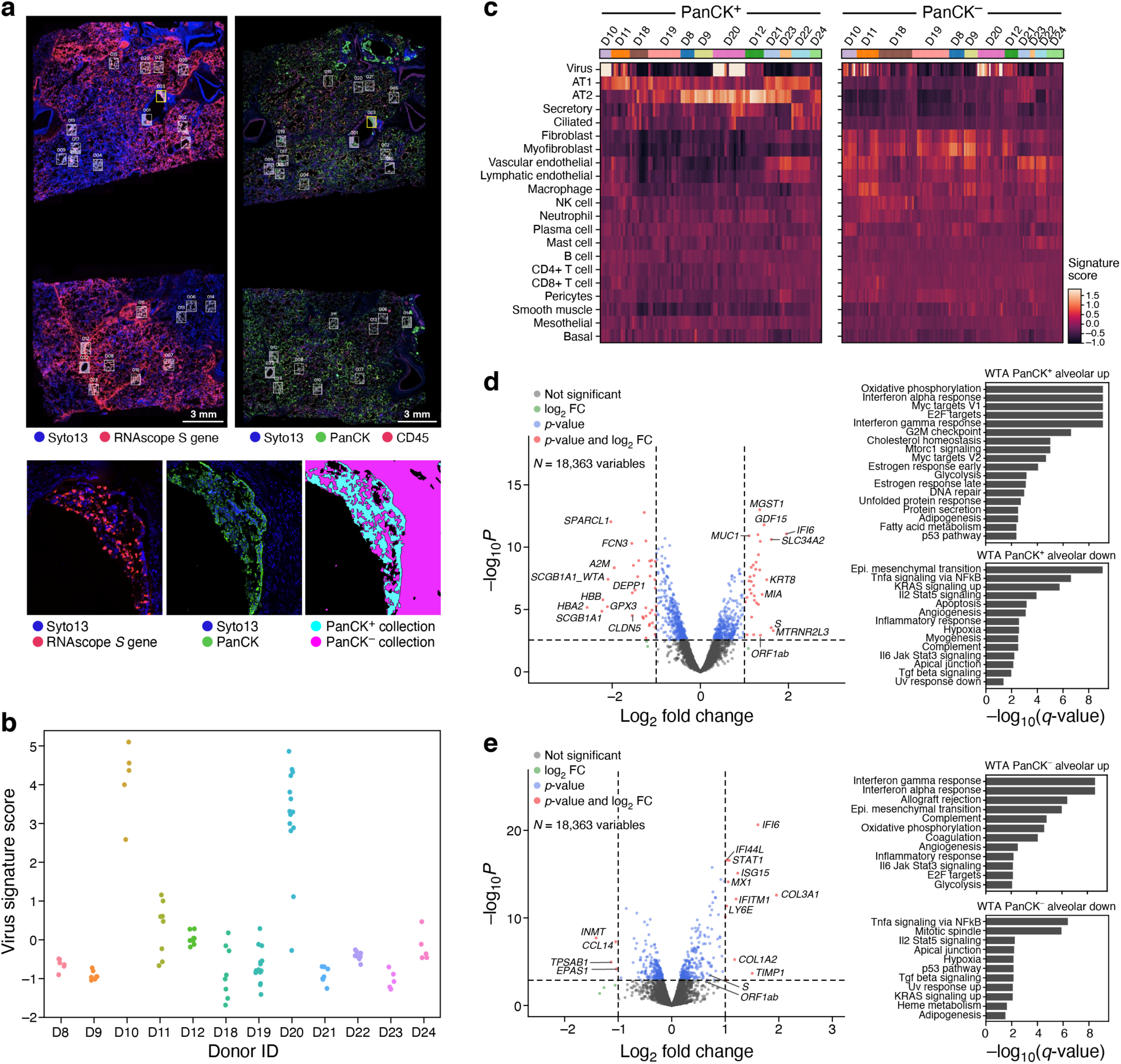
Spatial atlas of autopsy lung samples highlight composition and expression differences between infected and uninfected regions and with healthy lungs. **a.** Selection of ROIs and AOIs. Top: overview scans of donor D20 showing *S* gene RNAscope (left) and immunofluorescent staining (right). ROIs for CTA collection (right) are associated with RNAscope images (left) (white rectangles). Bottom: Zoom in of one ROI (yellow rectangle) from each scan (left and middle), and the defined segmentation masks for collection (right). **b.** Differences in viral load within regions and between donors. Viral signature score (*y* axis) for each WTA AOI (dots) in each donor (*x* axis). **c.** Deconvolution highlights differences in cell composition between PanCK^+^ and PanCK^-^ alveolar AOIs and between AOIs from SARS-CoV-2 (D22-24) and SARS-CoV-2 positive (all others) lung samples. Expression scores (color bar) of cell type signatures (rows) in PanCK^+^ (left) and PanCK^-^ (right) alveolar AOIs (columns) in WTA data from different donors (top color bar). **d,e.** Changes in gene expression in SARS-CoV-2 positive *vs.* negative lung. Left: Significance (-Log_10_(P-value), *y* axis) and magnitude (log_2_(fold-change), *x* axis) of differential expression of each gene (points) in WTA data between SARS-CoV-2 positive and negative AOIs for PanCK^+^ (**d**) and PanCK^-^ (**e**) alveoli. PanCK^+^ alveoli ROIs: 78 SARS-CoV-2 positive *vs.* 18 negative; PanCK^-^ alveoli ROIs: 112 positive *vs.* 20 negative. The horizontal dashed line indicates an FDR q-value cutoff of 0.05, and the two vertical dashed lines represent a fold-change of 2 in log_2_ scale. The names of top 10 SARS-CoV-2 positive and negative significant genes regarding fold-change are marked, respectively. Right: Significance (-log_10_(q-value)) of enrichment (permutation test) of different pathways (rows).

We retained high quality data for all kept AOIs, observing high sequence saturation (Supplemental Fig. 10b**, Methods**), and agreement between the CTA and WTA assays as reflected by correlation of expression levels for their overlapping genes (Supplemental Fig. 12a,b; cases with lower correlations are partly due to physical distance between serial sections; Supplemental Fig. 12c), viral expression scores (Fig. 5b, Supplemental Fig. 10c), inferred cell type composition (Fig. 5c, Supplemental Fig. 12d), and differential expression between COVID-19 and healthy samples (Fig 5d,e, Supplemental Fig. 12e,f, Supplemental Tables 7,8). We focused our subsequent analyses on the WTA data.

### Spatial data reveals compartment-specific expression phenotypes

Across samples D8-D12 and D18-D24 (D13-D17 were analyzed separately, below), the donor autopsies showed variation in SARS-CoV-2 RNA expression, with four donors demonstrating elevated levels (Fig. 5b, Supplemental Fig. 10c, **Methods**), consistent with the corresponding SARS-CoV-2 expression levels for the three samples also profiled by viral qPCR and sc/snRNA-Seq, and with the highest SARS-CoV-2 Spike signal in D20 by NanoString Protein assay (Supplemental Fig. 10c). Deconvolution of major cell type composition with gene signatures from our snRNA-Seq atlas (**Supplemental Tables 9, 10, Methods**) showed distinct composition of PanCK^+^ and PanCK^-^ compartments of alveolar AOIs (Fig. 5c**, Methods**), such that PanCK^+^ compartments were dominated by inferred AT1 and AT2 cells, with smaller contributions of endothelial, ciliated, and basal cells. The two donors with normal alveolar morphology (D23 and D21) showed expression of both AT1 and AT2 cell markers, while most other donors showed a preference for either AT1 or AT2 cells in the PanCK^+^ compartment. Compared to the PanCK^+^ compartment, PanCK^-^ compartment had greater inferred cellular diversity across and within subjects, with strong expression of genes associated with fibroblasts, myofibroblasts, vascular endothelial, and lymphatic endothelial cells. Within the set of PanCK^-^ AOIs, AOIs from COVID-19 autopsy samples showed increased scores for fibroblasts and myofibroblasts compared to AOIs from control autopsies.

In comparing gene expression between COVID-19 and control alveolar AOIs (Fig. 5d,e and **Supplemental Table 7**), we observed that interferon-ɑ and γ response genes and oxidative phosphorylation pathways were upregulated in COVID-19 samples, whereas TNF alpha signaling via NFKB, IL2-STAT5 signaling, TGFB signaling, apical junction and hypoxia were all downregulated. This occurred in both PanCK^+^ and PanCK^-^ alveolar AOIs. Several genes upregulated in highly infected bulk RNA-Seq samples (*IFIT1, IFIT3, IDO1, GZMB, LAG3, NKG7, PRF1*) and in SARS-CoV-2 RNA containing myeloid cells (*TNFAIP6, CXCL11, CCL8, ISG15, GBP5*) were also upregulated in the PanCK^-^ alveolar compartment from COVID-19 samples compared to controls. The decreased TNF alpha signaling observed in the PanCK^+^ alveolar compartment contrasts with the increased TNF signaling found in SARS-CoV-2 RNA containing epithelial cells, which could be explained by differential signaling in SARS-CoV-2 positive vs. negative epithelial cells. In addition, 111 of 565 genes up-regulated in COVID-19 PanCK^+^ alveolar AOIs overlapped with genes expressed at higher levels in epithelial cell types with high *ACE2* expression^41^ (Supplemental Fig. 10e **and Supplemental Table 11**).

AOIs characterized as bronchial epithelium, artery, and alveoli in donors (D13-D17) exhibited distinct transcriptional programs (Supplemental Fig. 11b). Specifically, 8,958 genes were differentially expressed (FDR < 0.05, **Methods**) between bronchial epithelial AOIs and matched normal alveoli from the same subjects, with cilium assembly being the top most enriched pathway (Komogorov-Smirnov statistic FDR q-value = 2.25*10^-25^). Furthermore, significant transcriptional differences were also identified between inflamed and uninflamed alveoli AOIs within the same group of samples (D13-D17). A total of 1,246 genes were differentially expressed between inflamed and normal-appearing alveolar AOIs in the donors, with increased expression of innate immune and inflammatory pathway genes by ssGSEA^53, 54^ (**Methods**), including neutrophil degranulation (FDR q-value = 5.2*10^-17^), interferon-γ signaling (FDR q-value = 3.4*10^-15^), and signaling by interleukins (FDR q-value = 1.4*10^-13^). Notably, contrasting transcriptional activity was observed in AOIs adjacent to each other, as demonstrated by the example of upregulated interferon-γ pathway in inflamed alveoli when comparing to normal-appearing alveoli within the same biopsy sample (Supplemental Fig. 11c), with dysregulation of numerous pathway genes (Supplemental Fig. 11d).

Finally, comparing expression differences within a donor in the D8-12, D18-24 sample subset between SARS-CoV-2 high and SARS-CoV-2 low AOIs (defined by their SARS-CoV-2 virus signature scores, Supplemental Fig. 13a,b**; Methods**), we found not only the expected induction of the viral *ORF1ab* and *S* genes, but also upregulation in the PanCK^+^ compartment of the chemokines *CXCL2* and *CXCL3*, and of the immediate early genes *EGR1, JUN, FOS, IER2, ZBTB10,* and *NR4A1*, consistent with our comparison of SARS-CoV-2^+^ and SARS-CoV-2^-^ epithelial cells by snRNA-Seq (**Supplemental Table 6**, Supplemental Fig. 8g). Interestingly, *NT5C*, encoding a nucleotidase with a preference for 5’-dNTPs, is consistently up-regulated in SARS-CoV-2 high in both PanCK^+^ and PanCK^-^ compartments (Supplemental Fig. 13c,d, **Supplemental Table 12**). There are no reports of this gene playing a role in lung injury, and it will be the target of future investigation.

### Impact of severe COVID-19 on heart, kidney, and liver

To gain insight as to how COVID-19 might affect other tissue types, we profiled liver (n = 16), heart (n = 15) and kidney (n = 11) tissues from a subset of the donors by snRNA-Seq (**Methods**), and constructed integrated atlases for each tissue spanning 36,662 heart, 47,001 liver and 29,568 kidney nuclei (Fig. 6). In all three tissues, we recovered the major parenchymal, endothelial, and immune cell subsets, with both automated and manual annotations (Fig. 6a,c,d,f,g,i). Cell classes were generally well-mixed among donors and SARS-CoV-2 entry factors (*ACE2*, *TMPRSS2*, *CTSL*) were expressed in the expected cell classes; for example, *ACE2* is highly expressed in kidney proximal tubular cell subsets and in heart cardiomyocytes and pericytes (Fig. 6j, Supplemental Fig. 14a,h-k), and *ACE2* and *TMPRSS2* are co-expressed in liver cholangiocytes (Fig. 6k).

**Figure 6.**
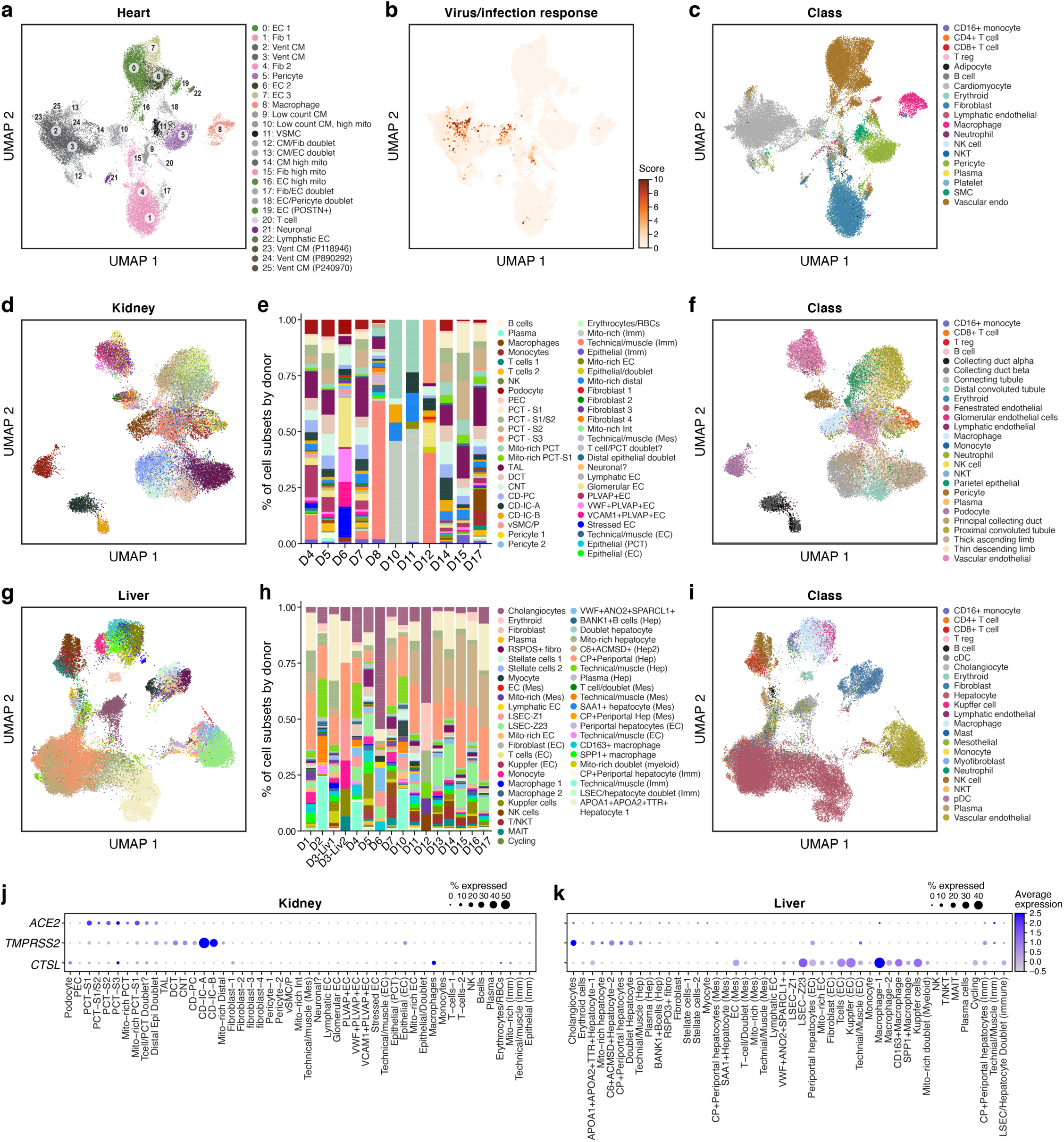
A single nucleus atlas of heart, kidney, and liver COVID-19 tissues. **a-c.** COVID-19 heart cell atlas. UMAP embedding of 36,662 heart nuclei (dots) from 15 samples, colored by clustering with manual *post hoc* annotations (**a**), signature scores of genes upregulated in SARS-CoV-2 infected iPSC cardiomyocytes^55^ (**b**), labeling mostly cardiomyocytes from patient D17 (see also Supplemental Fig. 10b-g), or by automatically derived cell type labels (**c**). **d-f.** COVID-19 kidney cell atlas. **d,f.** UMAP embedding of 29,568 kidney nuclei (dots) from 11 samples, colored by clustering with manual *post hoc* annotations (**d**) or by automatically derived cell type labels (**f**). **e.** Proportion of cells (*y* axis) in each subset (color legend, as in **d**) in each donor (*x* axis). **g-i.** COVID-19 liver cell atlas. **g,i.** UMAP embedding of 47,001 liver nuclei (dots) from 16 samples, colored by clustering with manual *post hoc* annotations (**g**) or by automatically derived cell type labels (**i**). **h.** Proportion of cells (*y* axis) in each subset (color legend, as in **g**) in each donor (*x* axis). **j,k.** SARS-CoV-2 entry factors are expressed in kidney and liver cells. Average expression (dot color) and fraction of expressing cells (color, size) of SARS-CoV-2 entry factors (rows) across cell subsets (columns) in the kidney (**j**) and liver (**k**).

Notably, we detected very few viral RNA reads in all three tissues, and almost all in droplets that would be deemed technical artifacts, such as misalignments (Supplemental Fig. 15). In one heart sample, the absence of viral reads was also confirmed by NanoString DSP and RNAscope (*data not shown*). Though no viral counts were present in the heart nuclei that passed quality control, we further examined the cardiomyocytes for evidence of gene expression changes similar to those observed in SARS-CoV-2 infected iPSC-derived cardiomyocytes *in vitro*^55^ (Fig. 3e). Scoring each nucleus by the expression of gene sets upregulated in infected vs. uninfected iPSC-derived cardiomyocytes revealed elevated expression among select cardiomyocytes (Fig. 6b). The high-scoring cells are predominantly from a single individual, D17, who was diagnosed with cardiac complications (Supplemental Fig. 14b-g).

### Specific cell types in lung, liver and kidney are associated with severe COVID-19 through integrated GWAS risk loci

A complementary view of the mechanisms and causes of severe COVID-19 is offered by genetic studies, based on either common genetic variation studied through GWAS^56^, or on rare, coding loss-of-function variants^57, 58^. While these studies identify loci that causally contribute to disease risk, mapping the specific genes, their mechanisms, and organ/cell of action is challenging, especially because these may only manifest in the disease context. We reasoned that integration of the common variant GWAS data with our rich atlas can help address some of these challenges, because both the COVID-19 (C2: COVID-19 versus population) and severe COVID-19 (B2: hospitalized COVID-19 versus the population) phenotypes show evidence of significantly non-zero heritability (B2: N.cases=8,638, N.controls= 1,736,547, h^2^ (liability scale): 0.06, Z-score (h^2^) = 4.8, and C2: N.cases= 30,937, N.controls= 1,471,815, h^2^ (liability scale): 0.01, Z-score (h^2^) = 2.7).

To study genetic variants associated with severe COVID-19, we integrated our atlas with the COVID-19 GWAS summary statistics from COVID-19 Host Genetics Initiative’s meta-analysis^59^ (round 4 freeze, Oct 20, 2020) for two phenotypes: severe COVID-19 and COVID-19. We formed a list of 27 curated genes proximal to the 6 regions with the genome-wide significant association signals thus far (chromosomes 3p, 3q, 9, 12, 19 and 21 loci (https://www.covid19hg.org/results/), **Supplemental Table 13**), but excluded *ABO* from the reported results, because its expression cannot be reliably reported in sc/snRNA-Seq, given red blood cell contamination. We also identified 64 genes with aggregated GWAS signals using MAGMA^60^ based upon two SNP-to-gene (S2G) linking strategies (**Methods**), which included 17 of the 25 curated GWAS genes (**Supplemental Table 13**). (We expect curated GWAS genes to overlap with MAGMA, as SNP associations lead to both GWAS loci and MAGMA genes.) Finally, we analyzed the 25 curated GWAS genes and 61 MAGMA implicated genes separately in each of the four tissue atlases (lung, liver, kidney, and heart).

The 25 curated GWAS genes across the four tissue atlases showed both cell type specific expression and differential expression between cells of the same cell type in COVID-19 vs. healthy tissue. In the lung, 21 of the 25 curated GWAS genes showed significant (FDR < 0.05) expression specificity in at least one cell type, including *DPP9* (chr 19) in lung AT2 cells and *CCR1* and *CCRL2* (chr 3) in lung macrophages (Fig. 7a **and Supplemental Table 14**). Furthermore, 18 of the 25 curated GWAS genes showed significant (FDR < 0.05) differential expression between cells of the same type in COVID-19 *vs*. healthy lung, including *SLC6A20* in goblet cells, *CCR5* in CD8 T cells and Tregs, and *CCR1* in macrophages and CD16 monocytes (Fig. 7b **and Supplemental Table 14**). For 55% of these GWAS genes, the average expression was significantly higher in the lung (p value < 0.05, t-test) compared to heart, liver, or kidney (Supplemental Fig. 16). A similar specificity in cell type expression in the lung was also observed for the extended list of 61 MAGMA implicated genes (**Supplemental Table 14**).

**Figure 7.**
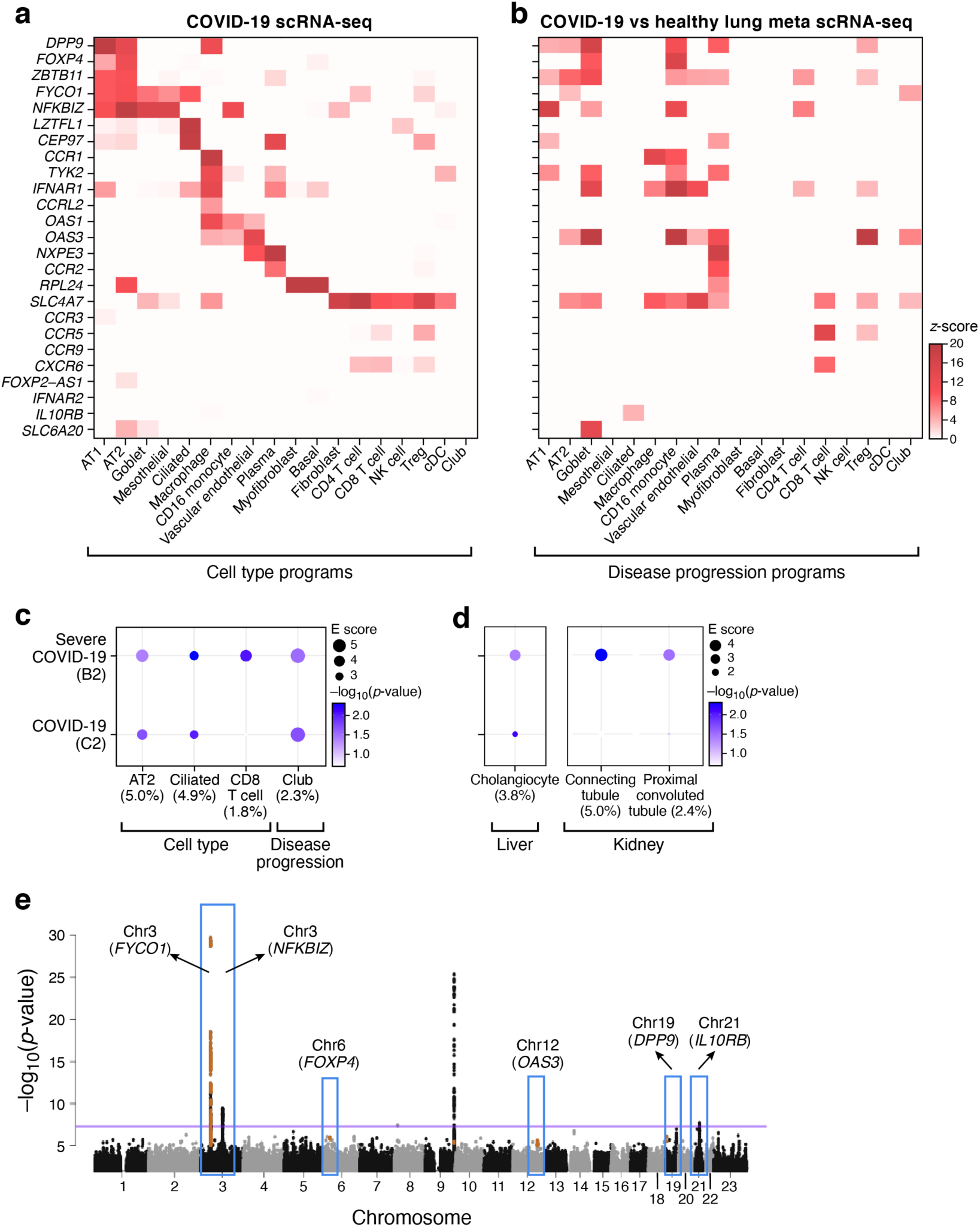
Integration of COVID-19 GWAS with COVID-19 lung, liver, and kidney single cell profiles helps nominate genes and cell types in severe disease. **a,b.** Relation of genes from GWAS-associated regions to cell type specific expression. Mean expression (**a**, z-score relative to all other cell types, color bar) or differential expression (**b**, z-score of DE analysis of expression in COVID-19 *vs.* healthy cells of the same type) of 25 genes (rows) from 6 genomic loci associated with COVID-19 (based on summary statistics data from COVID-19 HGI meta analysis^59^ across lung cell types (columns). **c**. Cell types and gene programs in the lung that contribute to heritability of COVID-19 severity by incorporating genome wide signals from GWAS. Magnitude (circle size, E score) and significance (color, −log_10_(P-value)) of the enrichment of cell type programs and cell-types specific disease programs (columns) that were significantly enriched for COVID-19 or severe COVID-19 phenotypes (rows). **d**. Extending the analysis in (c) to cell types and gene programs in liver and kidney with significant heritability enrichment signal for COVID-19 severity. All results in (**c**) and (**d**) are conditional on 86 baseline-LDv2.1 model annotations. **e**. Nomination of single best candidate genes at unresolved GWAS significant loci by aggregating gene level information across program classes and cell types. Significance (-log_10_(P-value), *y* axis) of GWAS association signal at locus (*x* axis). Blue boxes: Significantly associated loci^59^ at a genome-wide significance level (purple horizontal bar). Nominated genes are labeled. Numerical results are reported in Supplemental Tables 14, 15 and 16.

To relate COVID-19 severity traits to cell type level processes in each tissue, we analyzed gene programs characterizing cell types in general, as well as disease associated programs that are differentially expressed in the same cell type between COVID-19 and healthy tissue (**Methods**). We used sc-linker, a new computational approach (**Methods**, KAJ, KKD, ALP, AR, *unpublished work*), to link gene programs defined by scRNA-Seq to genetic signal using linkage disequilibrium (LD) score regression^61–63^.

In the lung, three cell type programs attained nominal significance (but none attained Bonferroni-corrected significance, and should thus be interpreted cautiously): AT2 cells, ciliated cells and CD8 T cells (Fig. 7c **and Supplemental Table 15**). The CD8 T cell program showed the highest excess overlap with the curated GWAS genes (3.4x), followed by the AT2 cell program (1.8x). In other tissues (kidney, liver and heart), nominal (but not Bonferroni-corrected) significance was attained in the proximal convoluted tubule and connecting tubule cell type programs in kidney and cholangiocytes in liver (Fig. 7d **and Supplemental Table 15**). Moreover, disease progression programs in the lung highlighted significant association and high excess overlap in GWAS genes (3.8x) (Fig. 7c **and Supplemental Table 15**).

Finally, we nominated genes in each cell type (across programs) as potentially relevant for COVID-19 by integrating the sc-linker results with MAGMA gene level analysis (**Methods**). In lung, this approach highlighted, for example, OAS3 in AT2 and club cells and SLC4A7 in CD8 T cells (**Supplemental Table 16**). The highest number of nominated genes was observed for lung AT2 cells and spanned several loci, hinting at a polygenic architecture linking AT2 cells with COVID-19 (**Supplemental Table 16**). We further used this approach in the lung to prioritize genes by cell type specificity in COVID-19 lung at unresolved, significantly associated GWAS loci (chromosomes 3, 9, 12, 19 and 21) (Fig. 7e), prioritizing, for example, *FYCO1* (AT2, ciliated, club; chr3p), *NFKBIZ* (AT2; chr3q), and *DPP9* (AT2; chr 19) (**Supplemental Table 16**).

## Discussion

Here, we established a clinical sample processing pipeline to collect and profile autopsy tissues from COVID-19 donors at the single cell and spatial level, offering a rare opportunity to examine the impact of severe disease in the same tissue across multiple donors, and multiple tissues within a single donor. To ensure long term success for meta-analyses, we collected careful clinical annotations, harmonized the common descriptors, and coordinated lab protocols with a sister atlas collected in a New York City hospital (Melms *et al.,* companion manuscript). To overcome variable tissue quality, given differences in necrosis and PMI, we optimized our tissue processing protocols and implemented new computational strategies to analyze the resulting sc/snRNA-Seq data. Using CellBender, we removed ambient RNA and empty droplets from the data, and addressed ambient viral RNA, leading to more defined cell subsets with specific expression of canonical cell type markers, and allowing us to reliably identify cells harboring viral RNA. By implementing a novel approach to automatically annotate individual cells by their types, we generated annotated atlases of the lung, heart, kidney, and liver, and were able to leverage existing human data sets to contextualize our findings.

Our parallel expert manual annotation of cell clusters by prior knowledge and literature-based gene sets helped further characterize cell subsets that may be more specific to COVID-19 host responses, including those that may not yet have been observed in other disease atlases and thus would not be readily captured through automated annotation with reference datasets. For example, this allowed us to distinguish different subtypes of macrophages, expressing different scavenger receptors and showing varying rates of viral enrichment.

In particular, our analysis of different epithelial cell subsets and programs provides a detailed cellular landscape of the catastrophic set of events in these infected terminal lungs, showing how multiple avenues of repair have failed. First, we see relative depletion of AT1 and AT2 cells, the expected devastating epithelial damage caused by direct viral infection of those cells. Next, we observe an intermediate epithelial cell state (*KRT8*^+^ PATS/ADI/DATP) that was previously described to represent the transition from AT2 cells to AT1 cells during alveolar regeneration^37–,39^, especially in the context of lung fibrosis^46^, suggesting that the loss of AT1 cells has triggered AT2 cell transition, but possibly with an unsuccessful intermediate. Third, to our knowledge, we observe for the first time the transcriptional profile of IPBLP-like cells in human lungs, enabling their further characterization in the pathologies of COVID-19 and other lung diseases in which they may emerge. Overall, the presence of both PATS and IPBLP-like cells in COVID-19 lungs indicates that multiple regenerative strategies are invoked to re-establish the alveolar epithelial cells that are lost to viral infection. The loss of AT2 cells and accumulation of PATS that we observe in COVID-19 lungs indicates that AT2 self-renewal and AT2 to AT1 differentiation are compromised, likely because of ACE2-mediated infection of AT2 cells. We hypothesize that the loss of AT2 cells and their insufficient self-replacement may trigger the mobilization of IPBLP as a last-resort strategy to regenerate the alveolar epithelial barrier, though this may come at the cost of fully-functional alveolar structure and function^46^. Notably, both airway secretory and alveolar AT2 cells express *ACE2*, indicating the tropism of SARS-CoV-2 for major progenitor populations in the lung, and suggesting that the loss of these progenitors may further compound the devastating epithelial damage caused by direct viral infection. Thus, the path to the terminal disease state of these lung samples is evidenced by the serial failure of epithelial progenitors to enact regeneration at the rate that functional cells are lost, first by secretory progenitor cells in the nasal passages and large and small airways, then followed by alveolar AT2 cells, PATS, and IPBLP cells, and eventually leading to complete lung failure (Fig. 3f).

To allow characterization of possible tissue pathologies, we also report possible “doublets” (cell barcodes with transcripts that would reflect cells of two different types and may reflect two cells/nuclei that are co-encapsulated) or cells/nuclei with high mitochondrial reads that passed initial quality controls. Further work will be needed to validate whether “possible doublets” are indeed technical artifacts or a result of biological events, such as dying or virally infected cells being phagocytosed (*e.g.*, T+endothelial/macrophage cells). Meanwhile, cells with relatively high mitochondrial reads could similarly be biologically relevant, as both proliferating/metabolically active and necrotic cells tend to express higher levels of these genes. Relatedly, at the macroscopic level, tissue necrosis was evident and we observed evidence of proliferating fibroblasts at the gene expression level (Fig. 2b).

The level of viral RNA detected in lung samples varied significantly by donor, and there was a negative correlation between viral burden in lung tissue and time from symptom onset to time of death, likely reflecting different stages of COVID-19 lung pathology and viral infection. We primarily detected viral RNA within myeloid and endothelial cells (Fig. 4h-m). Genes differentially expressed between SARS-CoV-2 RNA+ and SARS-CoV-2 RNA-cells of the same type were enriched for chemokine and cytokine signaling, and responses to interferon, TNF, intracellular pathogens, and viruses. Surprisingly, however, we did not observe significant concordance between the expression of SARS-CoV-2 entry factors (*ACE2, CTSL, FURIN*, *TMPRSS2, PCSK7, PCSK5, PCSK2,* and *NRP1*, Fig. 4g) and SARS-CoV-2 RNA+ cells (Fig. 4h). Notably, in the seven samples containing highest levels of total SARS-CoV-2 UMI and the only SARS-CoV-2 RNA+ cells, we did not detect substantial populations of viable epithelial cells, including AT1, AT2, and ciliated subtypes compared to samples with lower total viral loads. In particular, within the four individuals with the overall highest viral load, epithelial cells accounted for less than 10% of the total cells captured (Supplemental Fig. 7k, 8c). This may be due to excessive epithelial death secondary to high viral replication within targeted pneumocytes and a highly inflammatory environment. We further caution that cell-associated SARS-CoV-2 UMI here may represent a mix of true replicating virus in targeted cells, immune cells engulfing virions or virally-infected cells, and cells with virions or virally-infected cells non-specifically strongly attached to their cell surface (the last option with higher likelihood in lungs overwhelmed by high viral burden, Fig. 5a). Moreover, as we rely on an snRNA-Seq procedure (albeit with associated ribosomes^30^), we may under-represent viral RNA. Spatial analysis of COVID-19 lung tissue supports the observation that high viral levels are mainly found at the earlier stages of infection^64^, as we observed upregulation of the immediate early genes (*EGR1*, *JUN*, *FOS*, *IER2*, and *NR4A1*) only in SARS-CoV-2 high epithelial regions of the lung (Supplemental Fig. 9). While SARS-CoV-2 has been previously detected in the heart, liver and kidney, we did not detect viral reads by sc/snRNA-Seq in these tissues in the cohort in our study.

By necessity, our study is largely descriptive, due to a small sample size (n=17), but by combining our single-cell profiles with signals from GWAS of COVID-19 and severe COVID-19, we can begin to shed light on the relationship between genetic risk factors and tissue physiology. Analyzing both 27 genes from regions that have genome-wide statistically significant associations as well as genome wide association signals from the COVID-19 summary statistics, our atlas helps highlight likely functionally relevant genes in unsolved GWAS loci by their expression in relevant cells, especially in the diseased lungs, as well as highlight cell types that are implicated by overall genetic signal. Both analyses showed more prominent signal in lung cells than in kidney, heart, or liver, especially implicating genes in AT2 cells, ciliated cells, and CD8+ T cells at the cell type level, and AT2 cells and macrophages based on disease specific gene expression, and helped highlight specific genes from multi gene regions, such as *FYCO1*, *SLC4A7* and *NFKBIZ* in chromosome 3, as underlying the association to those cells. Proximal convoluted tubule and connecting tubule cells in the kidney, as well as cholangiocytes in the liver were also implicated in this analysis. Nevertheless, given the size of our cohort and limited power of the current GWAS, our results should be interpreted with care. As the GWAS grows, and as meta-analyses of multiple atlases can be done (for example, by integration with Melms *et al*., companion manuscript), such analyses can help better characterize the mechanisms of COVID-19 severe disease.

The data presented here can be used to inform future studies of COVID-19 tissue pathology and pathophysiology. The tissue processing and computational protocols we developed should be useful when studying an array of diseased or damaged tissues. In future studies, we will complete the profiling and release of the remaining seven tissue types collected, such as brain, spleen and trachea, to obtain a more complete view of COVID-19 derived organ pathology, and will integrate across atlases in meta-analyses, to provide critical resources for the community studying host-SARS-CoV-2 biology.

## Methods

### Human donor samples

Samples in this study underwent IRB review and approval at the institutions where they were originally collected. Specifically, Dana-Farber Cancer Institute approved protocol 13-416, Partners/Massachusetts General Hospital and Brigham and Women’s Hospital approved protocols: 2020P000804, 2020P000849, and 2015P002215; Beth Israel Deaconess Medical Center approved protocol 2020P000406, 2020P000418. All external sample cohorts were then added to the Broad Institute’s protocol 1603505962 and reviewed and approved by the MIT Institutional Review Board (IRB). No subject recruitment or ascertainment was performed as part of the Broad protocol.

### COVID-19 Autopsy Study Sample Processing

#### Tissue collection

Organs from COVID-19 donors were collected at Boston area hospitals (MGH, BWH, and BIDMC). Exclusion criteria included a postmortem interval > 24 hours and HIV infection. Donor ages ranged from 30-35 to > 89 years of age (Fig. 1, **Supplemental Table 1**).

All patients at the Massachusetts General Hospital (MGH) who succumbed from SARS-CoV-2 infection, as confirmed by the qRT-PCR assays performed on nasopharyngeal swab specimens, were eligible for clinical autopsy upon consent by their healthcare proxy or next of kin. A subset of these patients were also enrolled in the MGH Rapid Autopsy Protocol if they had a history of known or suspected malignancy. Their clinical data and research specimens were collected in accordance with Dana Farber/Harvard Cancer Center Institutional Review Board-approved protocol 13-416. Autopsies at MGH were performed in a negative pressure isolation room by personnel equipped with powered air-purifying or N95 respirators. Organs were removed from the body *en bloc*, and subsequently dissected for individual organ examination, including weighing and photographing. Representative samples of each lung lobe, left ventricle of the heart, liver, kidney, and brain (for donor D6) were placed in collection tubes, which were then placed inside a cooler for immediate transport to the Broad institute where all tissues were processed fresh.

Autopsies at Brigham and Women’s Hospital were performed in a negative pressure isolation room by personnel equipped with powered air-purifying or N95 respirators. Organs were removed from the body *en bloc*, and subsequently dissected for individual organ examination, including weighing and photographing. Representative samples of each lung lobe, trachea, left ventricle of the heart, aorta, coronary arteries, liver, kidney, lymph nodes, and spleen, as well as nasal and oral swab/scrapings were placed in 25 mL of RPMI-1640 media with 25 mM HEPES and L-glutamine (ThermoFisher Scientific) + 10% heat inactivated FBS (ThermoFisher Scientific) in 50 ml falcon tubes (VWR International Ltd). Collection tubes containing tissue were then placed inside a cooler for immediate transport to the Broad Institute. Additional tissue specimens from lung and heart were fixed in 10% formalin, processed and paraffin embedded using standard protocols. 5 µm-thick slides were prepared from the FFPE tissue blocks and transferred to the Broad Institute.

Autopsies at Beth Israel Deaconess Medical Center were performed on a cohort of 5 patients with positive SARS-CoV-2 nasopharyngeal swab on admission with a minimally invasive approach. Consent for autopsy and research was obtained from the healthcare proxy or the next of kin. Ultrasound-guided biopsies were performed within 3 hours of death on a gurney in the hospital morgue. The body was not removed from the bag until staff was wearing standard precautions and an N95 mask. Biopsies were obtained through 13G coaxial guide with 14G core biopsy and 20 mm sample length. Multiple core biopsies were acquired from each organ by tilting the coaxial needle a few degrees in different directions^65^. Cores for snRNA-Seq and spatial transcriptomics

#### Tissue processing

Tissues from deceased COVID-19 donors were processed immediately upon arrival in a Biosafety Level (BSL) 3 lab space. Tissues were washed in cold PBS and dissected in sterile Petri dishes. Grossly necrotic tissue areas were excluded. As phenotypes varied across organs, representative biopsies were collected from each tissue sample. Biopsies (each approximately 0.5 cm^3^) were cut and either directly dissociated for scRNA-Seq or frozen for future processing (each described below).

#### Biopsy collection for snRNA-Seq and scRNA-Seq

For snRNA-Seq, three dry biopsies per tissue were placed in one cryovial and flash frozen on dry ice before being placed at −80°C for long term storage. Two to four flash frozen tubes were collected per tissue.

For scRNA-Seq of viably frozen tissue, tissue was viably frozen by placing 4 biopsies in 500 ml of CryoStor CS10 freezing media (STEMCELL Technologies). Tubes were inverted and then placed on ice for up to 30 minutes. Samples were then stored at −80°C for future processing.

For scRNA-Seq of fresh tissue (*i.e.*, samples processed on the day of collection), single biopsies were placed in a dissociation buffer and processed as described below.

#### Fresh lung tissue processing and cryopreserved tissue processing for scRNA-Seq

For each sample, 1 mL of lung dissociation medium (100 µg/mL of DNAse I, Roche; 1.25 mg/mL of Pronase, Roche; 9.2 µg/mL of Elastase, Worthington; 100 µg/mL of Dispase II, Roche; 1.5 mg/mL of Collagenase A, Roche; 100 µg/mL of Collagenase IV, Worthington) was aliquoted in a 15 mL Falcon tube (VWR International Ltd) and kept on ice until use. For each tissue, 2 wells of a 24 well plate were prepared and contained 2 mL RPMI-1640 media with 25 mM HEPES and L-glutamine (ThermoFisher Scientific) + 10% heat inactivated FBS (ThermoFisher Scientific).

For viably frozen tissue, before sample thawing, tubes with lung dissociation media and the 24 well plate containing media were placed at 37°C to warm. One biopsy piece was transferred to one well of a 24 well tissue culture plate (Corning) and moved back and forth in the liquid to wash. The biopsy was then placed in a second well containing media for 1 minute before moving to a 1.5 mL Eppendorf tube containing warmed lung dissociation media. While the tube was in a rack, sterile scissors were used to mince the tissue until most pieces were < 1 mm. The Eppendorf tubes containing minced tissues were then placed in a thermomixer (Eppendorf) and incubated for 25 minutes at 300 RPM, 37°C. After 10-15 minutes, the sample was briefly triturated with a P1000 pipette, and returned to the thermomixer. After 25 minutes, 600 µL cold RPMI-1640 + 10% FBS + 1 mM EDTA (Invitrogen) was added to quench the digestion. Using a wide bore P1000 pipette tip (Rainin), the tissue was triturated∼ 20 times. The dissociated tissue was then placed on ice. One 50 mL conical per tissue preparation was placed in a rack and on ice, and a 100 µm filter was prepared by wetting with 1 mL cold RPMI-1640 + 10% FBS + 1 mM EDTA. Using a wide-bore P1000 pipette tip, digested tissue was passed through the filter. The filter was washed with 4 mL cold RPMI-1640 + 10% FBS. Tubes were spun at 300g x 8 minutes at 4°C. Supernatant was removed and pellet was resuspended in 2 mL ACK lysis buffer, placed on ice for 2-3 minutes. The lysis was quenched in 2 mL RPMI-1640 + 10% FBS, and the sample was spun at 300g x 8 minutes at 4°C. After centrifugation, most of the supernatant was removed and the sample pellet was resuspended in ∼1 mL of RPMI-1640 + 10% FBS. Cells were counted and immediately loaded on the 10x Chromium controller (10x Genomics) for partitioning single cells into droplets.

#### Flash-frozen tissue processing for snRNA-Seq

All sample handling steps were performed on ice. TST and ST buffers were prepared fresh as previously described^30, 66^. A 2x stock of salt-Tris solution (ST buffer) containing 292 mM NaCl (ThermoFisher Scientific), 20 mM Tris-HCl pH 7.5 (ThermoFisher Scientific), 2 mM CaCl_2_ (VWR International Ltd) and 42 mM MgCl_2_ (Sigma Aldrich) in ultrapure water was made and used to prepare 1xST and TST. TST was then prepared with 1 mL of 2x ST buffer, 60 µL of 1% Tween-20 (Sigma Aldrich), 10 µL of 2% BSA (New England Biolabs), 930 µL of nuclease-free water and supplemented with 0.2 U/ul RNaseIN Plus (Promega) and 0.1U/ul Superasin (ThermoFisher Scientific). 1xST buffer was prepared by diluting 2x ST with ultrapure water (ThermoFisher Scientific) in a ratio of 1:1, and was supplemented with 0.1% RNase inh (Enzymatics). 1 mL of 1xST without RNase was also prepared for sample dilution prior to 10X chip loading. Single frozen biopsy pieces were kept on dry ice until immediately prior to dissociation. With clean forceps, a single frozen biopsy was placed into a gentleMACS C tube on ice (Miltenyi Biotec) containing 2 mL of TST buffer. gentleMACS C tubes were then placed on the gentleMACS Dissociator (Miltenyi Biotec) and tissue was homogenized by running the program “m_spleen_01” one to three times, until tissue was fully dissociated. A 40 µm filter (VWR International Ltd) was placed on a 50 mL falcon tube (VWR International Ltd) and pre-wet with 1 mL of 1xST. Homogenized tissue was then transferred to the 40 µm filter and washed with 2 mL of 1xST buffer containing RNase inhibitors. Samples were then centrifuged at 500g for 10 minutes at 4°C. Sample supernatant was removed and the pellet was resuspended in 100 µL −500 µl 1xST buffer (without RNase inhibitor). Nuclei were counted and immediately loaded on the 10x Chromium controller (10x Genomics) for single nucleus partitioning into droplets.

#### Single-cell and single-nucleus RNA library preparation and sequencing

For each sample, 8,000-15,000 cells or nuclei were loaded in one channel of a Chromium Chip (10x genomics). 3’ v3 chemistry was used to profile lung nuclei. 5’v1.1 chemistry was used to profile freshly dissociated lung cells, as well as cells obtained from cryopreserved lung tissue. 3’ v3.1 chemistry was used to process all other tissues. cDNA and gene expression libraries were generated according to the manufacturer’s instructions (10x Genomics). cDNA and gene expression library fragment sizes were assessed with a DNA High Sensitivity Bioanalyzer Chip (Agilent). cDNA and gene expression libraries were quantified using the Qubit dsDNA High Sensitivity assay kit (ThermoFisher Scientific). Gene expression libraries were multiplexed and sequenced on the Nextseq 500 (Illumina) with a 150 cycle kit and the following read structure: Read 1: 28 cycles, Read 2: 96 cycles, Index Read 1: 8 cycles, or on the Novaseq (S2 flow cell; Illumina) with a 100 cycle kit and the following read structure: Read 1: 28 cycles, Read 2: 91 cycles, Index Read 1: 8 cycles.

#### Bulk RNA-Seq of lung tissue

With clean forceps, a single frozen biopsy was placed into a gentleMACS M tube (Miltenyi Biotec) containing 600 µL of 2x DNA/RNA shield (Zymo Research). gentleMACS M tubes were then placed on the gentleMACS Dissociator (Miltenyi Biotec) and tissue was homogenized by running the program “RNA_02”. Samples were centrifuged at 2,000xg for 1 minute and moved to new tubes. As DNA/RNA shield inactivates SARS-CoV-2, the samples could then be moved out of the BSL3 lab space. RNA was extracted using the quick-RNA MagBead kit (Zymo Research). RNA was treated with proteinase K according to manufacturer’s instructions. For some donors (D13-D17), we had access to a single frozen lung biopsy. Bulk nuclei were isolated as a suspension (as described above) and flash frozen in RLT + 1 % 1 BME. To isolate total RNA, we added ∼ 600 µl of 2x DNA/RNA shield. After proteinase K treatment, one volume of water was added to dilute the DNA/RNA shield. Total RNA was quantified via Nanodrop (ThermoFisher Scientific).

cDNA and gene expression libraries were constructed with Smart-Seq2 chemistry and the Nextera XT DNA library prep kit (Illumina), as previously described^67, 68^. DNA and gene expression library fragment sizes were assessed with a DNA High Sensitivity Bioanalyzer Chip (Agilent). cDNA and gene expression libraries were quantified using the Qubit dsDNA High Sensitivity assay kit (ThermoFisher Scientific). Gene expression libraries were multiplexed and sequenced on the Nextseq 500 (Illumina) with a 75 cycle kit and the following read structure: Read 1: 38 cycles, Read 2: 38cycles, Index Read 1: 8 cycles, Index Read 2: 8 cycles

#### Bulk RNA library preparation and sequencing of lung tissue for viral genome assembly

Metagenomic sequencing libraries were constructed for viral genomic analysis. Briefly, 5 µL of each RNA sample underwent RNAse H-based ribosomal RNA depletion^69^, then libraries were constructed using an Illumina TruSeq Stranded Total RNA Library Construction kit (20020596). This ligation-based library construction method utilized adapters (1.5 uM each) that contained a unique molecular identifier on the upstream of the i7^70^. Some libraries with lower viral loads were selected for targeted enrichment of SARS-CoV-2 sequences using a probe set designed to capture sequences from several respiratory pathogens, including SARS-CoV-2; this design was adapted from a published method^71^. Targeted enrichment was performed using a Twist Bioscience Target Enrichment workflow (https://www.twistbioscience.com/sites/default/files/resources/2019-11/Protocol_NGS_HybridizationTE_31OCT19_Rev1.pdf). All libraries were pooled and sequenced to a read depth of at least 2 million reads on an Illumina MiSeq V3 and a NovaSeq SP with the read structure: read 1: 146 cycles; read 2: 146 cycles; index read 1: 17 cycles; index read 2: 8 cycles.

#### Viral quantification by RT-qPCR

For each bulk RNA sample, SARS-CoV-2 RNA was quantified by an RT-qPCR assay targeting the nucleocapsid protein (modified from the CDC N1 assay, as described previously^72^). The assay was performed on samples (diluted 1:3) in triplicate, along with negative controls and a synthetic DNA standard curve. Reported quantifications, calculated from the standard curve, are mean values across triplicates, and are corrected by the dilution factor.

#### Sample processing for RNAscope® and DSP analysis

FFPE blocks were cut onto Superfrost Plus microscope slides at the BWH Histopathology Core and shipped to the Broad Institute to be stored at −80°C.

For the RNAscope® assay, staining was performed according to the manufacturer protocol using the RNAscope® Multiplex Fluorescent Reagent Kit v2 kit (ACD Bio, 323100) with the V-nCoV2019-S-C3 probe (ACD Bio, 848561-C3) and the TSA Plus Cyanine 5 fluorophore (Akoya Biosciences, NEL745001KT). For samples D13-17, serial sections were fluorescently stained with RNAscope Probes - V-nCoV2019-S –C3 d, *ACE2* and *TMPRSS2* (ACDBio) and GeoMx Nuclear Stain (NanoString Technologies, GMX-MORPH-NUC-12) in blue. Additionally, for the same samples, serial sections were DAB stained with 2 µg/mL of anti-SARS-CoV-2 spike glycoprotein antibody (Abcam, ab272504). Slides were scanned at 20x directly on the NanoString GeoMx instrument to facilitate alignment with subsequent WTA/CTA/Protein scans. For subjects D13-17, a semi-quantitative (5-tier) viral abundance score (Viral Score) was used based on RNAScope SARS-CoV-2 staining to identify ROIs exhibiting diverse viral loads. The score assignment per AOI was performed by board-certified pathologists.

For the NanoString GeoMx DSP RNA assays, slides were prepared following the Manual RNA Slide Preparation Protocol in the GeoMx DSP Slide Preparation User Manual (NanoString, MAN-10087-03 for software v1.4). Briefly, slides were baked at 60°C for 45 minutes. Deparaffinization and rehydration were performed in xylene for 3×5 minutes, 100% ethanol for 2×5 minutes, 95% ethanol for 1×5 minutes and once in 1x PBS (Sigma Aldrich, P5368-10PAK) for 1 minute. Antigen retrieval was performed in 1x Tris EDTA pH 9.0 (Sigma Aldrich, SRE0063) in a Tinto Retriever Pressure Cooker (BioSB, BSB 7008) for 20 minutes at 100°C. Thereafter, the slides were washed 1x PBS for 5 minutes. Tissue RNA targets were exposed by incubating the slides with 1mg/ml proteinase K (Ambion, 2546) in 1x PBS at 37°C for 15 minutes. Slides were washed in 1x PBS for 5 minutes, then immediately placed in 10% Neutral Buffered Formalin (EMS Diasum, 15740-04) for 5 minutes, followed by incubation in NBF Stop Buffer (1.48M Tris Base, 563mM Glycine) for 2×5 minutes, and once in 1x PBS for 5 minutes. *In situ* hybridizations with the Cancer Transcriptome Assay (CTA) RNA probes or Whole Transcriptome Assay (WTA) were performed at 4 nM final probe concentration in Buffer R (NanoString). Both probe sets were spiked with a panel of COVID-19 RNA plus probes (NanoString) at a final concentration of 4 nM. One at a time, slides were dried of excess 1x PBS, set in a hybridization chamber lined with Kimwipes wetted with 2x SSC, and covered with 200 µL prepared Probe Hybridization solution. HybriSlips (Grace Biolabs, 714022) were gently applied to each slide and they were left to incubate at 37°C overnight. The HybriSlips were removed by dipping the slides in 2x SSC (Sigma Aldrich, S6639)/0.1% Tween20. To remove unbound probes the slides were washed twice in 50% Formamide (ThermoFisher, AM9342)/2x SSC at 37°C for 25 minutes, followed by two washes in 2x SSC for 2 minutes. Slides were blocked in 200 µL Buffer W (NanoString), placed in a humidity chamber and incubated at room temperature for 30 minutes. Morphology marker solution was prepared for 4 slides at a time: 22 µL SYTO13 (NanoString), 5.5 µL Pan-cytokeratin morphology marker (NanoString), 7.5 µL CD45 morphology marker (NanoString), and 187 µL Buffer W for a total volume of 220 µL/slide. One at a time, slides were dried of excess Buffer W, set in a humidity chamber, covered with 200 µL morphology marker solution, and left to incubate at room temperature for 2 hours. Slides were washed twice with 2x SSC for 5 minutes and were immediately loaded onto the NanoString DSP instrument.

For the NanoString GeoMx DSP Protein Assay, slides were prepared following the Manual FFPE Protein Slide Preparation Protocol in the GeoMx DSP Slide Preparation User Manual (NanoString, MAN-10087-03 for software v1.4). Briefly, slides were baked at 60°C for 45 minutes. Slides were deparaffinized in xylene for 3×5 minutes and rehydrated in 100% ethanol for 2×5 minutes, 95% ethanol for 2×5 minutes and twice in DEPC water for 5 minutes. Antigen retrieval was performed with 1x Citrate Buffer pH 6.0 (Vector Laboratories, H-3300-250) in a Tinto Retriever Pressure Cooker (BioSB, BSB 7008) for 15 minutes at high pressure and high temperature. Pressure was released and the slides were removed to room temperature for 25 minutes. Slides were then washed twice in 1x TBS-T (Cell Signaling Technologies, 9997S). Hydrophobic barriers were drawn around each tissue section and the tissue blocked with 200 µL Buffer W (NanoString) for 1 hour at room temperature in a dark humidity chamber. Antibody solution consisting of eight protein panels was prepared by mixing 8 µL of each panel per slide of: Immune Cell Profiling Panel Core (NanoString), IO Drug Target Panel (NanoString), Immune Activation Status Panel (Nanostring), Immune Cell Typing Panel (NanoString), Cell Death Panel (NanoString), MAPK Signaling Panel (NanoString), PI3K/AKT Signaling Panel (NanoString) and COVID-19 Custom Panel (Abcam), 5 µL Pancytokeratin morphology marker (NanoString), 7 µL CD45 morphology marker (NanoString), and 123.5 µL Buffer W for a total volume of 200 µL/slide. Slides were dried of excess Buffer W and incubated with 200 µL antibody solution in a dark humidity chamber at 4°C overnight. Slides were washed three times in 1x TBS/0.1% Tween-20 for 10 minutes before being incubated with 200 µL 4% paraformaldehyde (Electron Microscopy Sciences, 50-980-487) in a dark humidity chamber at room temperature for 30 minutes. Nuclear stain solution was prepared for 4 slides at a time: 20 µLSYTO13 (Nanostring) and 180 µL 1x TBS (Cell Signaling Technologies, 12498S) for a total volume of 200 µL/slide. Slides were washed twice in 1x TBS-T for 5 minutes before incubation with nuclear stain solution in a dark humidity chamber at room temperature for 15 minutes. Slides were washed two times in 1x TBS-T and were loaded onto the DSP. Slides that were not immediately loaded onto the DSP were mounted with FluoromountG mounting media (SouthernBiotech, 0100-01), coverslipped, and stored in a dark humidity chamber at 4°C until they were ready to be processed. When ready, slides were soaked in 1x TBS-T for 30 minutes or until the coverslip fell off, and washed once in 1X TBS-T for 5 minutes before being loaded onto the DSP.

#### GeoMx DSP instrument and ROI selection

Slides were loaded into the slide holder of the GeoMx DSP machine and secured by closing the slide tray clamp. Each slide was covered with 2 ml of buffer S and the underside of the slides cleaned with 70% ethanol before loading the slide tray into the DSP. The scanning areas were set to only scan the tissues and the tissue boundaries were determined by adjusting the x and y scanning areas and the sensitivity. Each slide was scanned with a 20x objective and scan parameters: 100 ms FITC/525 nm, 300 ms Cy3/568 nm, 300 ms Texas Red/615 nm. 12-25 rectangular 700 µm x 800 µm region of interests (ROIs) were placed based on selection and assessment by a pathologist per tissue. When applicable, a segmentation mask was applied to ROIs selecting PanCK^+^ regions and SYTO13^+^ areas of interest (AOI). Each channel threshold for segments was manually adjusted to maximize inclusion of the AOI signal areas and minimize non-signal areas. In addition, a general setting of erode 1-2 µm, N-dilate 2 µm, hole size 160 µm^2^ and particle size 50 was used. After approval of the ROIs, the GeoMx DSP photocleaves the UV cleavable barcoded linker of the bound RNA probes and collects the individual segmented areas into separate wells in the DSP collection plate.

#### GeoMx protein nCounter pooling and preparation

DSP collection sample plates were dried down by sealing the plates with aeroseal and placed at room temperature overnight. Wells were resuspended with 7 µL DEPC-treated water, incubated for 10 min at room temperature and briefly spun down. Probe working pools were made corresponding to the number of rows of DSP aspirate in each collection. COVID-19 Custom Probe R was diluted 1:5 in 10 µL DEPC-treated water to achieve a working concentration. For full DSP collection plates, the Probe R working pool was prepared by mixing 8 µL of the following Probe Rs corresponding to the antibody panels used above: Immune Cell Profiling Panel Core, IO Drug Target Panel, Immune Activation Status Panel, Immune Cell Typing Panel, Cell Death Panel, MAPK Signaling Panel, PI3K/AKT Signaling Panel, COVID-19 Panel (1:5 working concentration) and 2 µL of DEPC-treated water for a total volume of 66 µL. The Probe U working pool was prepared by mixing 8 µL Probe U master stock and 58 µL DEPC-treated water for a total volume of 66 µL. For collections filling 2-3 rows of the 96-well DSP collection plate and requiring 2-3 Hyb Codes, the Probe R working pool was prepared by mixing 4 µL of the following Probe Rs corresponding to the antibody panels used above: Immune Cell Profiling Panel Core, IO Drug Target Panel, Immune Activation Status Panel, Immune Cell Typing Panel, Cell Death Panel, MAPK Signaling Panel, PI3K/AKT Signaling Panel, COVID-19 Panel (1:5 working concentration) and 1 µL of DEPC-treated water for a total volume of 33 µL. The Probe U working pool was prepared by mixing 4 µL Probe U master stock and 29 µL DEPC-treated water for a total volume of 33 µL. The final probe/buffer pool was prepared for the number of rows of DSP aspirate in each collection and a corresponding number of Hyb Codes required. For collections filling 8 rows of the 96-well DSP collection plate and requiring 8 Hyb Codes, the probe/buffer pool was prepared by mixing 64 µL Probe R working pool, 64 µL Probe U working pool, and 640 µLHybridization Buffer for a final volume of 768 µl. For collections filling 2 rows of the 96-well DSP collection plate and requiring 2 Hyb Codes, the probe/buffer pool was prepared by mixing 16 µL Probe R working pool, 16 µL Probe U working pool, and 160 µl Hybridization Buffer for a final volume of 192 µL. From these pools, 84 µL were aliquoted to each of the required Hyb Code tubes. 8 µL of each Hyb Code was distributed across a new hybridization 96-well plate row wise and 7µL DSP aspirates from the collection plate added to the corresponding wells in the hybridization plate. Hybridization plates were heat sealed, briefly spun down and loaded onto a thermal cycler at 67°C with the lid at 75°C for 16-24 hrs. Hybridization plates were cooled on ice and briefly spun down before pooling.

The amount of hybridized product from each plate to pool together was determined based on the total segment area (µm^2^) per column from the Lab Worksheet file produced by the DSP instrument following the volumes listed in Table 6 in the nCounter Readout User Manual (NanoString, MAN-10089-04 for software v2.0 JULY 2020). Hybridized products from each well were pooled column-wise into a 12-tube strip, briefly spun down, and loaded onto the nCounter MAX/FLEX Prep Station, run on the “High Sensitivity” setting. The prepared sample cartridge was then loaded onto the nCounter Digital Analyzer (NanoString) along with its respective Cartridge Definition File (CDF) from the DSP instrument, which contains vital indexing information, and was run with the setting provided by the CDF file. Once finished, the sample cartridge was wrapped in aluminum foil and stored at 4°C. The RCC file output by the Digital Analyzer was loaded onto the DSP instrument for analysis.

#### GeoMx RNA Illumina library preparation

DSP collection sample plates were dried down by sealing the plates with aeroseal and placed at room temperature overnight. Following resuspension of the wells with 10 µL of DEPC-treated water the plates were incubated for 10 min at room temperature before briefly spun down. 96-well PCR plates were prepared by mixing 2 µL PCR mix (NanoString), 4 µL of index primer mix (NanoString) and 4 µL of DSP sample. The following PCR program was used to amplify the Illumina sequencing compatible libraries; 37°C for 30 min, 50°C for 10 min, 95°C for 3 min, followed by 18 cycles of (95°C for 15 sec, 65°C for 1 min, 68°C for 30 sec), 68°C for 5 min and a final hold at 4°C. The indexed libraries were pooled with a 8-channel pipette by combining 4 µL per well from the 12 columns into 8-well strip tubes. The combined 48 µl pools in the 8-well strip were incubated with 58 µL AMPure XP beads (Beckman Coulter) (1.2x bead to sample ratio) for 5 min. The 8-well strip was placed on a magnetic stand for 5 min before removal of the supernatant. The beads were then washed twice with 200 µl of 80% ethanol and dried for 1 min before being eluted with 10 µL elution buffer (10mM Tris-HCl pH 8, 0.05% Tween-20). The 8 samples were combined into two pools of 40 µL each and incubated with 48 µL AMPure buffer (2.5M NaCl, 20% PEG 8000) for 5 min. Following magnetic incubation for 5 min and removal of supernatants the beads were washed twice with 200 µL of 80% ethanol and dried for 1 min before eluted with 17 µL elution buffer. The two pools were combined to a final elute of 34 µL. Sequencing library size was assessed with a DNA high sensitivity bioanalyzer assay (Agilent Technologies) and the expected library size of 161 bp was observed. Library concentration was assessed with Qubit dsDNA high sensitivity assay (Thermo Fisher Scientific).

Total target counts per DSP collection plate for sequencing were calculated from the total sampled areas (µm^2^) reported in the DSP generated Lab Worksheet. For CTA and WTA libraries, the target sequencing depths were 30 and 150 counts/µm^2^, respectively. Each library was diluted to 4-10 nM and combined to reach the estimated counts/µm^2^ per library in the final pool. All sequencing libraries were generated with unique indexes allowing pooled sequencing. We combined two of the CTA and five of the WTA libraries for sequencing with an Illumina NovaSeq S4 platform at a loading concentration of 365 pM with 5% PhiX. Two of the CTA collection plate sequencing libraries were sequenced with Illumina NextSeq500 HO 75 cycle kits at a loading concentration of 1.6 pM with 5% PhiX. The sequencing parameters used were: read 1, 27 cycles; read 2, 27 cycles; index 1, 8 cycles; index 2, 8 cycles.

### Computational methods

#### Sc/snRNA-Seq expression matrices

Cell Ranger mkfastq (10x Genomics) was used to demultiplex the raw sequencing reads, and Cell Ranger count on Terra using the cellranger_workflow in Cumulus^33^ was used to align sequencing reads and generate a counts matrix. Reads were aligned to a custom-built Human GRCh38 and SARS-CoV-2 RNA reference. ScRNA-Seq reads were aligned to “GRCh38_and_SARSCoV2”, and snRNA-Seq reads were aligned to “GRCh38_premrna_and_SARSCoV2’’. The GRCh38 pre-mRNA reference captures reads mapping to both exons or introns, and was built as previously described^66^. The SARS-CoV-2 viral sequence (FASTA file) and accompanying gene annotation and structure (GTF file) are as previously described^73^. The GTF file was edited to include only CDS regions, with added regions for the 5’ UTR (“SARSCoV2_5prime”), 3’ UTR (“SARSCoV2_3prime”), and anywhere within the Negative Strand (“SARSCoV2_NegStrand”) of SARS-CoV-2. Trailing A’s at the 3’ end of the virus were excluded from the SARSCoV2 fasta file, as we found these to drive spurious viral alignment in pre-COVID-19 samples.

#### Ambient RNA removal from expression matrices

CellBender remove-background^32^ was run (on Terra) to remove ambient RNA and other technical artifacts from the count matrices. The workflow is available publicly as cellbender/remove-background (snapshot 11) and documented on the CellBender github repository as v0.2.0: https://github.com/broadinstitute/CellBender.

This latest version of CellBender remove-background cleans up count matrices using a principled model of noise generation in scRNA-Seq. The parameters “expected-cells” and “total-droplets-included” were chosen for each dataset based on the total UMI per cell *vs*. cell barcode curve in accordance with CellBender documentation. Other inputs were left at their default values. The false positive rate parameter “fpr” was set to 0.01, 0.05, and 0.1. For downstream analyses we used the ‘FPR_0.01_filtered.h5’ file.

#### Quality control

Following CellBender, individual samples were processed using Cumulus, including filtering out cells/nuclei with fewer than 400 UMI, 200 genes, or greater than 20% of UMIs mapped to mitochondrial genes.

#### Sc/snRNA-Seq sample integration

Sc/sn RNA-Seq data from individual samples were combined into a single expression matrix and analyzed using Scanpy^74^ with identical pre-processing steps — aggregation, PCA, and batch correction using Harmony-Pytorch^34^. First, transcript counts were clipped and then normalized. UMI counts were clipped at 13, the value of the 99th percentile of non-zero counts. Clipped count data were then normalized so that UMI counts per cell/nucleus summed to 10,000 and then logged, resulting in log(1+10,000*UMIs / total UMIs) for each cell/nuclei, “logtp10k”.

Next, highly variable genes were identified by Scanpy’s highly_variable_genes function applied separately to data from each profiling modality: cryopreserved (cells), fresh tissue (cells), and fresh-frozen (nuclei), and retained genes that were highly variable across two or more methods. Next, data dimensionality was reduced to 75 top principal components by PCA, followed by Harmony^34, 75^ for batch correction, where each sample was considered a separate batch. Neighbors were computed using the Harmony corrected PCA coefficients and UMAP and Leiden clusters (below) were computed using the resulting nearest neighbors’ graph.

#### Clustering

Leiden clustering was performed on a *k*-nearest neighbor graph (*k*=100) with two different resolutions: 1.3 and 2. Resolution 1.3 was sufficient for coarse annotations of clusters. Clustering was performed using Pegasus^33^ with default parameters (except the resolution as mentioned above).

#### Automated cell annotation

Individual cell profiles were annotated by classification using a logistic regression model trained on six publicly available lung or bronchial alveolar lavage scRNA-Seq datasets with published manual labels, all profiled using the 10x Genomics platform. These datasets included healthy lung^35^ (and *unpublished*), pulmonary fibrosis^76, 77^, and COVID-19^78^ (and *unpublished*) samples. Two additional datasets were withheld from training to evaluate the predictive model^79, 80^. Since annotations between published datasets differed in resolution and nomenclature, we manually determined a single reference label set. We selected the most granular cell type labels that appeared in at least two datasets in the training set. Published cell type labels were mapped to that reference list or those cells were removed from training if they did not match a term.

To ensure consistent gene expression quantification across studies, training datasets were re-indexed to a single reference gene list and re-normalized from counts. Before training, data from all datasets were scaled together with the Scanpy scale function with zero_center=False.

An L2 penalized logistic regression model was trained using SciKit-Learn’s LogisticRegression function with C=0.1, max_iter=30, and multiclass=ovr. Performance on train and test sets was quantified by determining the frequency that predictions matched provided labels. If the predicted label was a cell type not included in the provided labels for that dataset, that cell was excluded from the accuracy calculation.

The learned model coefficients (**Supplemental Table 3**) represent the contribution of each of the 18,000 included genes to the prediction of all 28 cell types. These coefficients identify many known cell type markers. For example, the top 18 most important markers for Regulatory T cells include *CD3G*, *CD3D*, *CTLA4*, *ICOS*, *CD28*, *IL2RA* and *FOXP3*, the top 12 most predictive genes for AT2 cells are surfactant genes *SFTPA1*, *SFTPA2*, *SFTPB*, *SFTPC*, and *SFTPD*, and the top 5 smooth muscle cell genes are *ACTG2*, *ACTA2*, *CNN1*, *MYH11*, *DES*. While these genes are learned from training on other datasets, they are co-expressed within cell type specific modules in our lung samples.

Kidney, heart and liver were classified using a similar approach after ambient removal and basic quality control as described above. For classifying kidney nuclei, a published meta-atlas^81^ was leveraged as a training dataset. For liver and heart, we used publicly available 10x datasets for training (**Supplemental Table 2**).

#### Annotation by manual curation of lung

The manual annotation of clusters was done in two main steps: (1) Identifying main lineages in the global clustering, (2) Identifying sub population of cells by sub-clustering each lineage separately.

##### Identifying main lineages

Lung clusters were manually annotated by identifying elevated expression of pre-known markers in UMAP plots as well as visualizing mean Z-score expression of gene sets characterizing known populations (Supplemental Fig. 4b-4c). Immune cell types were identified using canonical markers (Supplemental Fig. 4b), whereas in order to identify the rest of the lineages we used both published^29, 82, 83^ and *unpublished* gene sets (Supplemental Fig. 4c, **Supplemental Table 17**).

##### Sub-clustering pre-processing

After identifying the main cell lineages, we studied each lineage by sub-clustering it separately from the rest of the cells, to enable identification of subtle cell states. For each of the three immune cell types and the endothelial cells we re-computed highly variable genes, performed PCA, batch corrected with Harmony, generated K neighbor neighbors (k=100), and performed Leiden clustering, all with Pegasus^33^. We tested different options for the number of highly variable genes and PCs, followed by manual inspection of the resulting UMAP, and choosing the parameters where cell clusters were evenly distributed and devoid of finger-like structure. We assessed the stability of clusters with rand-index^84, 85^, and chose the highest resolution that showed a rand_index value > 0.85.

The final parameters of number of highly-variable genes, number of PCs and resolution were: T and NK cells - (1500, 5, 0.7)

B and Plasma cells - (150, 4, 0.3)

Endothelial cells - (500, 15, 0.5)

Myeloid cells - (1500, 5, 0.9)

We tested each cluster for differentially expressed genes, and merged clusters that showed no difference. One cluster in the original T and NK sub-clustering was identified as a B cell subset and was reported in that lineage. Harmonization at the lineage level merged different samples evenly across clusters, but cells and nuclei data were separated in all the sub-clusterings, making the cell data merged in one cluster, with or without additional nuclei in the same cluster (Supplemental Fig. 4g).

##### Identifying unique differentially expressed genes per sub-cluster

Differentially expressed (DE) genes were determined for each final sub-cluster by two methods: (1) single cell differential analysis implemented in Pegasus, and (2) pseudo-bulk analysis, summing UMIs from all cells in each cluster per donor, and then performing DE analysis between clusters by fitting a linear regression using the functions “lmFit” and “eBayes” from the limma R package^86^ (Results from method 1 are available in **Supplemental Tables 18-21**).

To summarize the top DE genes in a heatmap (Supplemental Fig. 4d) we set different cutoffs to the statistics measured in each lineage to get the following number of DE genes per lineage: (a) the top 25 DEGs from method (1), (b) all DE genes above a threshold value of AUC from method (1), (c) all DE genes from with adjusted p-value < 0.05 and fold-change above a threshold. The threshold used for each lineage and the total number of DEGs found are as follows: Endothelial cells: 237 DE genes, minimum AUC=0.75, minimal log fold-change=3.8. T+NK cells: 229 DE genes, minimum AUC=0.8, minimal log fold-change=2.5. Myeloid: 263 DE genes, minimum AUC=0.75, minimal log fold-change=3.5. B+Plasma cells: 108 DE genes, minimum AUC=0.8, minimal log fold-change=2.

##### Sub-cluster annotations

Annotations of the unique sub-populations that were identified in the sub-clustering process were done by using legacy knowledge of these sub-populations through looking at unique gene markers associated with AUC statistics > 0.70 and pseudo-bulk DE over 2-fold difference in expression. Myeloid cells and B and plasma cells were split into states based on highly expressed genes in each sub-cluster. To annotate the endothelial cells we overlaid the 30 top DE genes of identified endothelial cells from a healthy lung cell atlas^35^ onto the UMAP of our endothelial data.

In the remaining four subpopulations, we did not detect known gene signatures that correspond to healthy human lung ECs, so we imputed the endothelial subpopulations using Adaptively-thresholded Low Rank Approximation (ALRA; implemented with Seurat v3.2.1) to compensate for EC signature dropout. After imputation, we were able to annotate the remaining EC subpopulations (Supplemental Fig. 4f): clusters 1-3. The remaining EC subpopulation (cluster 4) has a gene expression signature that overlaps with multiple EC subpopulations in the healthy lung reference data.

For the lung epithelial and fibroblast lineages, we harmonized the cells using LIGER^87^, a non-negative matrix factorization approach that allows cells to be described as occupying multiple different cell states. For the epithelial and fibroblast lineages, we used 8 factors and 7 factors respectively in the factorization. Each factor describes a different cell state, and to describe these states, we examined the top 100 genes loading each factor (**Supplemental Tables 22-23**). We identified the PATS cell state using the gene signature derived from alveolar organoids in Supplementary Table 1^37–39^.

Within the epithelial lineage, we further sub-clustered the KRT8/PATS/ADI/DATPs (cluster 7 in Fig. 2g) and airway epithelial cells (cluster 3 in Fig. 2g), and for each we applied LIGER with 4 factors. We identified intrapulmonary basal-like progenitor cells (IPBLPs) and airway basal cells by expression of the marker genes *TP63*, *KRT5*, and *KRT8*, and by testing for differentially expressed genes using the Mann-Whitney U test. To identify expression differences between IPBLPs and airway basal cells, we compared these two cell subsets using the same test.

#### Annotation by manual curation of heart, kidney, and liver

For heart, liver and kidney, we used standard approaches of graph-based clustering on the integrated datasets (Seurat v3 for kidney and liver; Scanpy for heart) to first derive high-level groupings followed by differential gene expression to determine cluster enriched genes. For kidney and liver, we defined the endothelial and immune compartments as predominantly *PECAM1+/CD31+* and *PTPRC+/CD45+* clusters respectively from the high-level assignment, and subjected them to iterative clustering, differential expression and post-hoc annotation to determine granular subsets. Similar iterative clustering was performed for the high-level mesenchymal groups in both tissues, proximal tubular compartment in the kidney and hepatocyte compartment in the liver. For heart, Leiden clustering at a resolution of 1.5 yielded clusters whose marker genes were compared to the heart atlas from Tucker *et al.*^88^. Where applicable, technical or putatively low-quality clusters are indicated as such to allow for permissive downstream analysis by the interested reader.

#### Doublet detection of snRNA-seq data for lung, heart, kidney and liver

We identified doublets using a two-step procedure for each tissue type based on ambient RNA removed gene count matrices. First, we independently identify doublets for each sample by calculating Scrublet-like^89^ doublet scores and then automatically determining a doublet score cutoff. Secondly, we integrated and clustered all samples from the sample tissue type and further marked any cluster that has more than 70% of cells identified as doublets as a doublet cluster. All cells within doublet clusters were marked as doublets.

We used a recent re-implementation of the Scrublet algorithm in Pegasus to calculate Scrublet-like doublet scores per sample. The original Scrublet algorithm consists of three major steps: preprocessing, doublet simulation and doublet score calculation using a k-nearest-neighbor (kNN) classifier. In the preprocessing step, we filtered low quality cells using the same criteria discussed in the quality control section, normalized each cell to TP100K, log-transformed expression values to log(TP100K+1) and identified highly variable genes using Pegasus^33^. We then standardized each highly variable gene based on the normalized count matrix (not log-transformed) and computed the first 30 principal components. In the doublet score calculation step, we built the kNN graph using Pegasus’ kNN building function.

We automatically determined a doublet score cutoff for each sample by log-transformed the Scrublet-like doublet scores for simulated doublets, estimated a smoothed density curve based on log-transformed doublet scores using kernel density estimation, identified an appropriate cutoff by inspecting the local maxima and signed curvature values of the density curve and finally transformed the cutoff from the log space by taking the exponential. Any cells with a doublet score larger than the cutoff were identified as doublets. For more details, please refer to documentation on gitub^90^.

The previous sample-level doublet detection procedure might miss some doublets. Inspired by work^91, 92^, we integrated quality-controlled samples for each tissue type, normalized each cell to TP100K and log transformed the expression value (log(TP100K+1), selected highly variable genes, computed the first 50 principal components (PC), corrected batch effects based on the PCs using Harmony-PyTorch, and clustered cells using the Louvain algorithm with a resolution of 1.3. We then tested if each cluster was statistically significantly enriched for doublets using Fisher’s exact test and controlling False Discovery Rate at 5%. Finally, among clusters that are significantly enriched for doublets, we picked those with more than 70% of cells identified as doublets and marked all cells in these clusters as doublets.

#### Deconvolution of bulk RNA-Seq lung samples

We performed composition estimation for the bulk lung samples using the MuSiC R package^93^. Ten bootstrap estimates, each using an independent sampling of 10,000 single cells, stratified by annotated cell type, to use as reference, were used to perform estimation.We confirmed that the subsampling of the reference to 10,000 cells was appropriate and did not introduce a specific sample composition bias by comparing the predicted sample composition with successively lower number of reference cells in the range of 10,000 to 5,000 cells (Supplemental Fig. 5d).

#### Healthy reference comparisons and differential expression

To test statistical significance of changes in cell type proportions between healthy and COVID-19 samples we used a Dirichlet-multinomial regression, as described by Smillie *et al*^94^.

We tested for differential expression between COVID-19 and other lung samples using Linear Regression models applied to each individual gene with the automated cell annotations. A model for each cell type was built using the “statsmodels’’ module in Python and included Boolean covariates for snRNA-Seq *vs*. scRNA-Seq and 10X V3 kit *vs.* other kits as well as a continuous covariate for UMI counts for each cell and a random intercept (Gene expression ∼ c1 + c2*(nuclei indicator) + c3*(10X v3 indicator) + c3*(number of UMIs) + c4*(COVID-19 indicator). Since each dataset had different numbers of cells, we sampled 2000 cells with replacement from the 10 healthy datasets and four COVID-19 datasets weighted specifically to balance the disease states and studies. We used Hold-Sidak multiple test correction in “statsmodels’’ with an error rate of alpha=0.05 to test which COVID-19 coefficients were predictive of gene expression levels.

#### Correction for ambient SARS-CoV-2 UMI

To address viral ambient RNA, we adapted several methods (CellBender^32^ and those described in Kotliar, Lin *et al*.^52^ and Cao *et al*.^95^) to create a conservative criteria for assigning a single cell or single nucleus as SARS-CoV-2 RNA+ or RNA-. We used a binomial test to determine the probability that the cell contained more SARS-CoV-2 UMIs than expected by ambient contamination, given the fractional abundance of SARS-CoV-2 aligning UMIs per cell, the abundance of SARS-CoV-2 aligning UMIs in the ambient pool, and the estimated percent ambient contamination of the single cell/nucleus. The fractional abundance of SARS-CoV-2 aligning UMIs per cell was defined as the number of UMIs to all viral genomic features (including the negative strand) divided by the total number of UMIs aligning to either SARS-CoV-2 or GRCh38 per cell. The abundance of SARS-CoV-2 aligning UMIs in the ambient pool was defined as the sum of all UMIs in the pre-CellBender output (from CellRanger counts) within discarded cells determined as “empty” or “low quality”, and was therefore determined on a per-sample basis. The estimated percent ambient contamination was defined as the ratio of the total UMIs for each single cell before *vs.* after CellBender. An exact binomial test was applied to each single cell using R given these parameters. P-values were adjusted using a Bonferroni correction and p < 0.01 was considered significant. Cells were assigned as “SARS-CoV-2 RNA+”, “SARS-CoV-2 Ambient” (if they had any UMIs to SARS-CoV-2 but were not significantly higher than the ambient pool) and “SARS-CoV-2 RNA-” (no UMIs to SARS-CoV-2).

#### Differential expression between SARS-CoV-2 RNA+ v*s.* RNA-cells

To test for host genes associated with the presence of SARS-CoV-2 RNA, we used the following approach to mitigate potential biases or confounding factors due to low cell numbers, differences in cell quality, or donor-to-donor variability: (**1**) We removed any cells classified as “SARS-CoV-2 ambient” from our analysis; (**2**) We analyzed DE genes only within cell types that contained at least 10 SARS-CoV-2 RNA+ cells; (**3**) Within a given cell type, we only considered donors where at least 2 cells were SARS-CoV-2 RNA+, and selected SARS-CoV-2 RNA-cells only from these donors; (**4**) We subsampled the SARS-CoV-2 RNA-cells to match complexity distributions to the SARS-CoV-2 RNA+ as previously described^41^ (briefly, cells were partitioned into 10 bins based on complexity (defined by the log_10_(# genes/cell)), and the SARS-CoV-2 RNA-cells were randomly subsampled to match the distribution in the SARS-CoV-2 RNA+ cells).

We tested differential expression using the DESeq2 package^96^, which fits a general linear model to each gene’s expression, and included % mitochondrial genes per cell as a continuous covariate, and the donor as a discrete covariate. Genes were considered DE if they had an FDR-corrected p-value < 0.05. Gene set enrichment testing with GSEA^53, 54^ was completed by defining a pre-ranked list of genes based on the log_2_(fold change) calculated from DESeq2 between SARS-CoV-2 RNA-cells and SARS-CoV-2 RNA+ cells of a given cell type. Hallmark, C2, and C5 gene sets were tested for enrichment, and an FDR-corrected p < 0.05 was considered significant.

#### Differential expression analysis of bulk RNA-Seq profiles from adjacent lung biopsies comparing “highly infected” and “uninfected” samples

Differential expression (DE) analysis was performed after FASTQ files were aligned to Hg19 using STAR ^97^. RSEM was used to generate a count matrix^98^ and DE genes were computed in R^99^ using DESeq2^96^. A log2 FC cutoff of 1.4 and an adjusted p-value of 0.05 was used as a threshold for significance. GSEA^53, 54^ was used to find gene ontology terms associated with the genes significantly up-regulated genes or down-regulated genes in highly infected tissue.

#### Viral Enrichment score

Inspired by Viral Track^100^, a program that identifies viral UMIs in human single cells and enumerates them per meta-cell, we computed a viral enrichment score per cluster. We could not employ the enrichment score of ViralTrack given the low number of SARS-CoV-2 RNA+ cells in our data, such that entire clusters, rather than higher resolution meta cells, were required for sufficient power.

The enrichment score for cluster C in a clustering of cells is computed as follows:

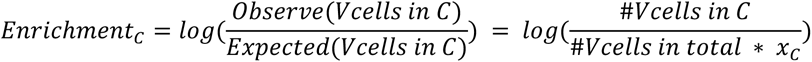

Where

*Vcells* are SARS-CoV-2 RNA+ cells

*X*_c_is the proportions of the total number of cells in cluster C out of the total number of cells in the given clustering (either all cells or within lineage).

We assessed the significance of each enrichment score empirically by permuting the data 100 times by randomly assigning the same number of SARS-CoV-2 RNA+ annotations to cells, such that the overall proportion of SARS-CoV-2 RNA+ cells was the same, computing the enrichment score per cluster, and calculating empirical p-value as the fraction of permutation that showed a higher enrichment score compared to the real result. We adjusted for multiple hypothesis testing by a Benjamini/Hochberg False Discovery Rate (FDR) as implemented in the function ‘fdrcorrection’ in the Python package statmodels^101^.

#### Generation of expression matrices from NanoString GeoMx CTA, WTA and protein data

For CTA and WTA (RNA) data, sequencing reads from NovaSeq or NextSeq was compiled into FASTQ files corresponding to each AOI using bcl2fastq. FASTQ files were demultiplexed and converted to Digital Count Conversion files using NanoString’s GeoMx NGS DnD Pipeline. The resulting DCC files were converted to an expression count matrix. AOIs (2 for CTA, 2 for WTA) were removed from the analysis if sequencing reads totaled less than 10,000. Target counts were generated by taking the geometric mean of all probe counts against the same gene target in each well. Individual CTA probes were dropped if they had counts less than 10% of the target counts (2 CTA) or if they were global Grubb’s outliers at alpha = 0.01 (6 for CTA). These probe screens were not performed on the WTA assay as each gene is sampled by one probe only. As a quality control step, sequencing saturation was calculated by dividing the number of non-unique sequencing reads by the total number of reads for each AOI. AOIs with sequence saturation below 50% were dropped from analysis (2 CTA, 0 WTA). The 75^th^ percentile of the target counts of each probe pool in each AOI were calculated and normalized to the geometric mean of the 75^th^ percentiles across all AOIs to give normalization factors. Target counts were divided by these normalization factors to give normalized expression counts.

WTA and COVID-19 Spike-in GeoMx DSP samples from D13-17 patients were processed similarly as above with a few modifications. Both WTA and COVID-19 Spike-in panels were sequenced on an Illumina NovaSeq platform, aligned to probe references and demultiplexed using Nanostring’s DnD pipeline (as above). The COVID-19 Spike-in panel contains multiple probes per target gene and counts that were not excluded by outlier detection were collapsed into a single value by calculating the geometric mean. In any cases where the probe counts were zero, the probe count was set to one prior to taking the geometric mean. Target counts for both the BIDMC’s WTA and COVID-19 Spike-in genes were then normalized by multiplying each count by its pool-specific negative probe normalization factor. Genes were further removed if their mean expression was greater than 100. Following these AOI-based and feature-based filtering steps, the data were normalized to the geometric mean of the 75^th^ percentile as above.

To measure DSP protein expression, indexing oligonucleotides were UV photocleaved from the profiled antibodies and deposited into a 96-well plate. Oligonucleotides were then hybridized to fluorescent barcodes and run through the nCounter MAX analysis system to generate raw digital counts. The raw counts were inputted into the DSP for calibration (adjusting for oligo-barcode binding efficiency) and for generation of a protein expression matrix; this matrix was subsequently normalized to internal spike-in positive controls (ERCC) to account for system variation. The calibrated and ERRC-normalized expression matrix was then normalized to the geometric mean of housekeeping genes^102^ as follows: first, the geometric mean of three housekeeping genes, histone *H6*, *GAPDH*, and *S6*, was calculated for each AOI; next, normalization factors were calculated as the geometric mean in each AOI dividing by the geometric mean of geometric means across all AOIs; finally, raw expression values were divided by the calculated normalization factors to give normalized protein expression levels.

Log-transformed matrices were obtained from normalized CTA, WTA and protein expression matrices by applying the function *f*(*x*) =*log* (*x* + 1) to each element of the matrices and were used for downstream analysis. We note that protein DSP data showed heterogeneity in immune cell marker levels across ROIs and donors (Supplemental Fig. 10d), but in many alveolar ROIs, PanCK and CD45 immunofluorescence stains showed significant mucus staining, which could affect antibody-based DSP data. Such off-target mucus binding was not observed in RNAscope images, and is thus less likely to be a concern in oligonucleotide probe-based WTA and CTA data.

#### Gene signature score calculation

A signature score summarizes the expression levels of a set of functionally-related genes for each AOI while controlling for gene abundance related technical noise^82^. Signature scores were calculated on the log-transformed expression matrices using Pegasus^33^’ new signature score calculation algorithm^103^. A gene set consisting of COVID-19 *S* and *ORF1ab* genes was used to calculate the SARS-CoV-2 virus signature score.

To score cell signatures we used a ‘reconciled’ annotation between the automated (‘predictions’) and manual annotations, as determined by the following rules. If a cell is assigned more than one label according to the rules, the first rule that applies to that cell determines its label. AT1: ‘leiden_res_2’ clusters number 10 and 26 (also consistent with ‘predictions’). AT2: ‘leiden_res_2’ clusters number 2 and 3 (consistent with ‘predictions’). B cell, plasma cell: Cells assigned as these types in ‘predictions’. CD4+ T cell: Cells predicted as either ‘CD4+ T cell’ or ‘Treg’ in ‘predictions’. CD8+ T cell: Cells predicted as ‘CD8+ T cell’ in ‘predictions’ and annotated as ‘T+NK’ in ‘manual_coarse_annotation’. NK cell: Cells predicted as ‘NK cell’ in ‘predictions’. Macrophage: Cells annotated as ‘Macrophages’ in ‘manual coarse annotation’. Mast cell: Cells annotated as ‘MAST’ in ‘manual coarse annotation’. Ciliated cell: Cells annotated as ‘Ciliated’ in ‘manual coarse annotation’. Secretory cell: Cells annotated as ‘Secretory’ in ‘manual coarse annotation’. Vascular endothelial cell: Cells predicted as ‘vascular endothelial’ in ‘predictions’. Lymphatic endothelial cell: Cells predicted as ‘lymphatic endothelial’ in ‘predictions’. Pericyte: Cells predicted as ‘pericyte’ in ‘predictions’. Smooth muscle cell: Cells predicted as ‘smc’ in ‘predictions’. Fibroblast: Cells predicted as ‘fibroblast’ in ‘predictions’. Myofibroblast: Cells predicted as ‘myofibroblast’ in ‘predictions’. Mesothelial cell: Cells predicted as ‘mesothelial’ in ‘predictions’.

#### Cell type specific markers for WTA and CTA deconvolution

To define cell markers for deconvolution, Pegasus v1.0 was used to conduct differential expression (DE) analysis on the snRNA-Seq data using the ‘reconciled’ annotation (above). For each cell type, Mann-Whitney U tests were conducted between cells annotated as this cell type and all other cells and with an FDR control at 0.05. DE genes were ranked by their Area Under the Receiver Operating Characteristics (AUROC) scores (**Supplemental Table 9**). Next, 10 marker genes were selected for each cell type by the combination of the DE results and published lung cell type markers^29, 41, 79^. We additionally picked 6 published marker genes for basal cells and neutrophils, respectively. These cell type markers were used for Fig. 5c and are available in **Supplemental Table 10**. Many of these markers were not available for the CTA data, which comprises only ∼1,800 genes. Thus, for cell types with less than 3 markers in the CTA data, we added genes based on the DE results to ensure at least 3 markers. These markers were used for Supplemental Fig. 12d and are available in **Supplemental Table 10.**

#### Differential expression analysis between subjects and controls for WTA and CTA data

All ROIs in WTA data annotated as alveoli PanCK^+^ and alveoli PanCK^-^ from donors D18, D19, D20, D21, D8, D9, D10, D11 and D12 were grouped together as the donor group (Fig. 5d,e); all ROIs in WTA data annotated as alveoli PanCK^+^ and alveoli PanCK^-^ from controls D22, D23 and D24 were grouped as the control group and then the limma package^86^ was used to perform differential expression analysis between the two groups (Fig. 5d,e). Specifically, Q3-normalized count data was transformed to log_2_-counts per million. The transformed data was fit with a linear model, considering the indicator functions of donor and control ROI, and empirical Bayes moderation was used to estimate the contrast S - C, where S stands for donor data and C for control. A similar approach is taken for alveoli CTA (Supplemental Fig. 12e,f).

#### Gene set enrichment analysis (GSEA) between subjects and controls for WTA and CTA data

GSEA analysis was conducted using fGSEA (R package)^104^ using the MSigDB Hallmark gene set (https://www.gsea-msigdb.org/gsea/msigdb/download_file.jsp?filePath=/msigdb/release/7.2/c2.cgp.v7.2.symbols.gmt), with gene ranks preset by their log_2_(fold-change) in a DE analysis, and the size of a gene set to test ranging between 15 and 500. Terms that are up-regulated (corresponds to COVID-19 samples) and down-regulated (corresponds to healthy samples) were plotted separately, with an FDR q-value threshold of 0.05.

#### CTA and WTA correlation analysis

Correlation analysis was performed across 1,783 genes that were shared between the CTA and WTA data and 306 ROIs where both CTA and WTA data were collected. For each ROI, the Pearson’s correlation coefficient between its corresponding CTA and WTA count vectors of the common genes. We excluded 10 ROIs where the WTA counts for the common genes were a constant vector of small numbers (resulting in NaN correlation).

#### Differential expression between SARS-CoV-2 high and low ROIs

WTA alveolar ROIs were first categorized by their SARS-CoV-2 virus signature scores (Supplemental Fig. 13a,b) as SARS-CoV-2-high (score>1.30), SARS-CoV-2-medium (0.75<=score<=1.30), and SARS-CoV-2-low (score<0.75). The limma package was used to perform differential expression analysis comparing SARS-CoV-2-high *vs.* SARS-CoV-2-low ROIs (separately for PanCK^+^ and PanCK^-^ AOIs), where Q3-normalized count data was transformed to log_2_-counts per million, the transformed data was fit into a linear model with indicator functions of SARS-CoV-2-high, SARS-CoV-2-medium, and SARS-CoV-2-low, respectively:

log2Y = b1 * I(X=SARS-CoV-2-high) + b2 * I(X=SARS-CoV-2-low) + b3 * I(X=SARS-CoV-2-medium),

where X is the ROI’s SARS-CoV-2 virus level, log2Y is the transformed count data on each gene, and I(X=·) is the binary indicator function on the corresponding SARS-CoV-2 virus level. Empirical Bayes moderation was then used to estimate the contrast SARS-CoV-2-high – SARS-CoV-2-low from this model.

#### Differential expression between inflamed and normal appearing alveoli

A total of 50 AOIs from lung biopsies (subjects D13-D17) were annotated as inflamed alveoli and 17 AOIs as normal alveoli within the same slides following review by board certified pathologists at the time of ROI selection (Supplemental Fig. 11a). Inflamed ROIs were selected based on histomorphology. Specifically, inflamed alveoli were defined as areas of alveolar tissue, excluding medium-sized bronchioles and large vessels, with intra-alveolar inflammatory cells (CD45^+^ or CD68^+^ cells) with or without fibroblastic foci. These areas generally also showed enlarged and detached type II pneumocytes and hyaline membrane formation. Non-inflamed ROIs (normal alveoli) were selected as areas of alveolar tissue from the same section. For reference, AOIs of bronchial epithelium (n=8) and of arterial vessels (n=5) were also selected. Q3-normalized count data were transformed to log_2_-counts per million and fit into a generalized linear model (Expression ∼ Patient + ROI Group). The patient-normalized residuals were utilized to generate plots following dimensionality reduction (UMAP, PCA). Differential expression analysis between inflamed (n=50) and normal appearing alveoli (n=17) was performed by incorporating the log_2_-counts per million into a generalized linear model and using contrasts to identify significant comparisons. All models were fitted using the quasi-likelihood negative binomial generalized log-linear model implemented in edgeR^105^.

#### Viral phylogenetic analysis

SARS-CoV-2 genomes were assembled using the open-source software viral-ngs version 2.0.21, implemented on the Terra platform (app.terra.bio), using workflows that are publicly available via the Dockstore Tool Registry Service (dockstore.org/organizations/BroadInstitute/collections/pgs). Briefly, reference-based assembly (assemble_refbased) was performed using the SARS-CoV-2 reference NC_045512.2. We constructed a phylogenetic maximum likelihood (ML) tree using the Augur pipeline (augur_with_assemblies) including unique genomes assembled with greater than 10X mean coverage depth and 772 previously reported genomes from unique individuals from Massachusetts^106^. We used Kraken2^107^ to search for the presence of 20 common respiratory viruses of interest (adenovirus, HCoV-229E, HCoV-HKU1, HCoV-NL63, betacoronavirus 1, parainfluenza 1, parainfluenza 2, parainfluenza 3, Parainfluenza 4, enterovirus A, enterovirus B, enterovirus C, enterovirus D, influenza A, influenza B, human metapneumovirus, respiratory syncytial virus, SARS-CoV, MERS-CoV, human rhinovirus) We used the classify_single and merge_metagenomics workflow with reference files as previously described^106^.

#### Integrated COVID-19 scRNA-Seq and GWAS analysis

Cell type gene programs were constructed from the scRNA-Seq data for each annotated cell type by performing a differential expression between the cell type on interest and the rest of the cells using a Wilcoxon rank sum test. Disease progression programs were constructed by a differential expression of disease and healthy cells of the same cell type as described in the “Healthy reference comparisons and differential expression” section earlier in the Methods. We also investigated gene programs based on genes that are differentially expressed in infected cells compared to uninfected, or genes that covary with the *ACE2* and *TMPRSS2* genes, but these programs showed no disease informative signal. Additionally, we performed MAGMA (de Leeuw 2015) on the COVID-19 summary statistics using both window-based and enhancer based SNP-to-gene linking strategies (union of Roadmap and Activity-By-Contact (ABC); RoadmapUABC).

#### Differential expression analysis in cardiomyocytes

Cardiomyocytes from the combined data (nuclei from Leiden resolution 1.5 clusters 2, 3, 12, 13, 14, 23, 24, and 25) were scored for the expression of genes upregulated in SARS-CoV-2 infected iPSC cardiomyocytes (*MYO1F, MYH13, MYH4, MYO1A, MYO7B, MYLK2, AMPD1, MYO15A, MYO1H, MYO3B, MYH15, MYH7, MYO3A, MYO7A, MYH2, OBSCN, MYL3, NEB, TNNT3, TRIM63, MYO1G, MYH8*) and genes downregulated in infected cells as compared to uninfected controls (*CALM1, ATP2A2, ENO3, ACTB, TPM2, TPM4, CALM3, TPM1, CALM2, MYL9, MYL6, TNNC1, CKM, MYL7, MYH11, MYLK, MYL4, TNNI1, MYH6, TPM3, TNNI3, CAPZB*) ^55^ using the “score_genes’’ function from Scanpy 1.5.1. A “virus/infection response score” was calculated as max(0, infected_score) - max(0, uninfected_score). The genes that contribute most to this score were assessed by conducting a differential expression test between high-scoring and low-scoring cardiomyocytes. Testing was performed by summing cardiomyocyte expression over individual and score-group (high: > 5, low: <=5) using voom and limma^86^ with the model “∼ 0 + scoregroup”, testing the contrast (scoregroup:high - scoregroup:low), and using Benjamini Hochberg to adjust p-values for multiple hypothesis testing.

#### Code availability

All samples were initially processed using Cumulus (https://github.com/klarman-cell-observatory/cumulus), which we ran on the Terra Cloud platform (https://app.terra.bio/). Code for all other analyses will be available on GitHub upon publication.

#### Data availability

Processed sequencing (sc/snRNA-Seq and bulk) data will be deposited in the GEO database (https://www.ncbi.nlm.nih.gov/geo/) and raw human sequencing data will be deposited in the controlled access repository DUOS (https://duos.broadinstitute.org/), upon publication.

Nanostring GeoMx raw and normalized count matrices will be available on GEO upon publication.

The processed lung data (sc/snRNA-Seq) is available on the Single Cell Portal: https://singlecell.broadinstitute.org/single_cell/study/SCP1052/

## Supporting information

Supplemental_Table_1_clinicalMetaData

Supplemental_Table_3_coefficients

Supplemental_Table_4_healthy_vs_covid

Supplemental_Table_6_SARS-CoV-2 RNA_posneg_DE

Supplemental_Table_7_WTA_Alveolar_spreadsheet

Supplemental_Table_8_CTA_Alveolar_spreadsheet

Supplemental_Table_9_broad_lung_reconciled.de

Supplemental_Table_10_cell_type_markers

Supplemental_Table_11_WTA_Alveolar_venn

Supplemental_Table_12_WTA_Alveolar_COVID_high_vs_low_DE_genes

Supplemental_Table_17_mannual_annotation_genesigs

Supplemental_Table_18_Myeloid_DE_results

Supplemental_Table_19_T_NK_DE_results

Supplemental_Table_20_B_Plasma_DE_results

Supplemental_Table_21_Endothelial_DE_results

Supplemental_Table_22_epithelial_sharedfactors_topgenes

Supplemental_Table_23_fibroblast_sharedfactors_topgenes

## Acknowledgements

We are deeply grateful to all donors and their families. We acknowledge the contribution of Casey Kania, Emmaline Kounaves, Nichole Lemelin, Justin Susterich, Jessica Teixeira, Claudia Bernal, Max Berstein, Allison Morris, Jordan N. Ray, Amanda Awley, Amanda Araujo and Erika Figueroa who all assisted in performing the autopsies at the Massachusetts General Hospital. We thank Ania Hupalowska and Leslie Gaffney for help with figure preparation. We thank Molly Veregge, Zachary Kramer, and Christopher Jacobs for their contributions in the execution of experimental procedures, and Dimitra Pouli who supported the creation of tissue annotation resources. We thank 10x Genomics, Illumina, BD Biosciences, and NanoString for help and support with instruments and/or lab reagents and technical advice. Portions of this research were conducted on the Ithaca High Performance Computing system, Department of Pathology, BIDMC, and the O2 High Performance Compute Cluster at Harvard Medical School. This project has been funded in part with funds from the Manton Foundation, Klarman Family Foundation, HHMI, the Chan Zuckerberg Initiative, and the Human Tumor Atlas Network trans-network projects SARDANA (Shared Repositories, Data, Analysis and Access). A.R. was an Investigator of the Howard Hughes Medical Institute. This project was also funded by DARPA grant HR0011-20-2-0040 (to D.E.I.) and the US Food and Drug Administration grant HHSF223201810172C (to P.C.S. and A.K.S.). A.-C.V. acknowledges funding support from the National Institute of Health Director’s New Innovator Award (DP2CA247831), the Massachusetts General Hospital (MGH) Transformative Scholar in Medicine Award, a COVID-19 Clinical Trials Pilot grant from the Executive Committee on Research at MGH, and the Damon Runyon-Rachleff Innovation Award. G.S. acknowledges support from the NIH R01AA0207440 and U01AA026933 research grants.

## Contributions

A.K.S., A.C.V., I.S.V., Z.G.J, O.R.-R., and A.R. conceived and led the study. T.M.D., C.G.K.Z., A. Segerstolpe, D.A. designed protocols and carried out experiments together with D.P, Z.B.-A., V.M.T., A.S, S.Z., J.G., J.H., E.N., M.S, C.M, L.T., A.E., D.Pa., L.P., and L.A.-Z. C.B.M.P.,C.G.K.Z., O.A., R.N.,G.H.,K.J.,K.S.,B.L.,Y.Y.,S.F.,A.S., P.N., Y.P.J., P.N., T.He., J.R., W.H., and I.S.V. designed and performed computational analysis, with input and assistance from M.B.,N.B. J.G., R.G., S.S.R., H.M., P.S, A.W., C.M., M. L.-G., T.H., D.T.M., S.W., D.Z., E.R., M.R, E.M., R.F.,P.D, A.G., C.Pe’er and M.N. K.J., K.D.,A.G., J.E., S.G., A.R and A.P., contributed methods and performed integrated analysis for GWAS. P.R.T., D.T.M., Z.G.J., Y.P., G.S., S.N., S.R., and J.R. provided clinical and biological expertise. D.F., D.J., D.E.M, C.P., S.K.V., E.K. and J.S. provided clinical expertise, performed sample acquisition and/or administrative coordination at MGH. J.H., R.R, R.N., O.R.B., Z.G.J., Y.P., and D.I. provided clinical expertise, performed sample acquisition and/or administrative coordination at BIDMC. I.H.S., D.A., L.C., J.G., R.P., M.S., provided clinical expertise, performed sample acquisition and/or administrative coordination at BWH. P.D, D.P, J.J-V, and J.B helped with sample coordination and sample receipt at the Broad Institute. N.B and S.R performed bulk RNA-Seq deconvolution analysis. E.N, M.R and K.S. performed viral qPCR, whole genome sequencing and phylogenetic analyses. M.S. provided input for sc/snRNA-Seq experiments and protocols. J.J.-V., E.T, O.R.R., and A.C.V managed the study and tissue acquisition. T.L.T contributed computational expertise and advice. T.M.D., C.B.M.P., C.G.K.Z., G.H., R.N., K.J., O.A., B.L., Z.G.J, I.S.V, Y.Y, S.F., A.S., D.T.M., A.K.S., A.C.V, O.R.-R. and A. Regev wrote the manuscript, with input from all authors. D.H., P.C.S., N.H., supervised research.

## Competing Interests

P.D., R.F., E.M.M., M.R., E.H.R., L.P., T.He., J.R., J.B., and S.W. are employees and stockholders at Nanostring Technologies Inc. D.Z., is a former employee and stockholder at NanoString Technologies. N.H., holds equity in BioNTech and Related Sciences. T.H.is an employee and stockholder of Prime Medicine as of Oct. 13, 2020. G.H. is an employee of Genentech as of Nov 16, 2020. R.N. is a founder, shareholder, and member of the board at Rhinostics Inc. A.R. is a co-founder and equity holder of Celsius Therapeutics, an equity holder in Immunitas, and was an SAB member of ThermoFisher Scientific, Syros Pharmaceuticals, Neogene Therapeutics and Asimov until July 31, 2020. From August 1, 2020, A.R. is an employee of Genentech. From October 19, 2020, O.R.-R is an employee of Genentech. P.C.S is a co-founder and shareholder of Sherlock Biosciences, and a Board member and shareholder of Danaher Corporation. A.K.S. reports compensation for consulting and/or SAB membership from Honeycomb Biotechnologies, Cellarity, Repertoire Immune Medicines, Ochre Bio, and Dahlia Biosciences. Z.G.J. reports grant support from Gilead Science, Pfizer, compensation for consulting from Olix Pharmaceuticals. Y.V.P. reports grant support from Enanta Pharmaceuticals, CymaBay Therapeutics, Morphic Therapeutic; consulting and/or SAB in Ambys Medicines, Morphic Therapeutics, Enveda Therapeutics, BridgeBio Pharma, as well as being an Editor – American Journal of Physiology-Gastrointestinal and Liver Physiology. GS reports consultant service in Alnylam Pharmaceuticals, Merck, Generon, Glympse Bio, Inc., Mayday Foundation, Novartis Pharmaceuticals, Quest Diagnostics, Surrozen, Terra Firma, Zomagen Bioscience, Pandion Therapeutics, Inc. Durect Corporation; royalty from UpToDate Inc., and Editor service in Hepatology Communications. P.R.T. receives consulting fees from Cellarity Inc., and Surrozen Inc., for work not related to this manuscript.

## Supplemental Figure Legends

**Supplemental Figure 1.**
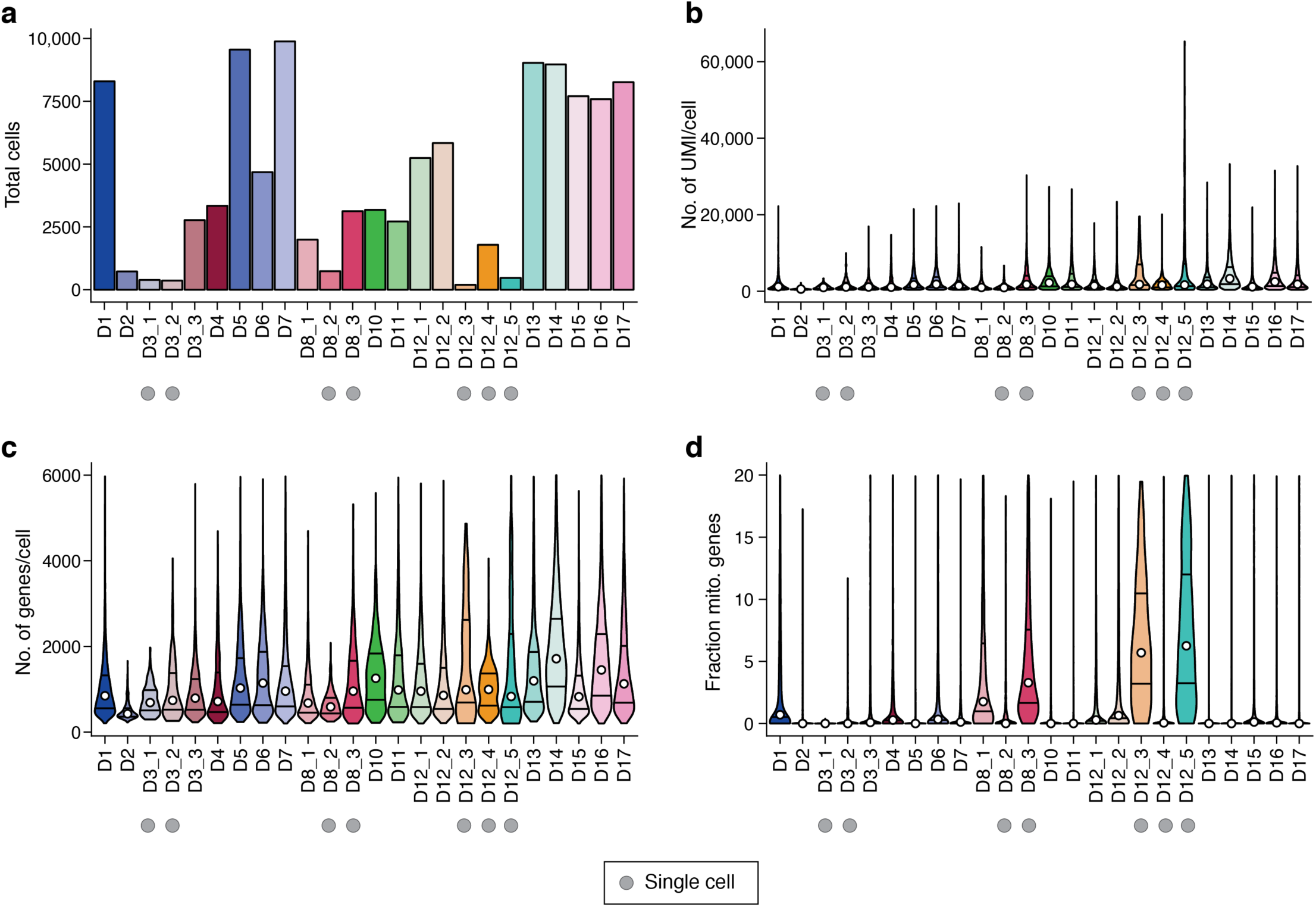
Quality control metrics for a single cell/nucleus lung atlas. Number of cells/nuclei (**a.**, *y* axis) and distributions of number of UMI per cell/nucleus (**b.**, *y* axis), number of genes per cell/nucleus (**c.**, *y* axis) and fraction of mitochondrial genes per cell/nucleus (**d.**, *y* axis) across the samples (*x* axis) in the lung atlas. ScRNA-Seq samples are labeled by a grey circle.

**Supplemental Figure 2.**
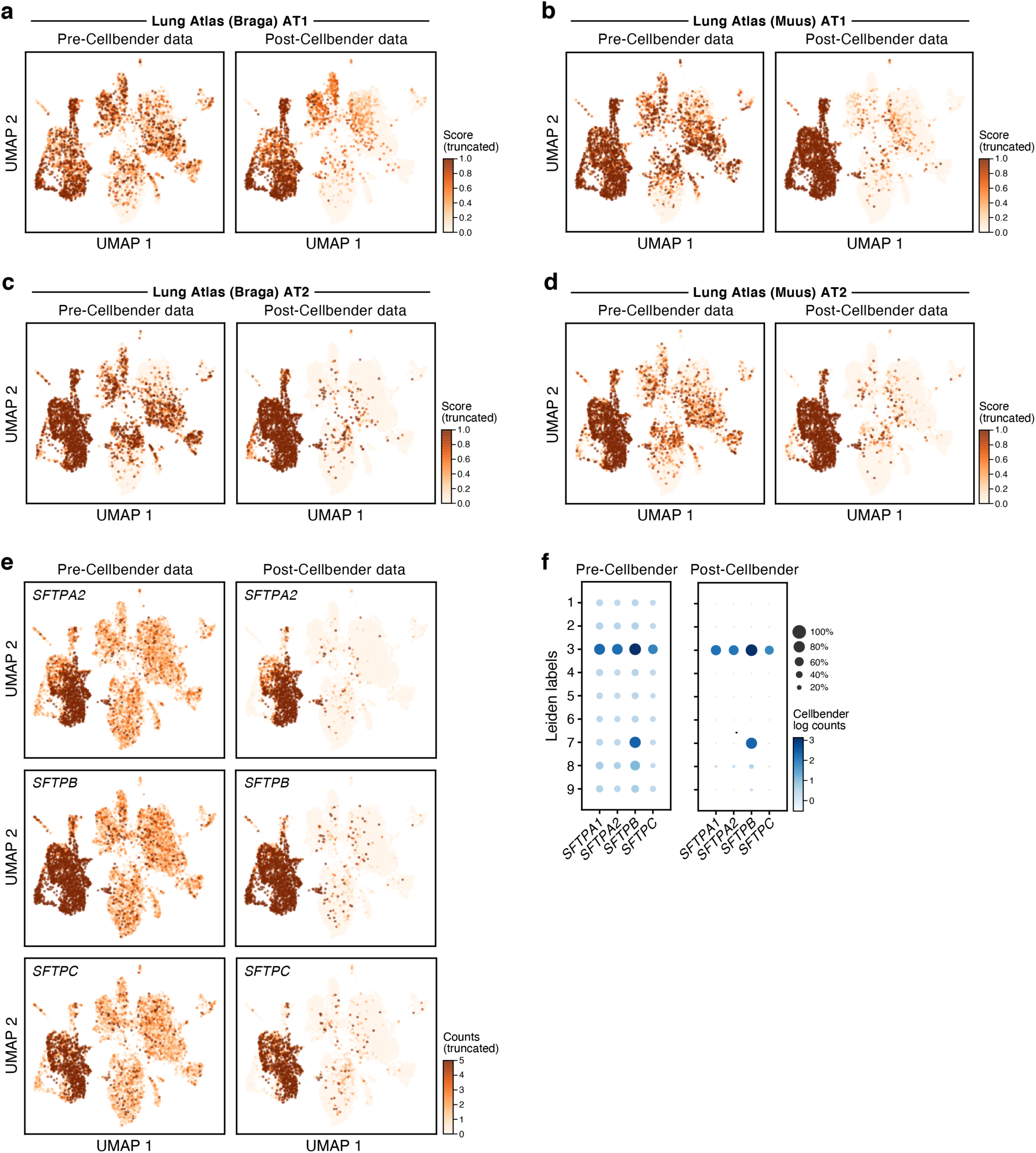
CellBender ‘remove-background’ improves expression specificity. **a-e.** CellBender improves specificity of individual genes and cell type signatures. UMAP embedding of single nucleus profiles pre-CellBender (left) and post-CellBender (right) processing from a single sample (D1), colored either by signature score (Scanpy’s^74^ score_genes function, color bar) for genes sets specific to lung (AT1 **a.**^79^ and **b.**^41^ and AT2 (**c.**^79^, **d**^41^), or by expression of surfactant protein transcripts (**e.** color bar, UMI counts). Color bar saturation chosen to emphasize low expression. **f.** Improved specificity of surfactant gene expression. Expression level (log(average UMI count per cell), color) and percent of cells with nonzero expression (dot size) of surfactant genes (columns) across cell clusters (rows) before (left) and after (right) CellBender processing.

**Supplemental Figure 3.**
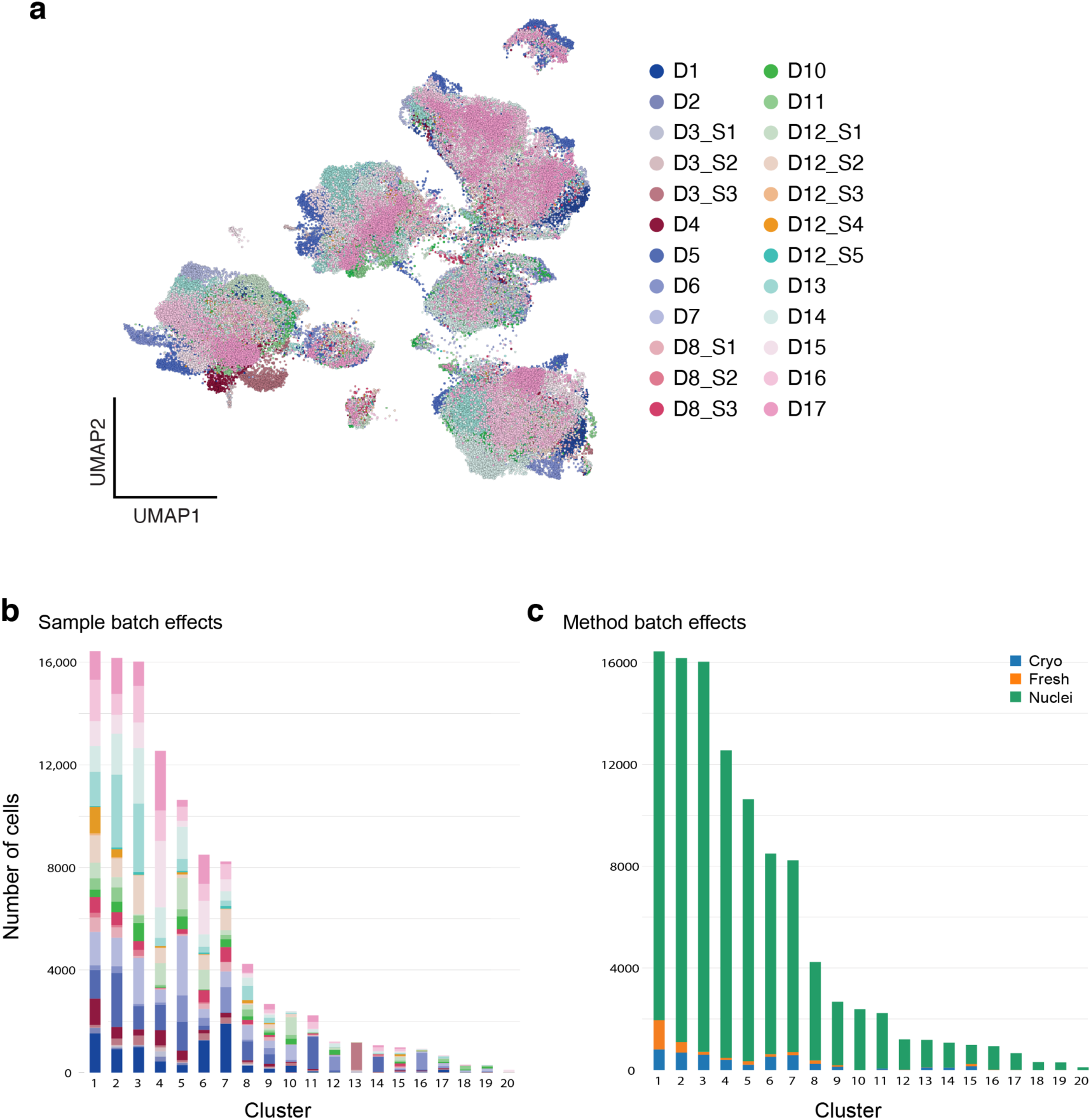
Cross-sample integration corrects batch effects of the COVID-19 lung cell atlas. **a.** UMAP (as in Fig. 2a) of 106,792 sc/snRNA-Seq profiles post-Harmony^34^ correction (**Methods**) colored by sample ID. **b,c.** Donors and processing protocols are represented across clusters. Number of cells (*y* axis) from different donors (**b**) or processing protocols (**c**) in each Leiden cluster (*x* axis).

**Supplemental Figure 4.**
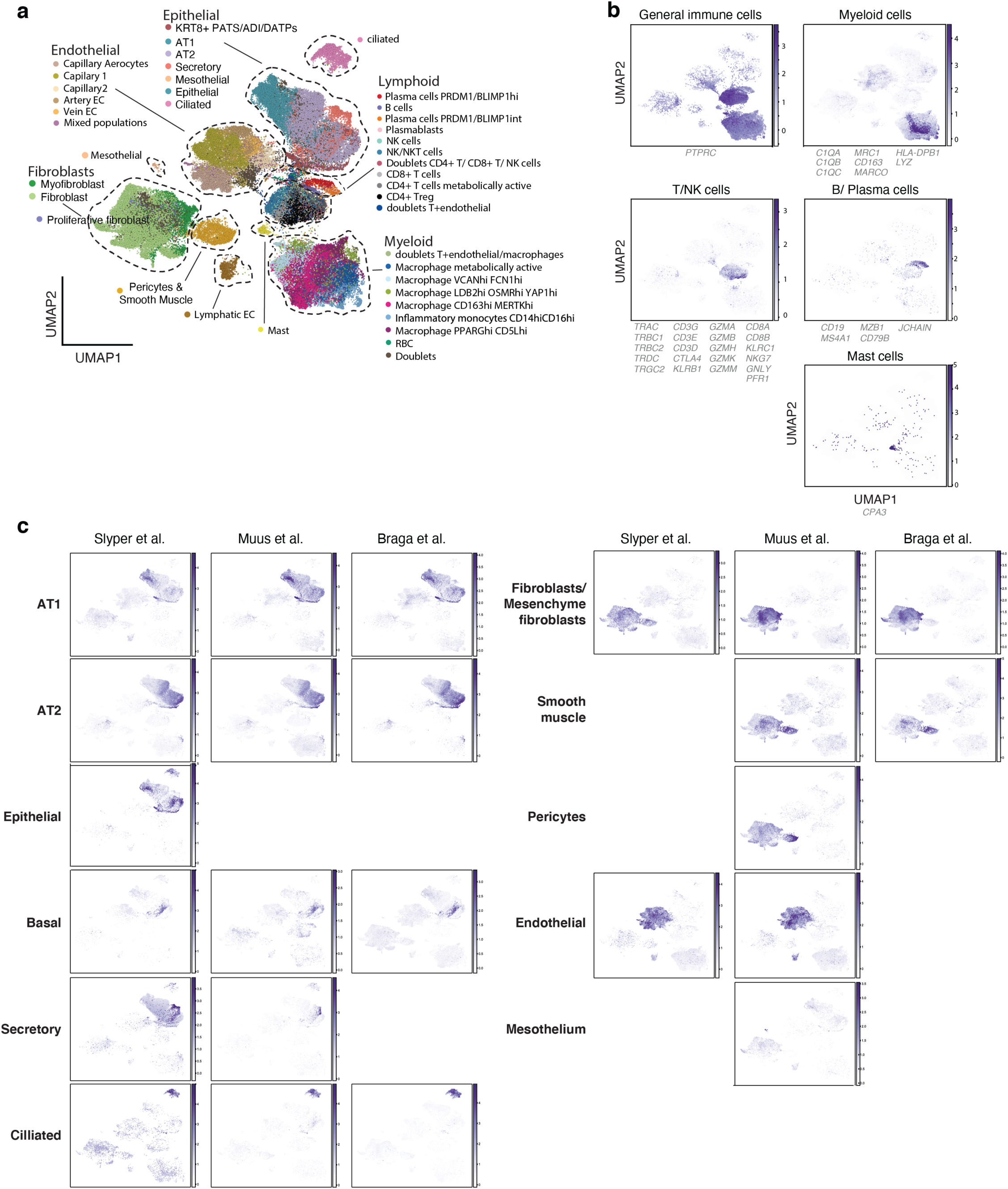
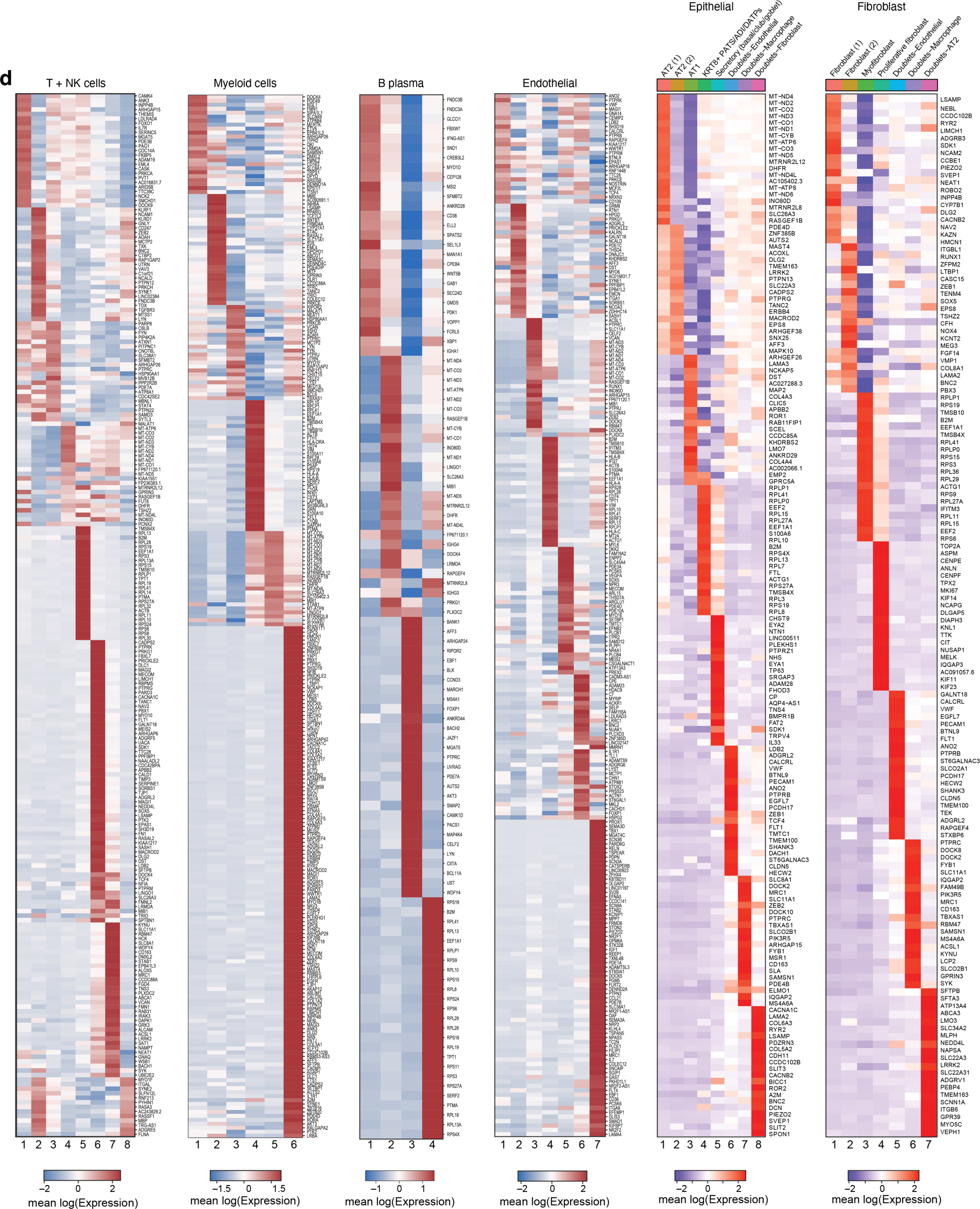
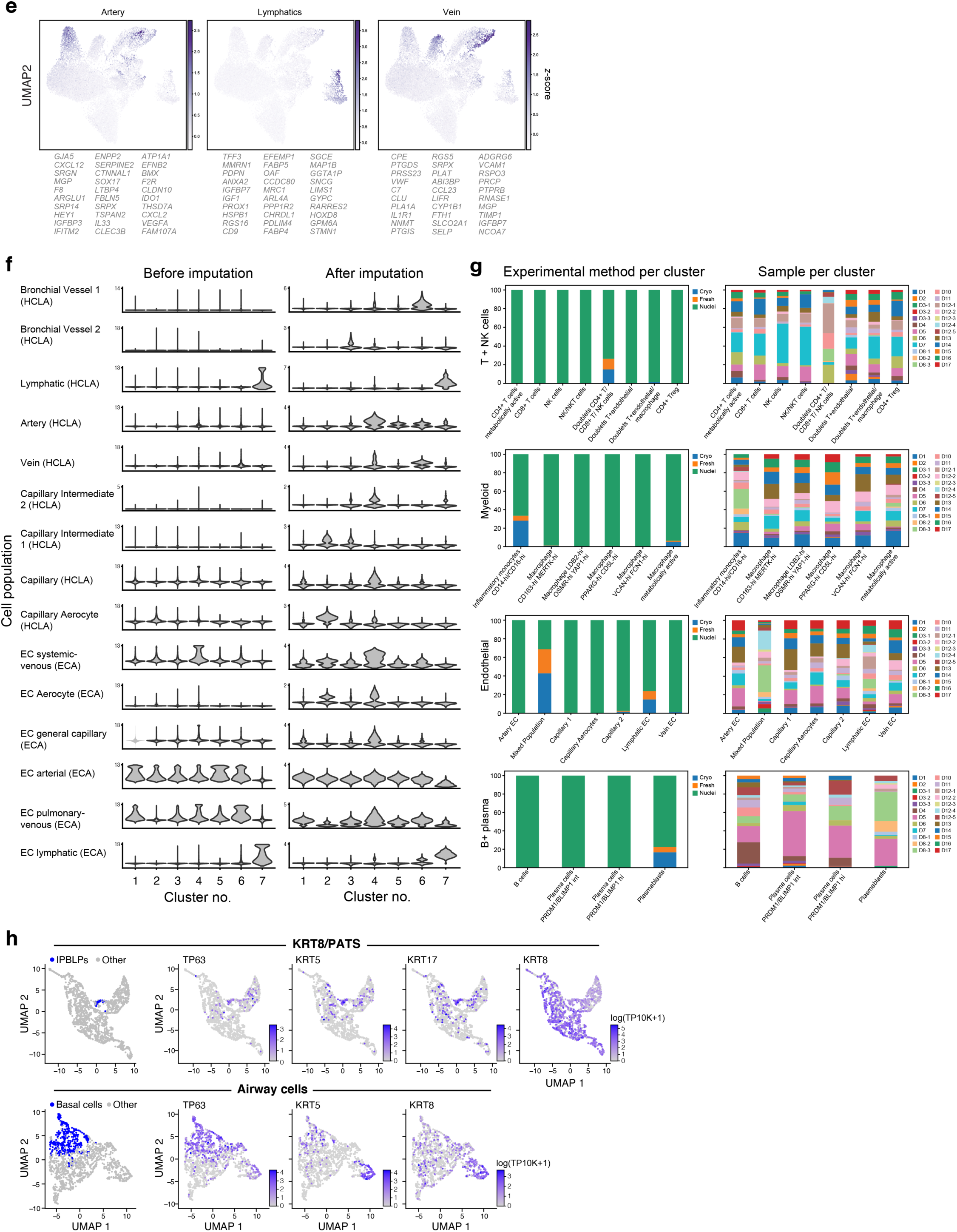
Manual annotation in the COVID-19 lung cell atlas. **a-c.** Identification of main lineage annotations. UMAP of 106,792 sc/snRNA-Seq profiles post-Harmony^34^ correction (as in Fig. 2a) colored either by manual annotation done in sub-clustering of each lineage (**a**), by expression of genes used to separate immune cell sub-lineages (**b**), or by signatures from published compartment signatures (**Supplemental Table 17**) used to identify other lineages (**c.**). Dashed lines in **a**: chosen compartments for sub-clustering. **d.** Differentially expressed genes between sub-clusters within each lineage. Expression (color bar) of genes (rows) that are differentially expressed (**Methods**) across the sub-clusters (columns) within each compartment (label on top). **e.** Endothelial cell annotations for sub-clusters 5,6,7. UMAP (as in Fig. 2e) of endothelial genes with the Z-scores of published endothelial cell signatures (top 30 differentially upregulated genes in each cell type from a lung cell atlas^35^). **f.** Imputation allows retrieval of lost signal in endothelial cells and identifying endothelial subsets defined in prior studies. ECA - endothelial cell atlas markers Schupp *et al*.^36^, HCLA - human cell lung atlas markers Gillich *et al*.^108^ **g.** Batch correction within lineage. Fractional abundance (*y* axis) from different processing protocols (left) or different donors (right) in each sub-cluster (*x* axis) after batch correction with Harmony^34^ within each lineage. **h**. UMAP embeddings of KRT8^+^ PATS-expressing cells (top) or of airway epithelial cells (bottom) colored by basal cells (blue, leftmost panels) or characteristic markers (purple, remaining panels).

**Supplemental Figure 5.**
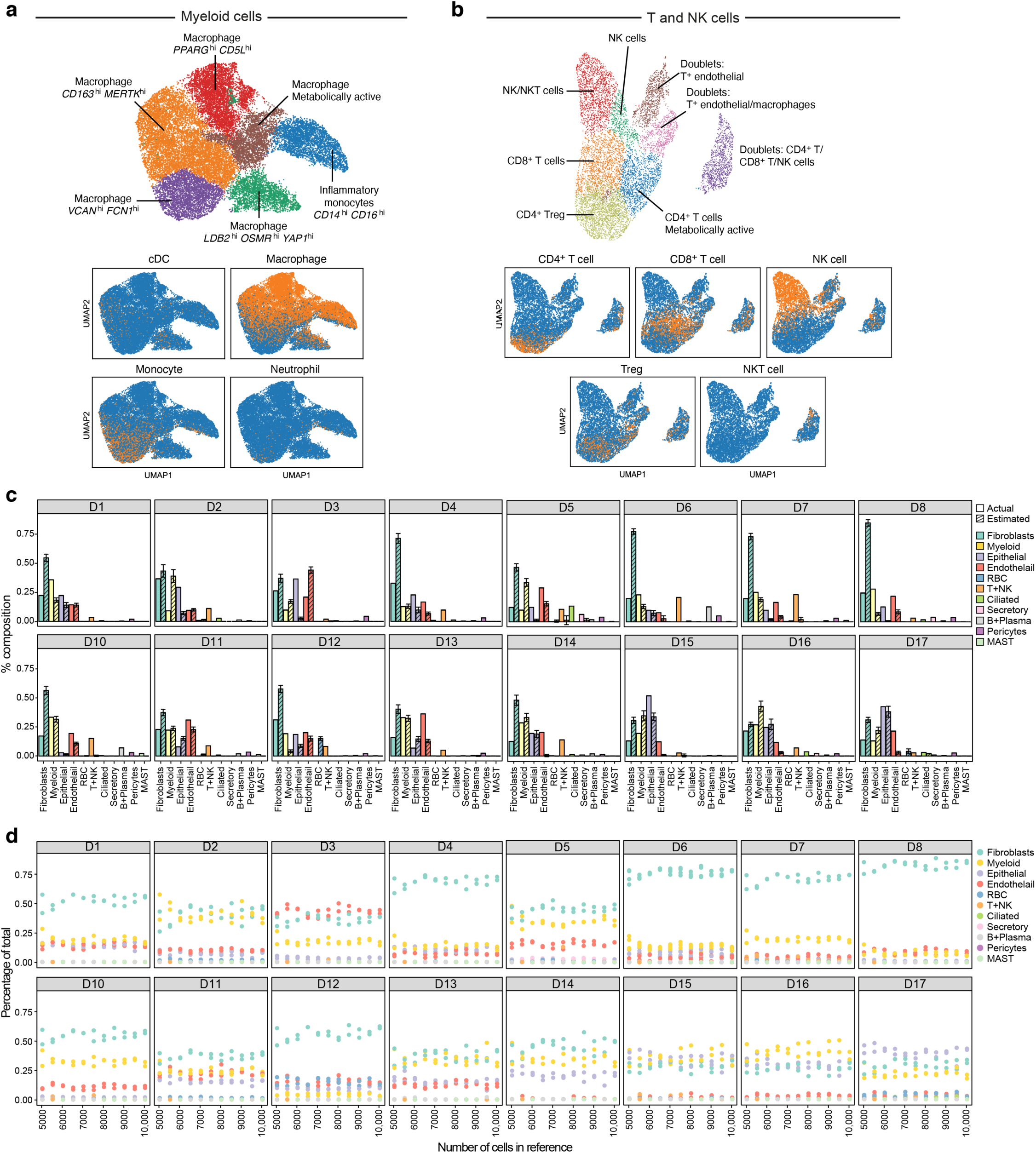
Manual annotation in the COVID-19 lung cell atlas and bulk RNA-Seq deconvolution. **a-b.** Visualization of the comparison between the automated predictions (bottom) and the manual annotations (top) of select subsets of myeloid cells **(a**) and T and NK cells (**b**). **c.-e.** Results from deconvolution of bulk RNA-Seq libraries from adjacent lung tissue. **c.** Mean proportion (*y* axis, from MuSiC^93^) estimates for each of 11 cell subsets (*x* axis) in each of 16 bulk RNA-Seq lung samples (panels) from 10 random samples of 10,000 cells each. **d.** Robustness of cell proportion estimates to the number of single cells sampled for the reference data. Mean proportion (*y* axis, from MuSiC) estimates for each of 11 cell subsets (color dots) in each of 16 bulk RNA-Seq lung samples (panels) when using three independent samples of 1,000 to 10,000 cells from the single cell reference (*x* axis). **e.** Proportions of single-cell types in the reference annotation. Fibroblasts, myeloid, epithelial, endothelial, and T+NK cells make up the bulk of the annotated cells in the sc/snRNA-Seq data.

**Supplemental Figure 6.**
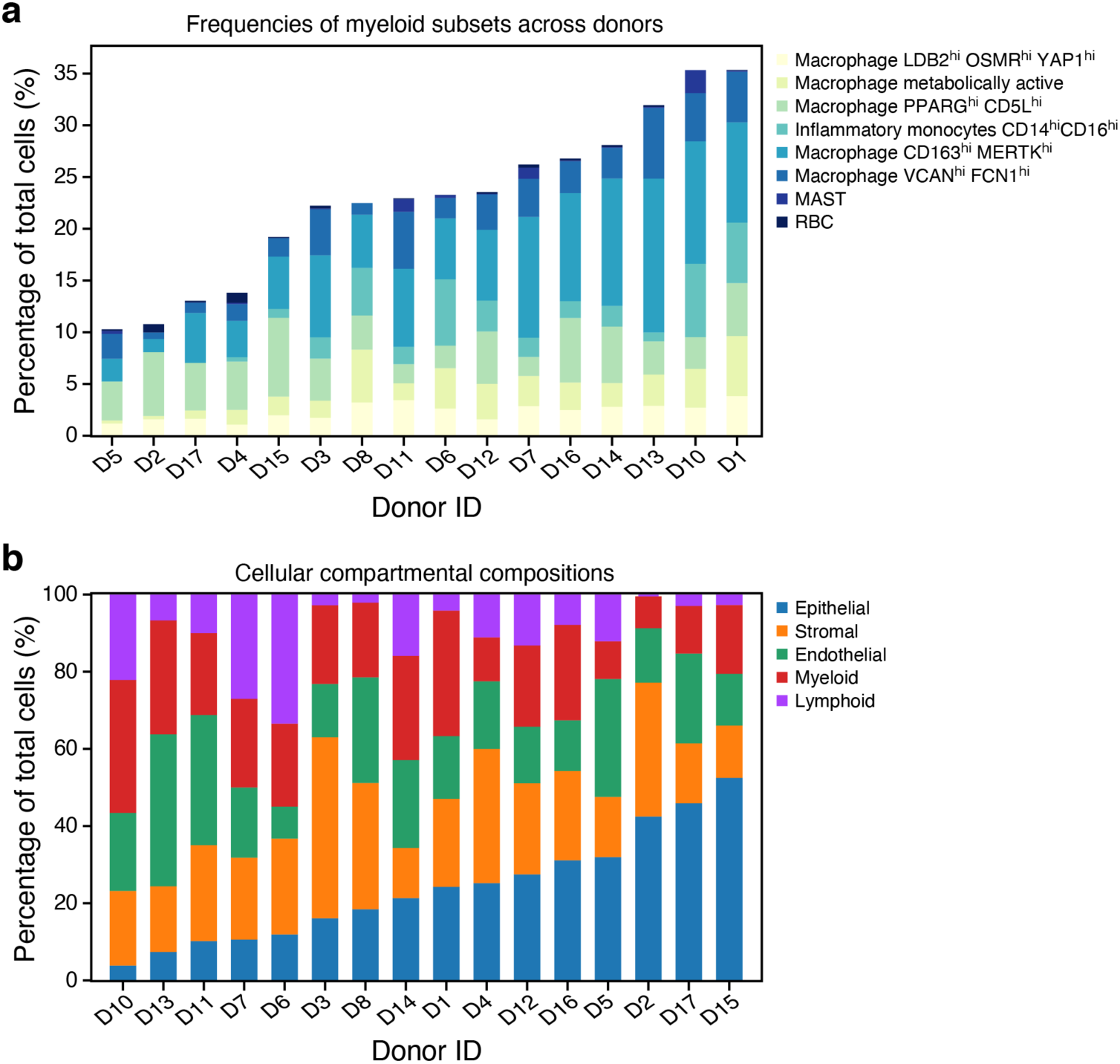
Donor heterogeneity of subtype and lineage composition. **a.** Percent composition of myeloid subsets per donor. **b.** Percent composition of main lineages by donor, sorted by the epithelial lineage.

**Supplemental Figure 7.**
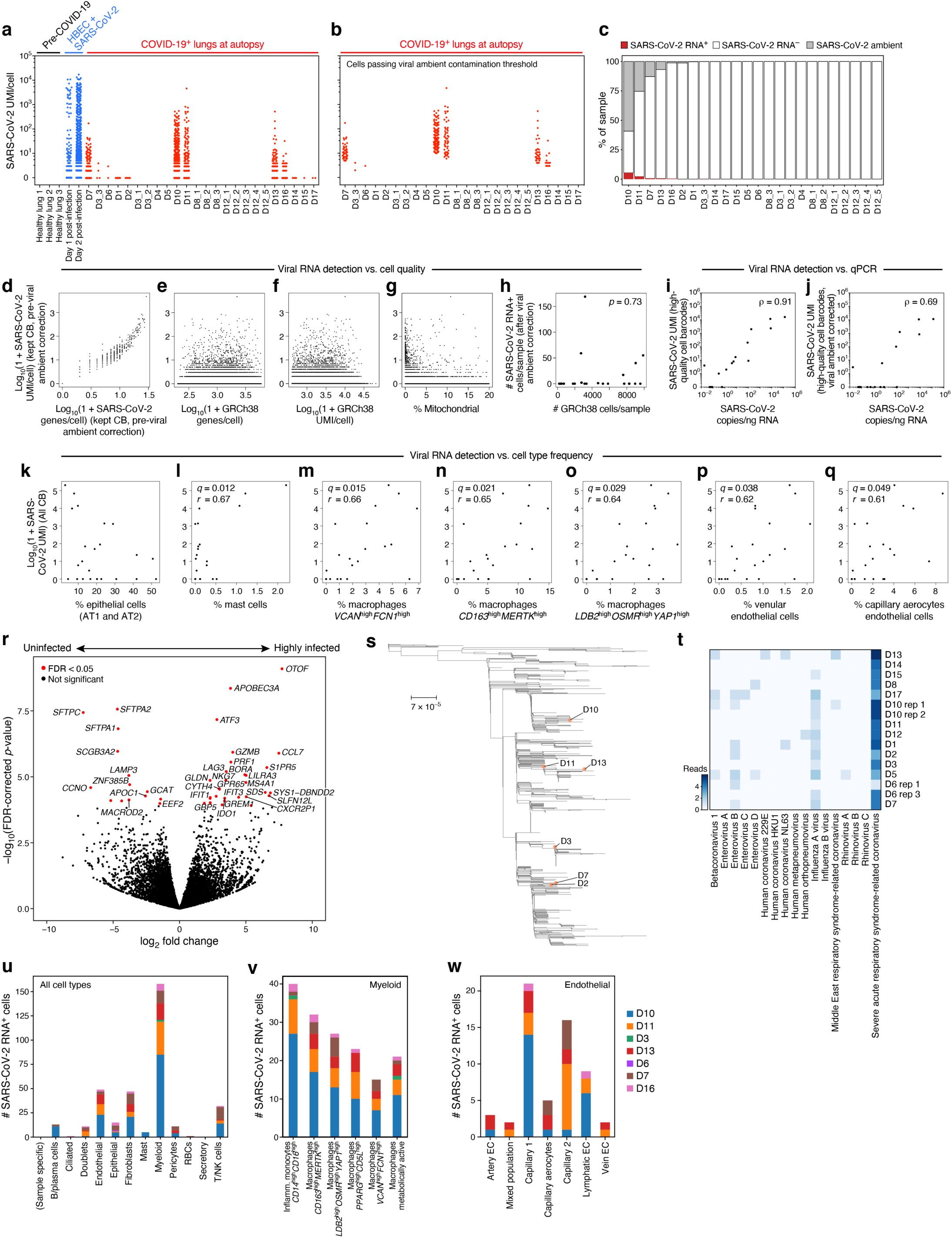
SARS-CoV-2-RNA+ cells distinguished by sc/snRNA-Seq. **a,b.** Impact of ambient RNA on SARS-CoV-2 RNA+ detection. Number of SARS-CoV-2 aligning UMI per Cell Barcode (CB) (*y* axis) in healthy lung (**a**, black), *in vitro* SARS-CoV-2 infected human bronchial epithelial cells (HBEC)^109^ (**a**, blue) or lung samples from COVID-19 donors at autopsy with CB with high-quality capture of human mRNA (red, **a**) or after removal of cells whose viral alignments were attributed to ambient contamination (red, **b**, **Methods**). **c.** Variation in SARS-CoV-RNA+ cells across donors. Percent of cells (*y* axis) assigned as SARS-CoV-2 RNA^-^ (white), SARS-CoV-2 RNA^+^ (red), or SARS-CoV-2 ambient (grey, **Methods**) across the donors (*x* axis), sorted by proportion of SARS-CoV-RNA+ cells. **d-h.** Viral RNA detection does not correlate with cell quality metrics. **d-g.** Number of SARS-CoV-2 UMIs (prior to ambient viral correction) for each cell (*y* axis) and either number of SARS-CoV-2 genes for that cell (**d**, *x* axis), number of human (GRCh38) genes per cell (**e**, *x* axis), number of human (GRCh38) UMI per cell (**f**, *x* axis), or % of human (GRCh38) mitochondrial UMIs per cell (**g**, *x* axis). **h.** Number of retained high-quality cells (*x* axis) and number of SARS-CoV-2 RNA+ cells (*y* axis) in each sample (dots) following correction for ambient viral reads. Pearson’s r = 0.07, p = 0.7, not significant. **i,j.** Agreement in viral RNA detection between qPCR and snRNA-Seq. Number of SARS-CoV-2 copies measured by CDC N1 qPCR on bulk RNA extracted from matched tissue samples (*x* axis) and the number of SARS-CoV-2 aligning UMI (*y* axis) for each sample (dot) from all reads (**i**) and after viral ambient RNA correction (**j**). Spearman’s ⍴ reported, p < 0.01 for each. **k-q.** Relation between SARS-CoV-2 RNA and different cell types. Number of SARS-CoV-2 aligning UMIs in each (including all CB) and the proportion of epithelial (AT1 and AT2) (**k**), mast (**l**), macrophage *VCAN*^high^*FCN1*^high^ (**m**), macrophages *CD163*^high^*MERTK*^high^ (**n**), macrophages *LDB2*^high^*OSMR*^high^*YAP*1^high^ (**o**), venular endothelial (**p**) or capillary aerocytes (**q**) cells in these samples (*x* axes). Pearson’s r denoted in the upper left corner with significance following Bonferroni correction (q). **r.** Impact of viral load on bulk RNA profiles. Significance (-log_10_(P-value), *y* axis) and magnitude (log_2_(fold-change), *x* axis) of differential expression of each gene (dots) between 3 donors with highest viral load and 6 donors with lowest/undetectable viral load profiled by bulk RNA-Seq. Red points: FDR < 0.05. **s.** Genetic diversity of SARS-CoV-2. Maximum likelihood phylogenetic tree of 772 SARS-CoV-2 genomes from cases in Massachusetts between January-May 2020. Orange points: donors in this cohort. **t.** Specificity of SARS-CoV-2 infection. log_10_(1+reads) in each donor (columns) assigned to different viruses (rows) by metagenomic classification using Kraken2 from bulk RNA-Seq. **u-w.** Distribution of SARS-CoV-2 RNA+ cells across cell types and subsets. Number of SARS-CoV-2 RNA+ cells (*y* axis) from each donor (color) across major categories (**u**, *x* axis), within myeloid subsets (**v**, *x* axis), or within endothelial subsets (**w**, *x* axis).

**Supplemental Figure 8.**
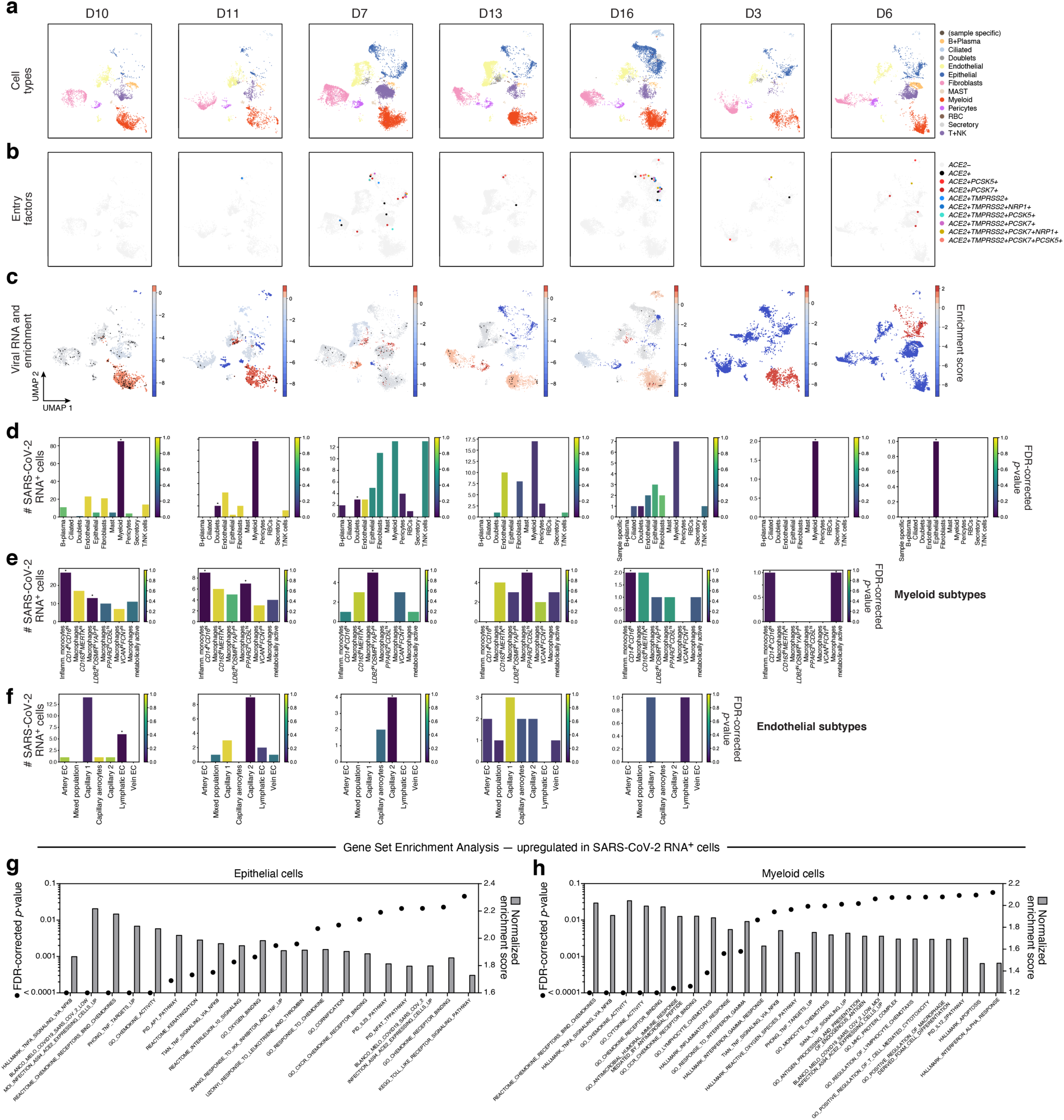
Donor-specific enrichment of SARS-CoV-2 RNA+ cells and host responses to viral RNA. **a-c**. UMAP embedding of sc/snRNA-Seq profiles from each of 7 donors with SARS-CoV-2 RNA+ cell barcodes (columns), colored by major cell categories (**a**), expression of SARS-CoV-2 entry factors (**b.**, as in Fig. 4g), or SARS-CoV-2 RNA enrichment per cluster (**c.**, red/blue colorbar; red: high enrichment; black points: SARS-CoV-2 RNA+ cells). **d-f.** Number of SARS-CoV-2 RNA^+^ cells (*y* axis) across major cell type (**d**, *x* axis) or within myeloid (**e**, *x* axis) or endothelial (**f**, *x* axis) cell subsets. Bars are colored by enrichment score (dark blue: stronger enrichment). *FDR < 0.05. **g,h.** Normalized enrichment score (bars, right *y* axis) and significance (points, FDR, left *y* axis) (by GSEA^53, 54^, **Methods**) of different functional gene sets (*x* axis) in genes upregulated in SARS-CoV-2 RNA+ epithelial (**g**) or myeloid (**h**) cells *vs.* bystanders.

**Supplemental Figure 9.**
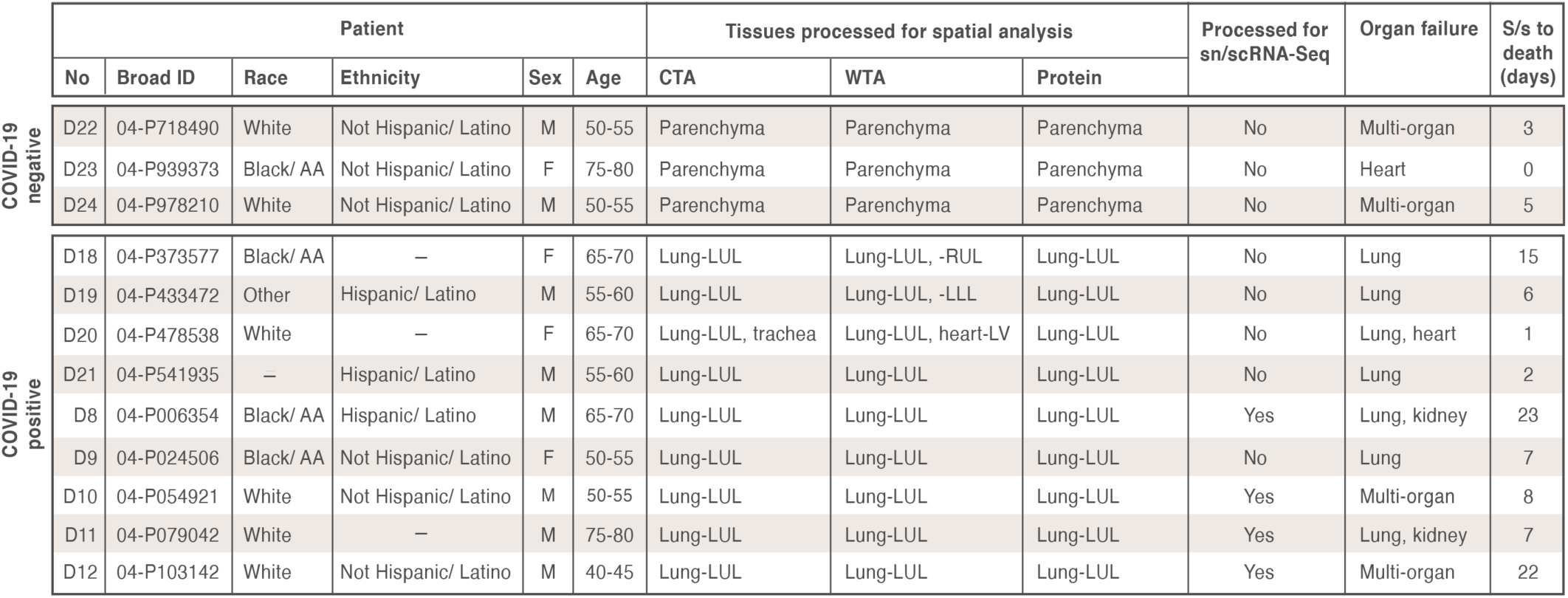
Metadata for the COVID-19 autopsy lung spatial atlas. Metadata for FFPE tissues from 12 donors (3 SARS-CoV-2 negative, 9 SARS-CoV-2 positive). S/s to death: time from onset to death, in days.

**Supplemental Figure 10.**
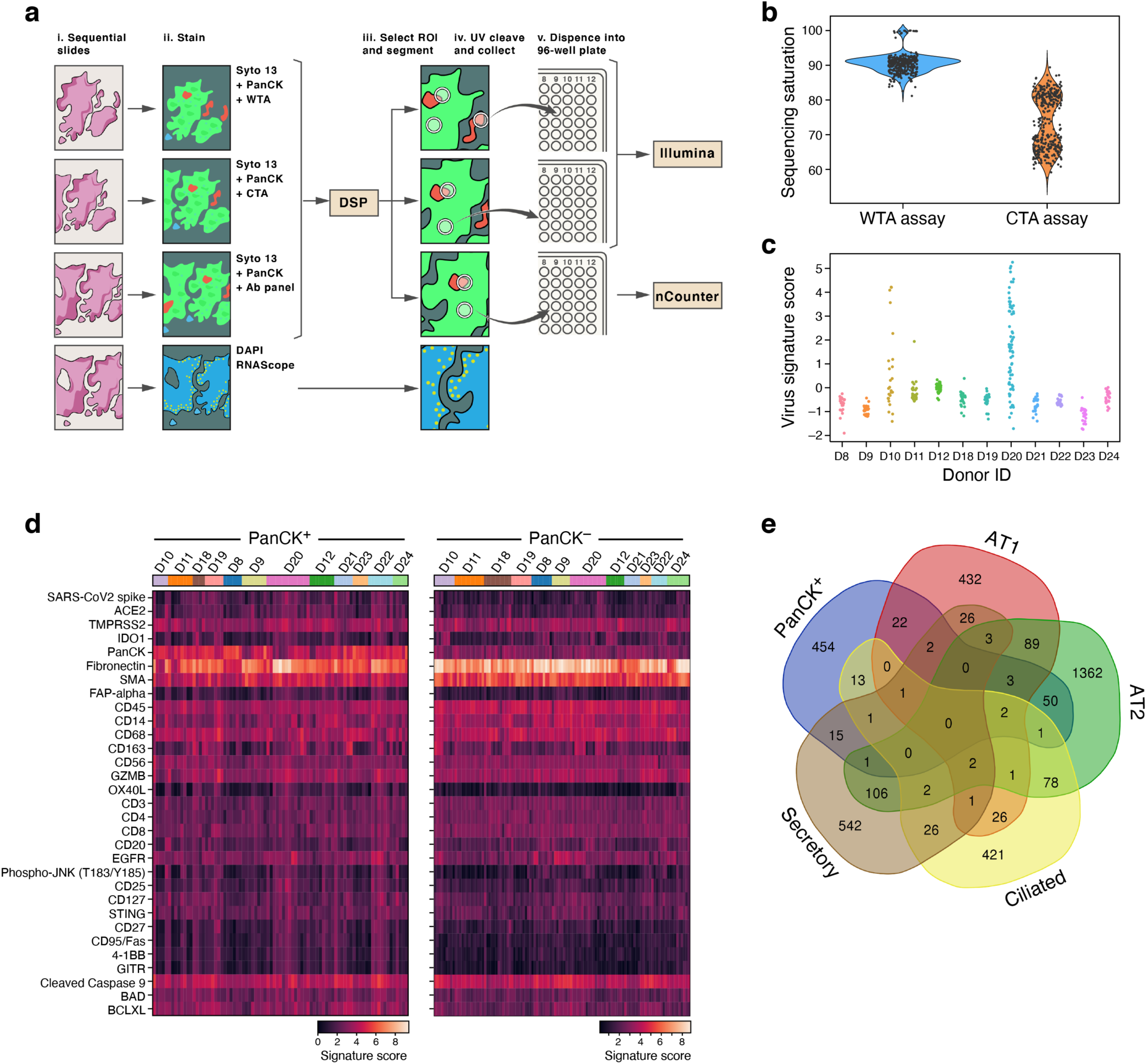
NanoString GeoMx experiment design and analysis. **a.** Overview of spatial profiling experiments. **b.** Sequencing saturation. Distribution of sequencing saturation (*y* axis, %) for AOIs from WTA and CTA AOIs (*x* axis). **c.** Variation in viral load across AOIs and between donors. Viral signatures score (*y* axis) for each CTA AOI (dots) from each donor (*x* axis). **d.** Expression (log(normalized intensity + 1), normalized by geometric mean of 3 housekeeping genes: Histone *H3*, *S6*, *GAPDH*) of 31 of 77 measured proteins (rows) in PanCK^+^ (left) and PanCK^-^ (right) alveoli AOIs (columns) across donors (horizontal color bar). Donors D22-24 are SARS-CoV-2 negative while the rest are SARS-CoV-2 positive. Measured protein levels were heterogeneous across AOIs, without clear donor-specific effects, but the PanCK^-^ compartment was enriched for immune protein. **e.** Relation between PanCK^+^ alveolar expression and different cell type programs. Venn diagram of 565 upregulated genes in the PanCK^+^ alveolar compartment of SARS-CoV-2 positive samples *vs.* SARS-CoV-2 negative samples and cell type markers from Muus et al. Supplementary Table 6 of different *ACE2*-high epithelial cells (FDR < 0.05).

**Supplemental Figure 11.**
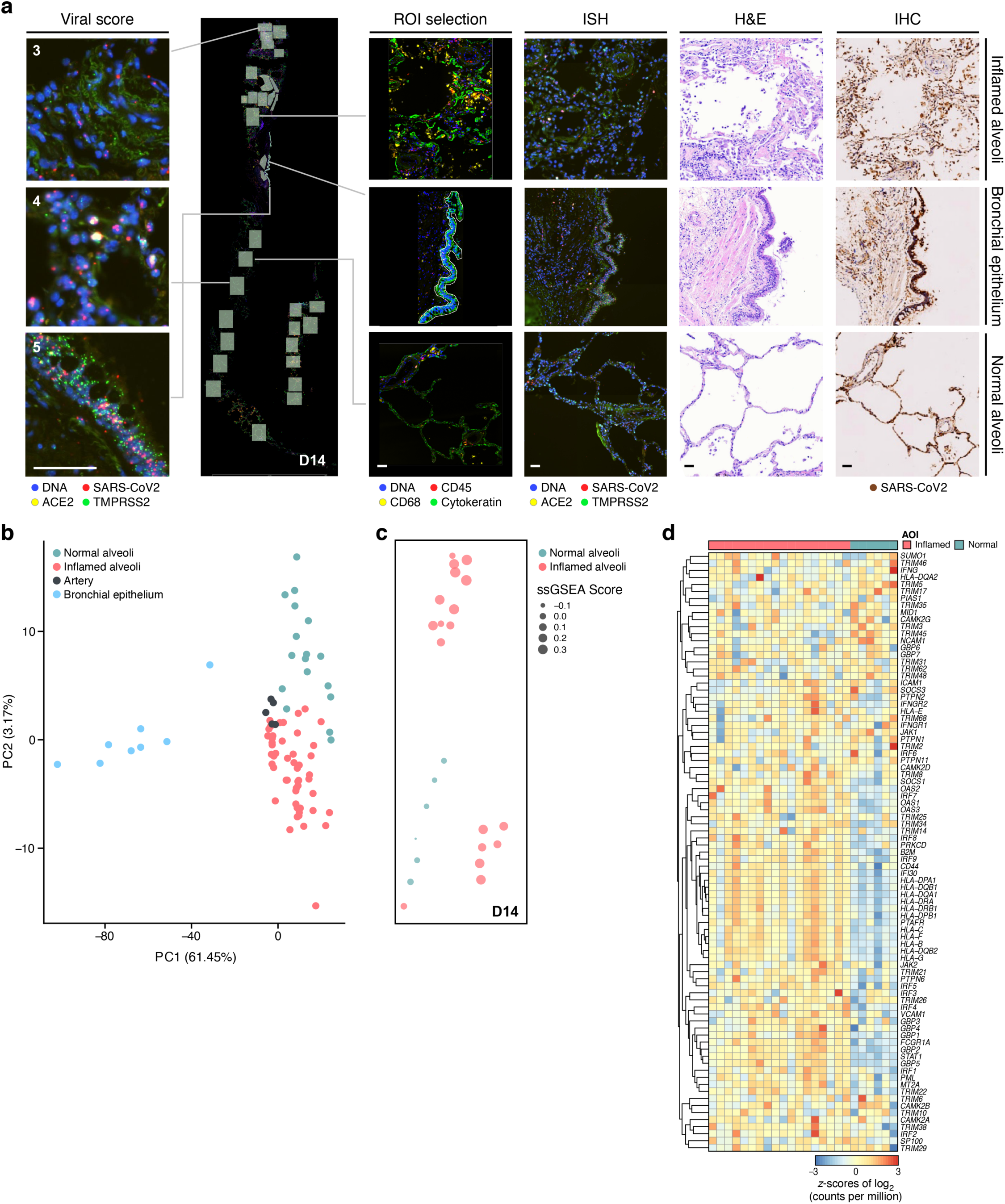
GeoMx WTA DSP analysis of lung biopsies reveals region- and inflammation-specific expression programs. **a.** Region selection. Serial sections of lung biopsies (D13-17, plot depicting serial sections of D14) processed with GeoMx WTA-DSP with 4-color staining (DNA, *CD45*, *CD68,* PanCK), RNAscope with probes against (SARS-CoV-2 *S*-gene (utilized to derive semi-quantitative viral load scores), *ACE2*, *TMPRSS2*), H&E staining, and immunohistochemistry with anti-SARS-CoV-2 S-protein. Scale bar: 100 µm. **b-d**. Regions and inflammation specific expression programs. **b.** The first two principal components (PCs, *x* and *y* axis) from lung ROI gene expression profiles from donors D13-17 spanning normal-appearing alveoli (green), inflamed alveoli (magenta), bronchial epithelium (blue), and arterial blood vessels (black). **c.** GSEA score (circle size, legend) of the enrichment of the interferon-γ pathway in each normal (green) and inflamed (magenta) alveolar ROI (dot) from the section of donor D14 (in **a**), placed in their respective physical coordinates on the tissue section (as in **a**). **d.** Normalized expression (color bar) of the interferon-γ pathway genes (rows) from normal (green) and inflamed alveoli (magenta) ROIs (columns) from D14 lung biopsy.

**Supplemental Figure 12.**
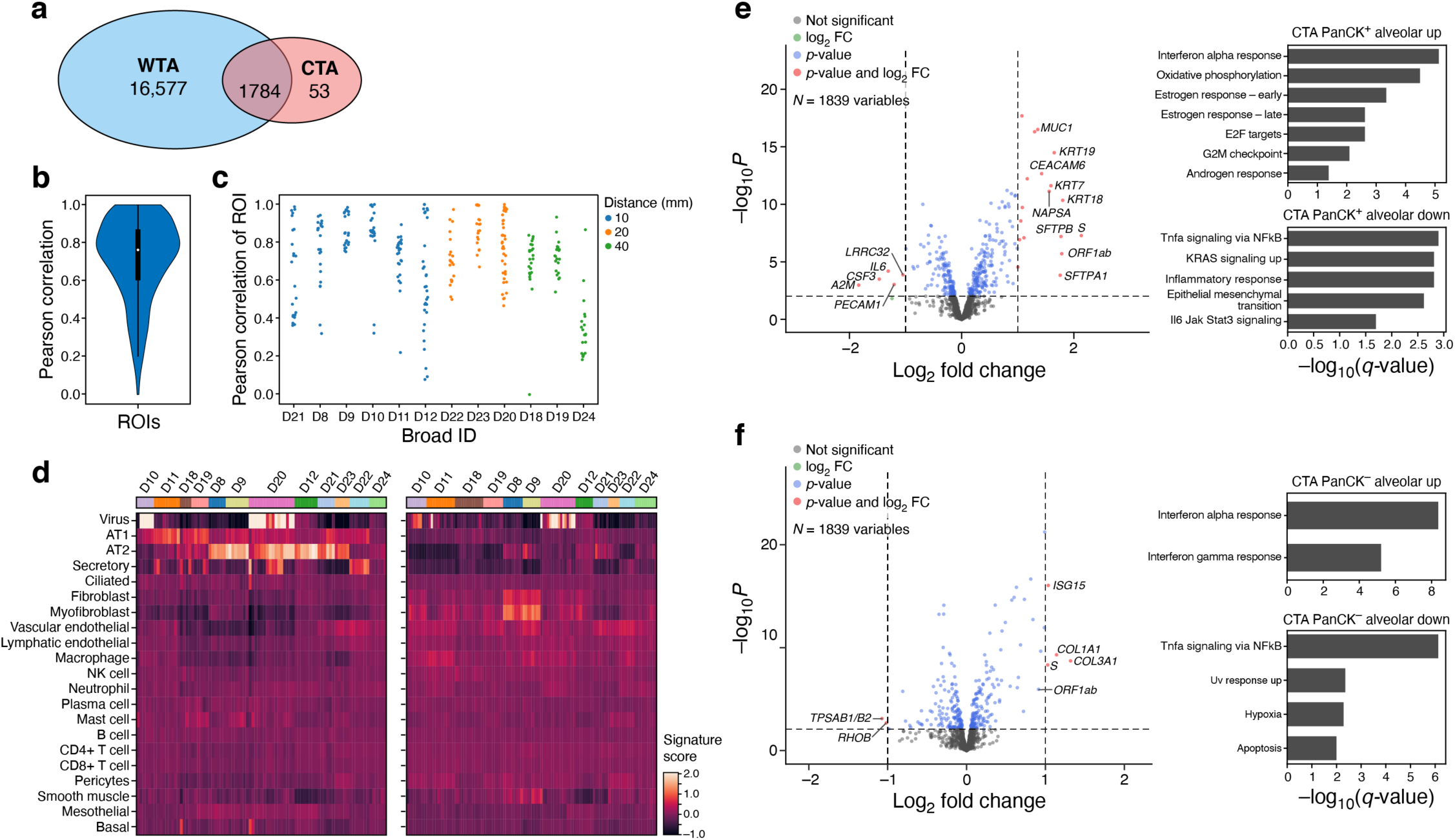
CTA characterization of lung autopsy samples. **a.** Overlap of WTA and CTA genes, including the 26 supplemented SARS-CoV-2 biology-associated genes. **b,c.** Agreement between WTA and CTA. **b.** Distribution of Pearson correlation coefficients (*y* axis) between WTA and CTA profiles (for common genes) across 206 ROIs from paired sample locations. **c.** Pearson correlation coefficient (*y* axis) of WTA and CTA common genes for each AOI pair (dot) in each donor (*x* axis), sorted by distance between WTA and CTA sections. **d**. Differences in cell type composition between PanCK^+^ and PanCK^-^ alveolar AOIs. Expression scores (color bar) in CTA data for different cell type signatures (rows) in the PanCK^+^ (left) and PanCK^-^ (right) alveolar AOIs (columns) across donors (horizontal color bar). Donors D22-24 are SARS-CoV-2 negative while the rest are SARS-CoV-2 positive. **e,f.** Changes in gene expression in COVID-19 *vs*. healthy alveolar AOIs in CTA data. Left: Significance (-Log_10_(P-value), *y* axis) and magnitude (log_2_(fold-change), *x* axis) of differential expression of each gene (dots) in CTA data between COVID-19 and healthy AOIs for PanCK^+^ (**e**) and PanCK^-^ (**f**) alveoli. PanCK^+^ alveoli ROIs: 69 SARS-CoV-2 positive *vs.* 18 negative; PanCK^-^ alveoli ROIs: 92 SARS-CoV-2 positive *vs.* 22 negative. The horizontal dashed line indicates an FDR q-value cutoff of 0.05, and the two vertical dashed lines represent a fold-change of 2 in log_2_ scale. The names of the 10 genes with the highest, statistically significant fold change between SARS-CoV-2 positive and negative are marked. Right: Significance (-log_10_(q-value)) of enrichment (permutation test) of different pathways (rows).

**Supplemental Figure 13.**
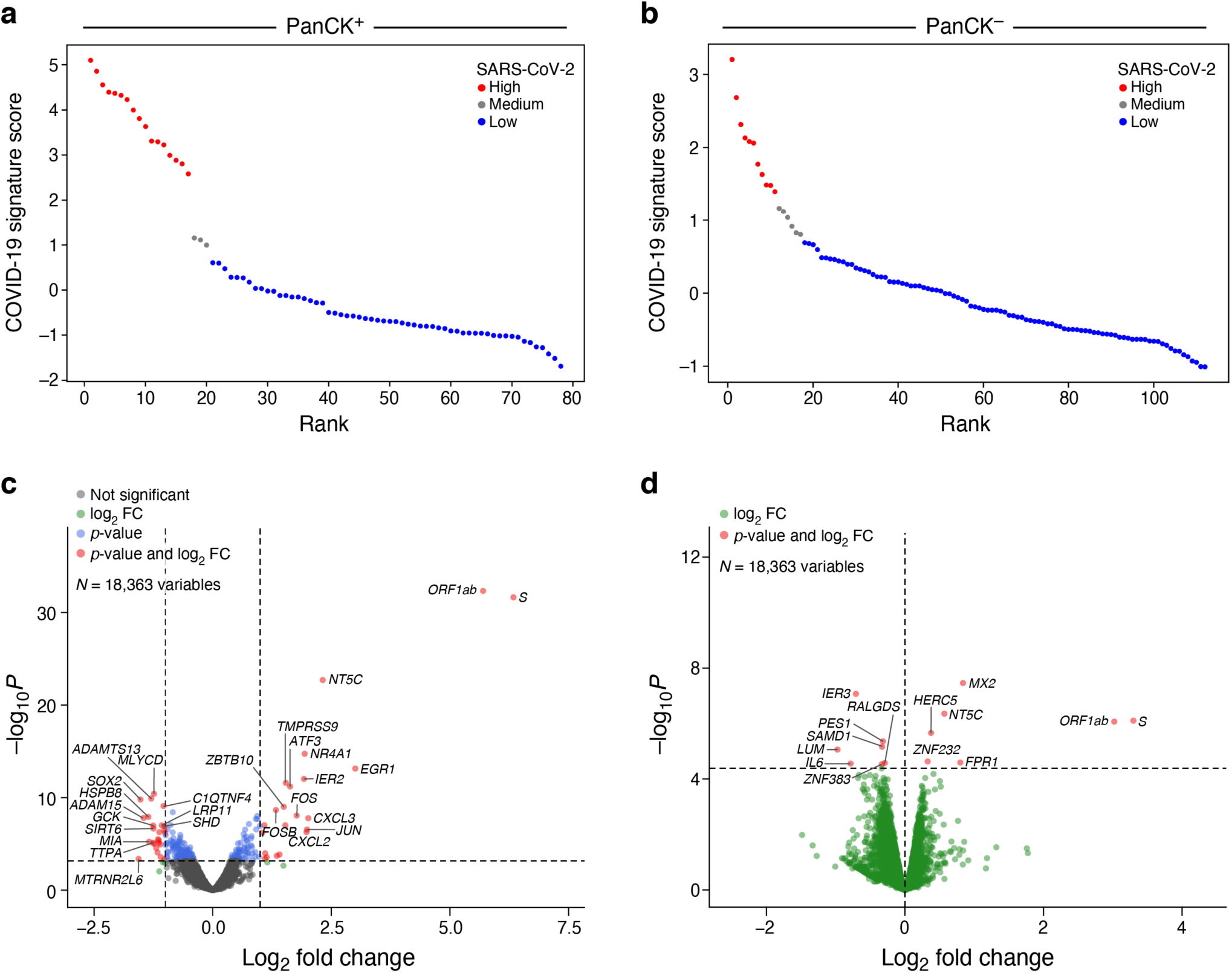
Differential expression between SARS-CoV-2 high and SARS-CoV-2 low compartments in alveolar AOIs. **a,b.** SARS-CoV-2 high and SARS-CoV-2 low alveolar AOIs. PanCK^+^ (**a**) or PanCK^-^ (**b**) alveolar AOIs (dots) rank ordered by their SARS-CoV-2 signature score (*y* axis) in WTA data, and partitioned to high (red), medium (grey) and low (blue) SARS-CoV-2 AOIs. **c,d.** Changes in gene expression in SARS-CoV-2 high *vs.* low AOIs in WTA data. Significance (-Log_10_(P-value), *y* axis) and magnitude (log_2_(fold-change), *x* axis) of differential expression of each gene (dots) in WTA data between SARS-CoV-2 high and low AOIs for PanCK^+^ (**c**) and PanCK^-^ (**d**) alveoli. PanCK^+^ alveoli ROIs: 17 high, 3 medium, 58 low. PanCK^-^ alveoli ROIs: 11 high, 6 medium, 95 low. The horizontal dashed line indicates an FDR q-value cutoff of 0.05 in both (**c**) and (**d**). The two vertical dashed lines in (**c**) represent a fold-change of 2 in log_2_ scale, while there is no fold-change cutoff in (**d**). The names of top 10 SARS-CoV-2 high and SARS-CoV-2 low significant genes regarding fold-change are marked, respectively.

**Supplemental Figure 14.**
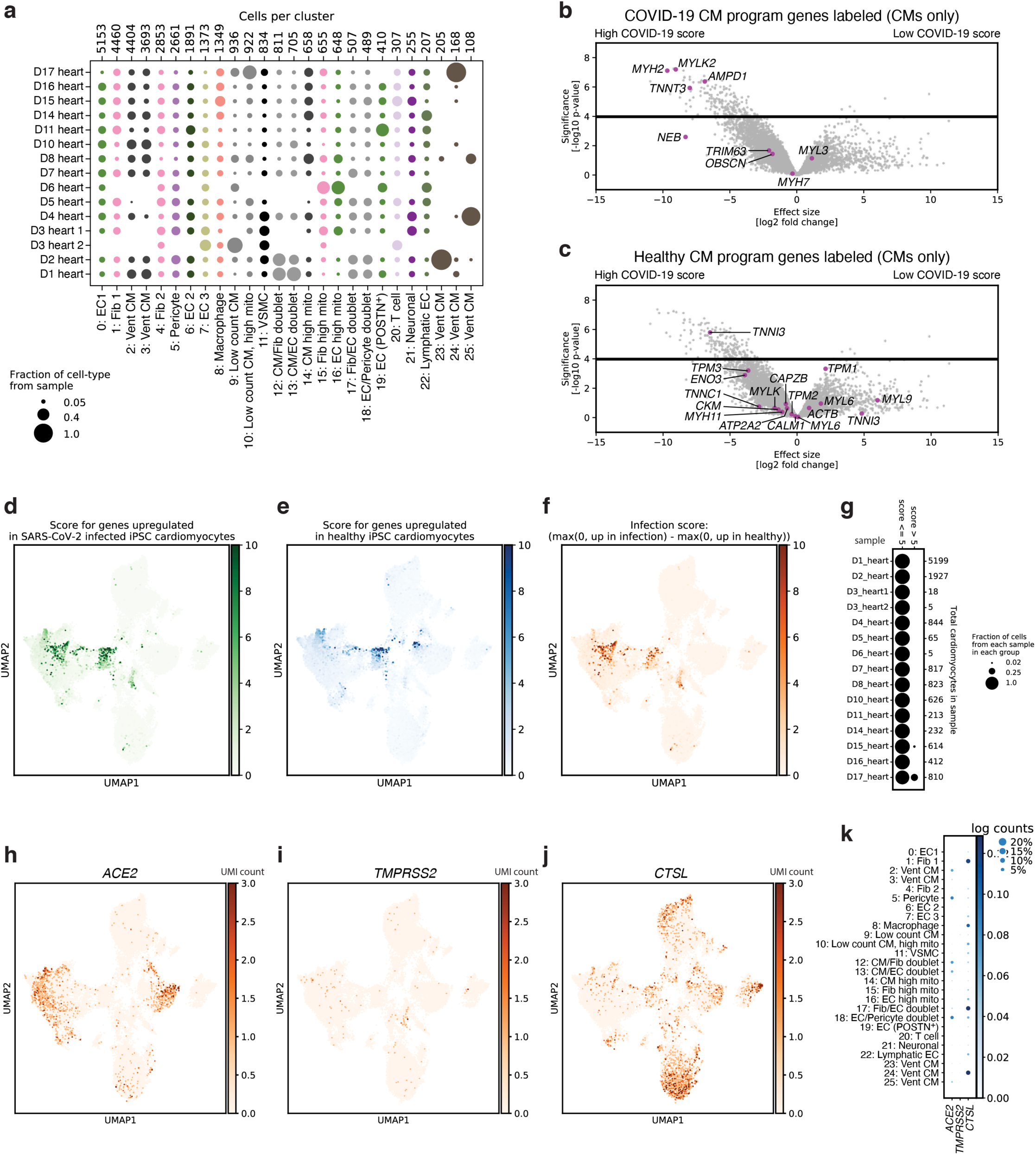
A COVID-19 heart cell atlas. **a.** Variation of cell type composition across donors. Proportion of cells (dot size) in each cluster (columns, labels from manual annotation as in Fig. 6a) derived from each sample (rows). Clusters 23-25 are largely sample-specific. **b-f.** Cardiomyocytes from one COVID-19 donor’s heart show signatures reported in *in vitro* infected iPSC derived cardiomyocytes. **b,c.** Significance (-Log_10_(P-value), *y* axis) and magnitude (log_2_(fold-change), *x* axis) of differential expression of each gene (dots) in cardiomyocytes (clusters 2, 3, 12-14, 23-25) between nuclei high-scoring for an *in vitro* infected cardiomyocyte signature (score > 5) vs. all others cardiomyocyte nuclei, colored for genes either upregulated (**b**) or downregulated (**c**) in SARS-CoV-2 infected *vs.* uninfected iPSC-derived cardiomyocytes^55^. The “score” as computed here correlates most strongly with upregulation of *MYH2*, *MYLK2*, *TNNT3*, *NEB*, and *AMPD1*. **d-f.** UMAP of COVID-19 heart snRNA-Seq profiles colored by the signature score (computed with Scanpy’s “score_genes” function, **Methods**) of genes upregulated (**d**) or downregulated (**e**) in genes in in SARS-CoV-2 infected *vs.* uninfected iPSC-derived cardiomyocytes^55^, or their difference (**f**, max(0, panel D score) -max(0, panel E score); also in Fig. 6b). **g.** Proportion of cardiomyocytes (dot size) in each sample (row) that score low (score<5) or high (score >5) (columns) for the infected iPSC-derived cardiomyocytes^55^ signature. Nearly all high scoring nuclei come from a single sample. **h-k**. Cell type specificity of SARS-CoV-2 viral entry factors. **h-j**. UMAP of COVID-19 heart snRNA-Seq profiles colored by expression of *ACE2* (**h**), *TMPRSS2* (**i**), and *CTSL* (**j**) (color is saturated to emphasize small counts). **k.** Mean expression (color) and proportion of expressing cells (dot size) for each viral entry factor (columns) across the cell subsets in the heart dataset (rows).

**Supplemental Figure 15.**
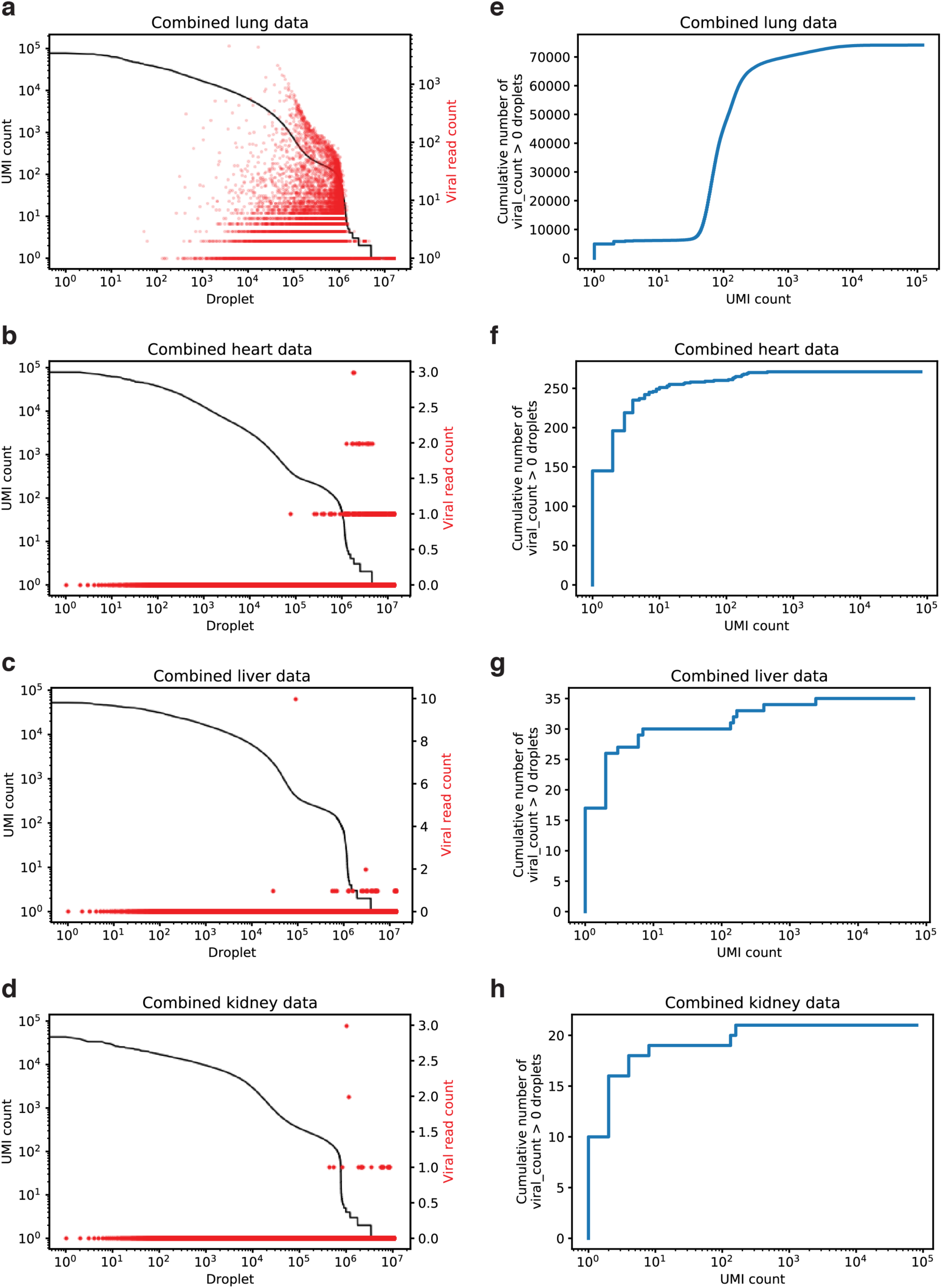
Different source of viral UMIs in heart, liver and kidney, compared to lung. **a, b, c, d.** Number of viral UMIs (red, right *y* axis) and all UMI (black, left *y* axis) for each droplet (*x* axis), rank ordered by total number of UMIs, in lung (**a**), heart (**b**), liver (**c**), and kidney (**d**). In lung (**a**) viral reads are in either ambient RNA (UMI counts (∼100 for many droplets) in the empty droplet plateau #200,000 - 1,000,000) or in nuclei, above and beyond the ambient RNA level. In heart (**b**), liver (**c**), and kidney (**d**) most viral UMI counts do not come from empty droplets, but mostly from droplets in the tail (beyond droplet 1,000,000), suggesting these are not ambient RNA, but a technical or mapping artifact. **e, f, g, h.** Cumulative distribution functions of the number of viral-positive droplets as a function of total droplet UMI count. In lung (**e**) most viral-positive droplets are empty droplets (total UMI count ∼ 100) with some viral-positive droplets which contain nuclei (UMI count > ∼1,000), but in heart (**f**), liver (**g**), and kidney (**h**), most of the “viral-positive” droplets have fewer than 10 total UMI counts, suggesting these reads are not trustworthy.

**Supplemental Figure 16.**
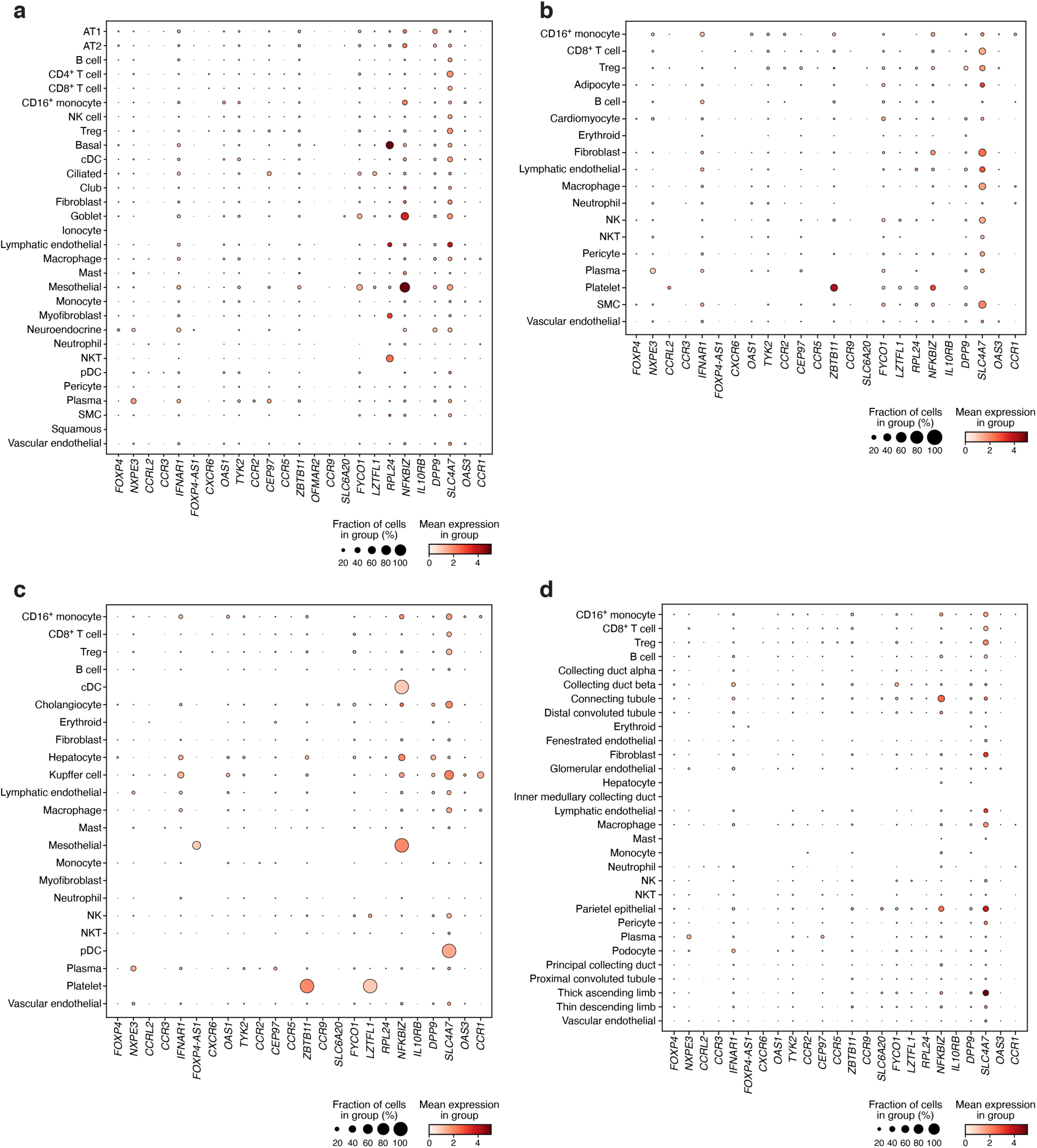
Expression of GWAS curated genes across lung, heart, liver and kidney atlases. Mean expression (dot color, log(TP10K + 1)) and proportion of expressing cells (dot size) for each curated GWAS gene (columns) in each cell subset (rows) for lung (**a**), heart (**b**), liver (**c**) and kidney (**d**) COVID-19 autopsy atlases. Some GWAS genes have higher expression in the lung compared to the other three tissues.

## Supplemental Tables and Files

**Supplemental Table 1. Clinical meta-data for all donors**

**Supplemental Table 2. Studies included in meta-atlas**

**Supplemental Table 3. Annotation classifier coefficients**

**Supplemental Table 4. Healthy vs. COVID-19 differentially expressed genes**

**Supplemental Table 5. Differentially expressed genes in cluster 7, epithelial cell sub-cluster**

**Supplemental Table 6. Cell-Type Specific Differentially Expressed Genes Between SARS-CoV-2 RNA+ vs. SARS-CoV-2 RNA-Cells.** This table contains 4 sheets, listing the DESeq2 output statistics following comparison of SARS-CoV-2 RNA+ cells *vs*. SARS-CoV-2 RNA-cells as listed in each header and sheet name. Only cell types with significant DE genes are listed.

**Supplemental Table 7. Differentially expressed genes between subject and control tissues of WTA data in alveolar AOIs.** Sheets of titles ending with “.up” give significant up-regulated genes on subject data, while those ending with “.down” give significant genes on control data. Only genes with adjusted p-value < 0.05 are listed; “PanCK” in sheet names refers to PanCK^+^ AOIs, while “Syto13” refers to PanCK^-^ AOIs.

**Supplemental Table 8. Differentially expressed genes between subject and control tissues of CTA data in alveolar AOIs.** Sheets of titles ending with “.up” give significant up-regulated genes on subject data, while those ending with “.down” give significant genes on control data. Only genes with adjusted p-value < 0.05 are listed; “PanCK” in sheet names refers to PanCK^+^ AOIs, while “Syto13” refers to PanCK^-^ AOIs.

**Supplemental Table 9. DE results of snRNA-Seq based on the reconciled annotation.** Each sheet lists up-regulated genes by comparing nuclei inside the corresponding cluster with those outside, under q-value control at 0.05 using Mann-Whitney U test. Genes are sorted by AUROC in descending order.

**Supplemental Table 10. Cell type markers used for cell type deconvolution of WTA and CTA data.** This table contains two sheets, one for WTA and one for CTA.

**Supplemental Table 11. List of up-regulated genes in patient PanCK^+^ alveolar AOIs overlapping with genes expressed at higher levels in epithelial cell types with high *ACE2* expression.** In each sheet title, the corresponding cell types are separated with spaces.

**Supplemental Table 12. SARS-CoV-2 high *vs.* SARS-CoV-2 low. Differentially expressed genes from subject tissues of WTA data in alveolar AOIs.** Sheets of titles ending with “.up” give significant up-regulated genes on SARS-CoV-2 high data, while those ending with “.down” give significant genes on SARS-CoV-2 low data. Only genes with adjusted p-value < 0.05 are listed; “PanCK” in sheet names refers to PanCK^+^ AOIs, while “Syto13” refers to PanCK^-^ AOIs.

**Supplemental Table 13. Gene descriptions for genes relevant to COVID-19 GWAS.** We list 26 genes (with gene descriptions) identified by the COVID-19 Human Genetics Initiative based on their proximity to GWAS peaks, and 64 genes identified by MAGMA analyses using 0kb and enhancer map S2G strategies.

**Supplemental Table 14. DE analysis of COVID-19 relevant genes for different program classes.** The Z-scores from DE analysis of cell type program (cells in a cell type *vs.* other cells) and DE analysis of disease progression program (cells in disease cell type *vs.* cells in healthy cell type) for the 25 curated GWAS genes and the 61 MAGMA implicated genes from **Supplemental Table 13**.

**Supplemental Table 15. Sc-linker heritability results for gene programs.** Enrichment and p-value of enrichment for annotations derived from cell type and disease progression programs in four different tissues (lung, liver, kidney and heart) for COVID-19 and severe COVID-19 phenotypes. The annotations are generated by combining the gene programs with the Roadmap∪ABC enhancer S2G strategy. Results only reported for programs that attain nominal significance in enrichment (p value < 0.05). All analyses are conditional on the 86 baseline-LD (v2.1) model annotations.

**Supplemental Table 16. Nominate functionally important genes in each cell type.** For each cell type, we report genes with high grade (> 0.8) in COVID-19/severe COVID-19 disease enriched gene program (in sc-linker analysis) that are additionally implicated by GWAS curated genes or MAGMA.

**Supplemental Table 17. Gene signatures used for high-level manual annotation.**

**Supplemental Table 18. List of differential gene expression between clusters of sub-clustering myeloid cells, as computed by Pegasus.**

**Supplemental Table 19. List of differential gene expression between clusters of sub-clustering T and NK cells, as computed by Pegasus.**

**Supplemental Table 20. List of differential gene expression between clusters of sub-clustering B and plasma cells, as computed by Pegasus.**

**Supplemental Table 21. List of differential gene expression between clusters of sub-clustering endothelial cells, as computed by Pegasus.**

**Supplemental Table 22. List of top genes for LIGER factors of epithelial cells.**

**Supplemental Table 23. List of top genes for LIGER factors of fibroblasts.**

